# Spatial metabolomics identifies localized chemical changes in heart tissue during chronic cardiac Chagas disease

**DOI:** 10.1101/2020.06.29.178038

**Authors:** Danya A. Dean, Gautham, Jair L. Siqueira-Neto, James H. McKerrow, Pieter C. Dorrestein, Laura-Isobel McCall

**Affiliations:** Department of Chemistry and Biochemistry, University of Oklahoma, Norman, Oklahoma, United States of America; Laboratories of Molecular Anthropology and Microbiome Research, University of Oklahoma, Norman, Oklahoma, United States of America; Department of Biology, University of Oklahoma, Norman, Oklahoma, United States of America; Skaggs School of Pharmacy and Pharmaceutical Sciences, University of California San Diego, La Jolla, California, United States of America; Center for Microbiome Innovation, University of California San Diego, La Jolla, California, United States of America; Collaborative Mass Spectrometry Innovation Center, University of California San Diego, La Jolla, California, United States of America; Department of Microbiology and Plant Biology, University of Oklahoma, Norman, Oklahoma, United States of America

**Author notes:** These authors contributed equally to this work (Danya A. Dean, Gautham).

## Abstract

Chagas disease (CD) is one of thirteen neglected tropical diseases caused by the parasite *Trypanosoma cruzi*. CD is a vector-borne disease transmitted by triatomines but CD can also be transmitted through blood transfusions, organ transplants and congenital transmission. While endemic to Latin America, *T. cruzi* infects 7-8 million people worldwide and can induce severe cardiac symptoms including apical aneurysms, thromboembolisms and arrhythmias during the chronic stage of CD. However, these cardiac clinical manifestations and CD disease pathogenesis are not fully understood. Using spatial metabolomics (chemical cartography), we sought to understand the localized impact of infection on the cardiac metabolome of mice chronically infected with two divergent *T. cruzi* strains. Our data showed chemical differences in localized cardiac regions upon chronic *T. cruzi* infection, indicating that parasite infection changes the host metabolome at select sites in chronic CD. These sites were distinct from the sites of highest parasite burden. In addition, we identified acylcarnitines and phosphocholines as discriminatory chemical families within each heart region, comparing infected and uninfected samples. Overall, our study indicated overall and positional metabolic differences common to infection with different *T. cruzi* strains, and identified select infection-modulated pathways. These results provide further insight into CD pathogenesis and demonstrate the advantage of a spatial perspective to understand infectious disease tropism.

**Author Summary:** Chagas disease (CD) is a tropical disease caused by the parasite *Trypanosoma cruzi*. CD originated in South America; however, there are now 7-8 million people infected worldwide due to population movements. CD is transmitted through a triatomine vector, organ transplants, blood transfusions and congenital transmission. It occurs in two stages, an acute stage (usually asymptomatic) and the chronic stage. Chronic stage CD presents with severe cardiac symptoms such as heart failure, localized aneurysms and cardiomyopathy. Unfortunately, what causes severe cardiac symptoms in some individuals in chronic CD is not fully understood. Therefore, we used liquid chromatography-tandem mass spectrometry to analyze the heart tissue of chronically *T. cruzi-*infected and uninfected mice, to understand the impact of infection on the tissue metabolome. We identified discriminatory small molecules related to *T. cruzi* infection. We also determined that regions with the highest parasite burden are distinct from the regions with the largest changes in overall metabolite profile; these locations of high metabolic perturbation provide a molecular mechanism to why localized cardiac symptoms occur in CD. Overall, our work gives insight to chronic cardiac CD symptom development and shapes a framework for novel treatment and biomarker development.

## Introduction

Chagas disease (CD) is a parasitic disease caused by the protozoan *Trypanosoma cruzi* and is one of the designated “neglected tropical diseases” [1]. *T. cruzi* is endemic to Latin America and infects 7-8 million people worldwide [1]. An estimated 300,000 infections have been recorded in the United States due to a large Latin American immigrant population and endemic transmission [2–4]. CD is primarily transmitted through triatomine insects of the *Triatoma* and *Rhodnius* genera [2]. Non-vectorial modes of transmission involve blood transfusion, transplacental transmission, and food and drink contaminated with *T. cruzi* [1]. The *T. cruzi* life cycle includes three main stages: epimastigotes, trypomastigotes and amastigotes. *T. cruzi* in the insect vector undergoes transformation from trypomastigotes to epimastigotes in the midgut, and then migrates to the hindgut and differentiates into infective trypomastigotes [1]. Upon triatomine defecation on the human host, the infective trypomastigotes enter the host through scratching or rubbing of the bite wound, or through eyes and mucosal surfaces [1]. Following mammalian host cell infection, trypomastigotes differentiate into amastigotes, which proliferate and subsequently transform into trypomastigotes [1]. CD has two disease stages: acute and chronic [1,2]. The acute stage is usually asymptomatic, or presents with non-specific symptoms (fever, malaise) [1,2]. 20-30% of infected individuals will then progressively develop clinical manifestations of chronic CD, including cardiomegaly, cardiac arrhythmias, apical aneurysms, megacolon, and megaesophagus [2]. *T. cruzi* infections are treated with either benznidazole or nifurtimox; however, these treatments cause significant adverse effects to the point that up to 30% of treated individuals fail to complete the full treatment course [5,6].

CD was previously considered to have an autoimmune etiology, but parasite persistence has now conclusively been demonstrated to be required for disease pathogenesis [7]. Along with parasite persistence, chronic pro-inflammatory responses, including cytokine release and CD8+ T cell-mediated cytotoxicity, contribute to tissue damage [8]. A heterogeneity of interacting parasite-host factors, including *T. cruzi* strain, load and tissue tropism, host genetic background, and mode of infection, influence the clinical outcomes of the disease [9,10]. However, CD disease pathogenesis is not yet completely understood [2]. A holistic understanding of the molecular pathways involved in disease progression could help identify new drug development avenues and outcome-predictive biomarkers.

Metabolites are the final products of mRNA and protein expression and protein activity, thus providing information closely linked to phenotype [11]. Metabolic pathways are druggable. They also change dynamically in response to disease [12,13]. As such, an improved understanding of metabolism in CD may lead to new avenues for drug development and CD patient monitoring. Acute *T. cruzi-*infection affects *in vitro* and *in vivo* host metabolic pathways, including decreasing mitochondrial oxidative phosphorylation-mediated ATP production [8,14–16]. In addition, acute *T. cruzi*-infected mice heart tissue and plasma showed significant up- or down-regulation of certain metabolic pathways, such as glucose metabolism (glucose levels elevated in heart tissue and lowered in plasma over time), tricarboxylic acid cycle (TCA) (decrease in select TCA metabolites in the heart tissue and a decrease in all TCA metabolites in plasma), lipid metabolism (increased long-chain fatty acids in the heart tissue and the opposite in plasma), and phospholipid metabolism (high accumulation of phosphocholine precursor metabolites in the heart in comparison to plasma) [14]. Prior analysis of hearts from acutely-infected mice also showed that cardiac metabolite profiles reflected disease severity, with changes in cardiac acylcarnitines and phosphatidylcholines predictive of acute infection outcome [8]. Metabolomic analysis of chronic CD has been limited to serum and gastrointestinal tract samples [17][18]. Serum analysis demonstrated significant changes in amino acid and lipid metabolism, particularly acylcarnitines, sphingolipids, and glycerophospholipids [17]. Analysis of GI tract samples observed persistent metabolic perturbations in the oesophagus and large intestine in chronic CD, including infection-induced elevation of acylcarnitines, phosphatidylcholines and amino acid derivatives [18]. However, metabolic changes in the heart may differ from those in the circulation or in the GI tract [14]. It is therefore essential to perform metabolomic analysis of tissues collected from the heart in chronic CD. Many sudden fatalities due to chronic cardiac CD are often attributed to apical aneurysms which occur at the bottom of the heart [19,20]. We therefore focused on liquid chromatography-tandem mass spectrometry-based metabolomic analysis of horizontally-sectioned hearts from mice chronically infected with *T. cruzi* strains CL and Sylvio X10/4. These samples had previously been analyzed in terms of positional differences (heart apex vs heart base), but not in the context of metabolic changes associated with chronic *T. cruzi* infection [8]. Overall, we observed significant localized chemical differences associated with infection, with a disconnect between parasite localization and overall positional metabolic perturbations. Our data also showed infection-induced variations in acylcarnitine and phosphocholine chemical families.

## Methods

### Ethics statement

All vertebrate animal studies were performed in accordance with the USDA Animal Welfare Act and the Guide for the Care and Use of Laboratory Animals of the National Institutes of Health. The protocol was approved by the University of California San Diego Institutional Animal Care and Use Committee (protocol S14187).

### *In vivo* experimentation

All *in vivo* experimentation, sample collection and qPCR analysis was conducted and previously reported in [8].

### Metabolite extraction and UHPLC-MS/MS

The two-step procedure for metabolite extraction was adapted from Want *et al [21]* and was conducted in McCall *et al [8]*, with “aqueous” and “organic” extracts referring to the metabolites recovered from the first 50% methanol extraction and the second 3:1 dichloromethane-methanol extraction, respectively. UHPLC-MS/MS analysis was conducted using a Thermo Scientific UltiMate 3000 Ultra High Performance Liquid Chromatography with a C8 LC column and MS/MS detection on a Maxis Impact HD QTOF mass spectrometer (Bruker Daltonics), as previously reported [8][21].

### LC-MS/MS data analysis

Data processing was performed as previously reported using Optimus, July 21, 2016 version [8][22]. Total ion current (TIC) normalization was performed in R studio. Principal coordinate analysis (PCoA) was performed on total ion current (TIC) normalized MS1 feature data table using the Bray-Curtis-Faith dissimilarity metric using QIIME1 [23], for both organic and aqueous extractions combined. The three-dimensional PCoA plots were visualized in EMPeror [24]. Heart three-dimensional modelling was completed using ‘ili’ (http://ili.embl.de/) [25]using a three dimensional heart model from 3DCADBrowser.com (http://www.3dcadbrowser.com/).

Global Natural Products Social Molecular Networking (GNPS) was used to perform molecular networking according to the parameters in Table 1 [26]. Cytoscape 3.7.0. was used to visualize the molecular networks [27].

**Table 1:** Global Natural Products Social Molecular Networking (GNPS) parameters.

|  |  |
| --- | --- |
| Parent mass | 2.0 Da |
| MS/MS fragment ion tolerance | 0.5 Da |
| Cosine score | 0.7 |
| Minimum matched fragment ions | 6 |
| Analog search against the library | not allowed |
| Network TopK | 10 |
| Maximum Connected Component Size | 100 |
| Minimum cluster size | 2 |
| Score threshold | 0.7 |
| Library search Min Matched Peaks | 6 |
| Filter precursor window | filter |
| Filter peaks in 50Da window | filter |

Random forest analysis was performed in Jupyter Notebook using R with the number of trees set to 500. Random forest classifier cutoff was based on ranked variable importance score of differential metabolites in combination with unadjusted p-values<0.05 where 4 consecutive non-significant unadjusted p-values defined the cutoff. FDR-corrected Mann Whitney p<0.05 for all positions was also used as an alternate method to determine significant metabolite differences. Venn diagrams were used to visualize the unique and common metabolites differential between CL and Sylvio X10/4 infection, compared to uninfected samples, based on heart segment positions, random forest classifier for all positions, and FDR-corrected Mann Whitney p<0.05 for all positions, using http://bioinformatics.psb.ugent.be/webtools/Venn/.

### Data availability

Metabolomics data has been deposited in MassIVE (http://massive.ucsd.edu/, accession #MSV000080450). Molecular networks can be accessed at https://gnps.ucsd.edu/ProteoSAFe/status.jsp?task=f16bc44c3d5040d098c978823f50c68f (all samples, Aqueous extract), https://gnps.ucsd.edu/ProteoSAFe/status.jsp?task=5f8af6d62d8549358966f3896a81063a (all samples, Organic extract).

## Results

The purpose of this study was to compare the metabolic impact of chronic *T. cruzi* infection in the mouse heart between divergent *T. cruzi* strains and between cardiac regions. To do so, we analyzed previously-collected positive mode LC-MS/MS data from mice chronically infected with *T. cruzi* strain CL or Sylvio X10/4 [8]. While this prior study focused on positional differences between uninfected samples and on the impact of acute infection on the cardiac metabolite profile, here we specifically focused on the impact of chronic infection on the cardiac metabolite profile.

We observed a clear distinction in the impact of *T. cruzi* infection on the overall metabolite profile between *T. cruzi* strains by heart position (Fig 1, S1 figure, S2 figure). As previously described [8], parasite burden was highest at the base of the heart (position A) for strain CL and central positions (position C) for strain Sylvio X10/4 (Fig 1A). PERMANOVA analysis indicated that the highest significant perturbation in the overall metabolite profile occurred at central positions for strain CL infection (PERMANOVA analysis of Bray-Curtis-Faith distance matrix R^2^=0.20813, p-value=0.004 at position C) and at apical positions for strain Sylvio X10/4 infection (PERMANOVA analysis Bray-Curtis-Faith distance matrix R^2^=0.27923, p-value=0.014 at position D) (Fig 1 B). Strikingly, in both cases chemical disturbance was greatest at sites distinct from the- highest parasite burden, which corroborates our observations in the context of chronic gastrointestinal *T. cruzi* infection in mice [18]. The localization of chemical disturbance also provides a molecular mechanism explaining the apical aneurysms observed in CD patients.

**Fig 1.**
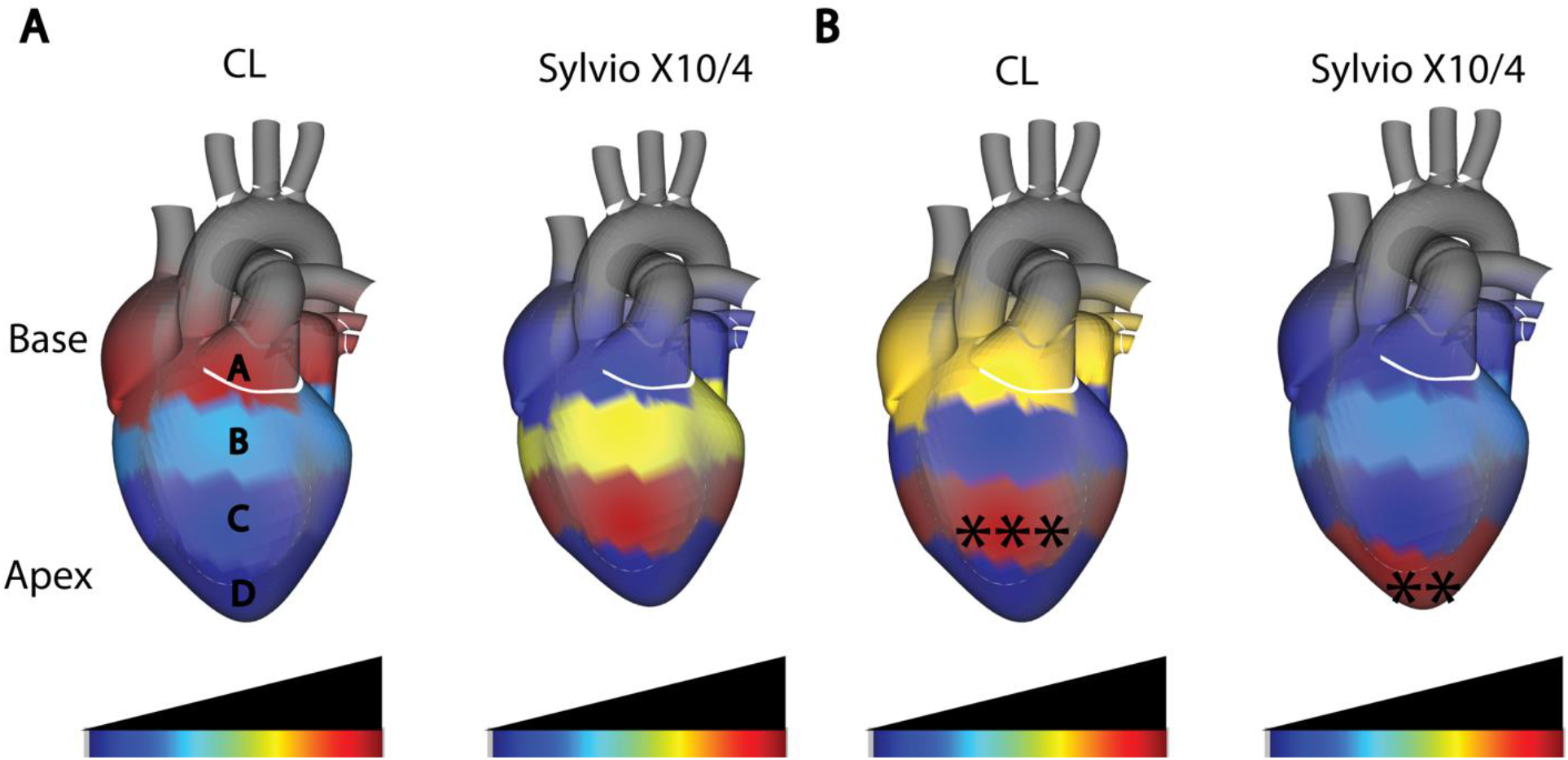
Disconnect between sites of parasite persistence and metabolic alterations in chronic cardiac CD. (A) Median cardiac parasite burden, as determined by qPCR. Parasite burden was highest at the heart base (position A) for strain CL and central heart segments (position C) for strain Sylvio X10/4, indicating parasite strain-specific differences in parasite tropism. (B) Statistically significant perturbations in the overall metabolite profile between uninfected and strain CL-infected (left), and between uninfected and strain Sylvio X10/4-infected mice (right) were analyzed using PERMANOVA. The highest significant metabolite perturbation was at central heart segments (position C) for strain CL (***, p < 0.001 by PERMANOVA) and at the heart apex (position D) for strain Sylvio X10/4 (**, p < 0.01 by PERMANOVA).

To identify the specific cardiac metabolites spatially perturbed by infection, initially we built a random forest classifier for each position, each strain and each extraction method, comparing to uninfected matched control samples (S1-S6 tables). We first assessed the overlap between the top-ranked most differential metabolites by random forest for the two different strains, as described in Methods. Limited overlap of these significant metabolites was observed between strains (Figure 2). However, annotation of these differential metabolites using molecular networking through the GNPS platform [26] revealed that while differing in terms of *m/z,* many were part of the same chemical families, including acylcarnitines and phosphocholine (S1 – S6 Table; S3 and S4 figure).

**Fig 2.**
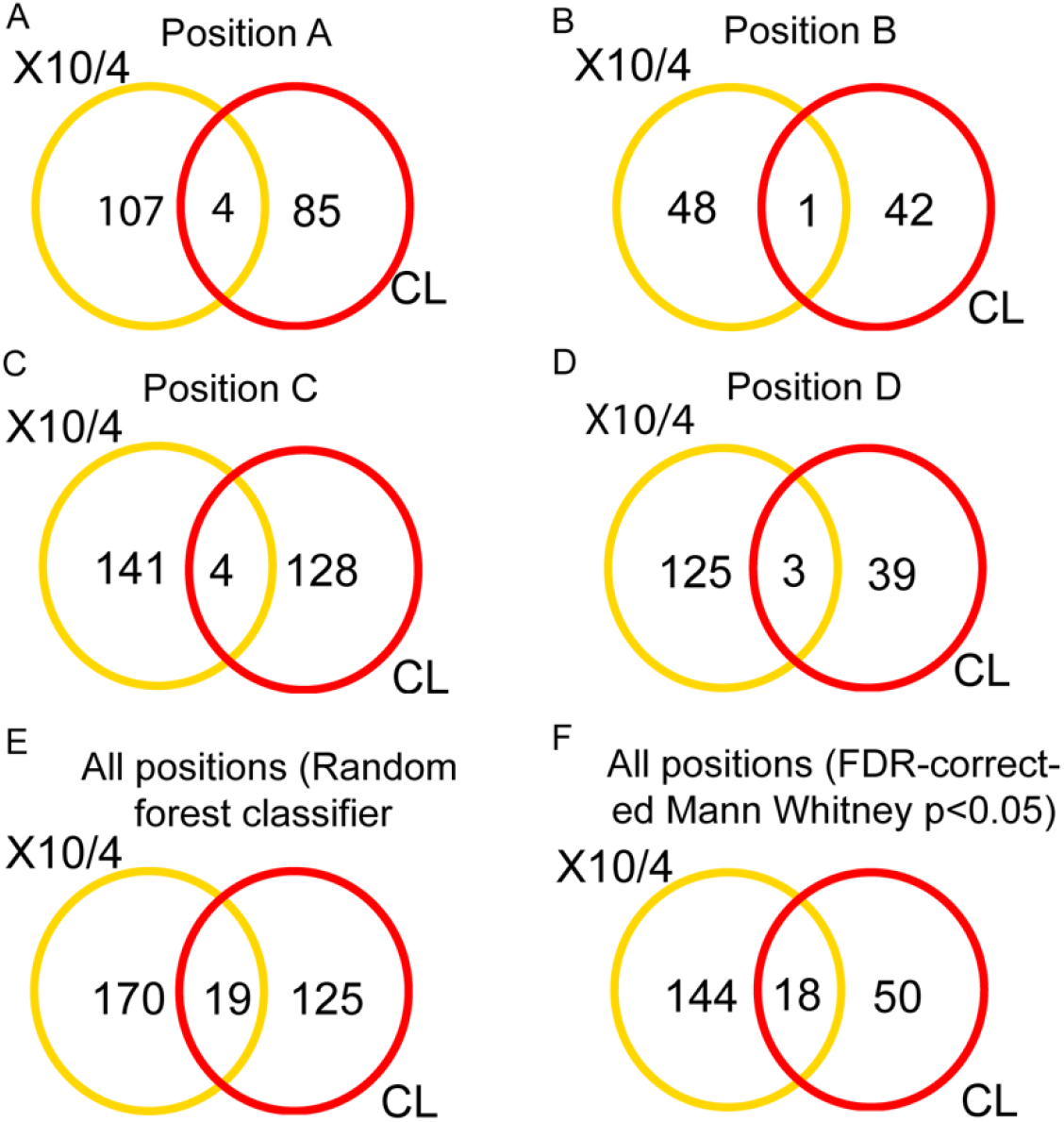
Limited overlap of specific differential metabolites between strains. Yellow and red circles represent differential metabolites between strain Sylvio X10/4-infected and matched uninfected controls, and between strain CL-infected and matched uninfected controls, respectively. Intersect are metabolites impacted by infection in both strains. (A-D) Differential metabolites for each strain, at given heart positions. (E) Metabolites impacted by infection with each strain, irrespective of position, as determined by random forest classifier, with variable importance score cutoff as described in Methods. (F) Metabolites impacted by infection with each strain, irrespective of heart position using FDR-corrected Mann Whitney p<0.05 cutoff.

Random forest classifier identified several acylcarnitines and phosphocholines as impacted by infection (S1 – S5 Tables). Both chemical families play a major role in several biochemical pathways. Carnitine serves as a shuttling mechanism for fatty acids, in the form of acylcarnitines, from the cytosol into the matrix of the mitochondria for beta-oxidation [28]. Phosphocholines are major components of lipid metabolism, cell membrane structure, and choline production, the latter of which is essential for select amino acid and neurotransmitter synthesis [29,30].

Total acylcarnitines in central positions of the heart were decreased by strain Sylvio X10/4 infection compared to the uninfected group (Fig 3 A and B, Mann-Whitney p<0.05). A similar trend was observed for total acylcarnitines following strain CL infection when compared to matched uninfected samples, even though this difference was not statistically significant (Fig 3A and B). Previous studies demonstrated that acylcarnitines of different lengths were associated with infection outcome in acute *T. cruzi* mouse models [8]. Therefore, we sought to understand how different length acylcarnitines were affected by chronic infection. Acylcarnitines are classified based on the number of carbons in their fatty acid chain as short- (≤C4), mid- (C5 – C11), and long-chain (≥C12) acylcarnitines.

Central and apical positions (positions B, C and D) had the largest abundance of mid and long chain acylcarnitines in both CL and Sylvio X10/4 strain compared to the heart base (Mann Whitney p<0.05) (Fig 3 C-H). In the case of CL strain infection, when compared to uninfected samples, short chain acylcarnitines were significantly decreased at the heart base and apex (positions A and D, p<0.05 Mann-Whitney)(Fig 3 C and D). Mid-chain acylcarnitine levels were decreased by strain CL infection compared to uninfected samples at central positions (position B, p<0.05 Mann-Whitney) (Fig E and F). In contrast, long chain acylcarnitines were significantly increased at central and apical positions (positions C and D respectively) by strain CL infection (Mann-Whitney p<0.05) (Fig 3 G and H). Strain Sylvio X10/4 infection significantly decreased mid chain acylcarnitine at all positions (Mann-Whitney p<0.05)(Fig 3 E and F).

**Fig 3.**
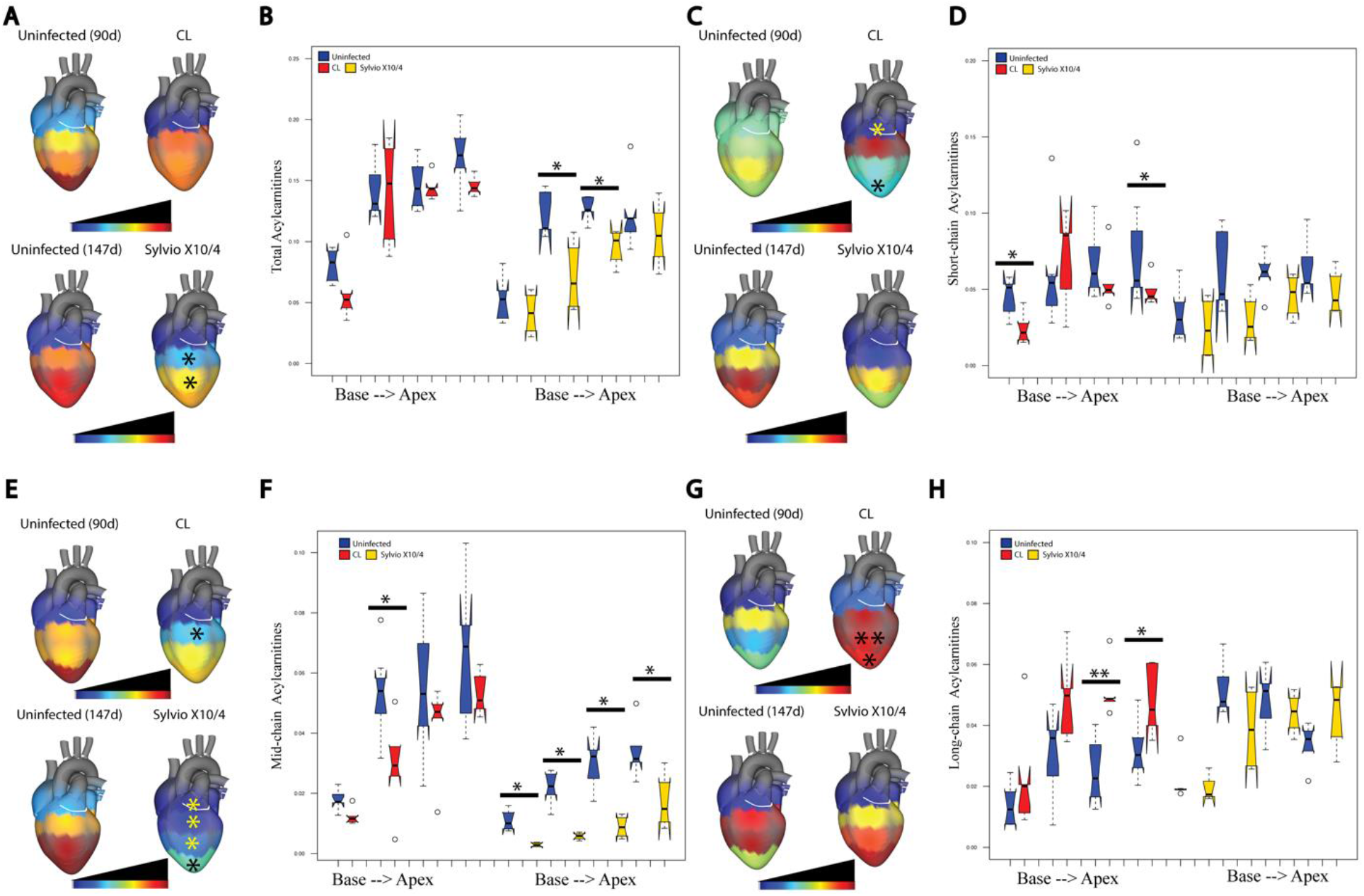
Spatial impact of chronic *T. cruzi* infection on cardiac acylcarnitines. (A) and (B) Differential total acylcarnitine distribution between uninfected and infected heart sections for both CL and Sylvio X10/4 strains. (*, p<0.05 by Wilcoxon-Mann-Whitney). (C, D) CL-infected mice showed statistically significant decreases (*, p<0.05 by Mann-Whitney test) in short-chain acylcarnitine (≤ C4) at heart positions A and D. (E, F) Both strains of *T. cruzi* showed statistically significant decreases in mid-chain acylcarnitines. (G, H) CL-infected mice increased long-chain acylcarnitines (≥C12) at position C (**, p<0.01) and D (*, p<0.05).

CL strain infection significantly increased total phosphocholines at central position C compared to uninfected samples (Mann-Whitney p<0.01), with a similar but non-significant trend for strain Sylvio X10/4 infection at the heart apex (Fig 4 A and B). Further analysis based on phosphocholine mass was performed, because previous studies showed differences in phosphocholine mass range between fatal and non-fatal acute mouse infection [8]. Phosphocholines were categorized into four mass ranges: short (200 – 400 *m/z*), mid (401 – 600 *m/z*), long (601 – 800 *m/z*), and very long (>801 *m/z*). Significantly elevated short phosphocholines were observed in central heart positions (position C) for Sylvio X10/4 infection compared to uninfected samples (Fig 4 C and D). This same pattern was also observed for CL strain infection in mid and very long phosphocholines at the same position (p <0.01 and <0.05, respectively), when compared to uninfected samples (Fig 4 E and F, I and J).

**Fig 4.**
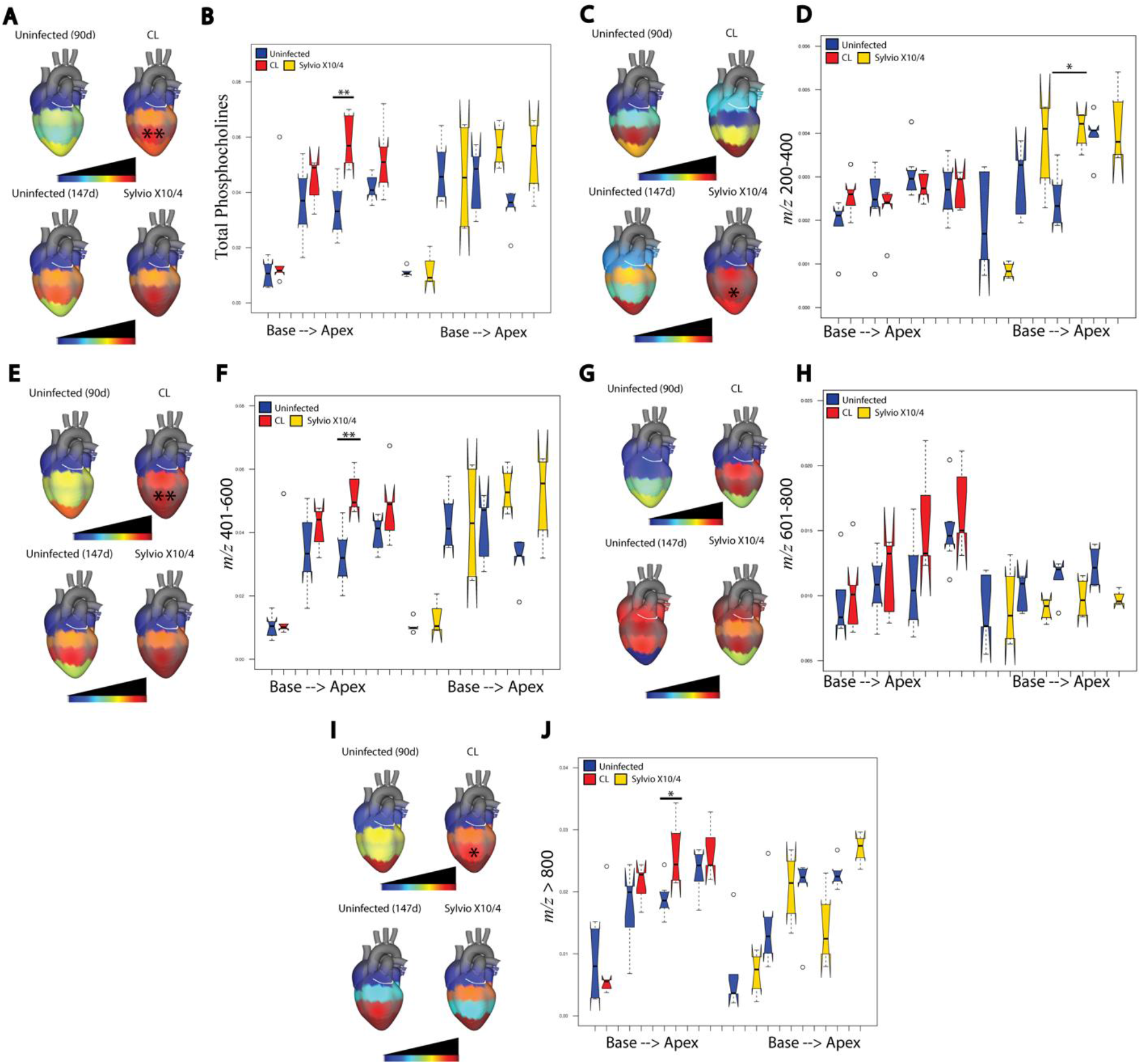
Spatial impact of chronic *T. cruzi* infection on cardiac phosphocholines. (A, B) Statistically significant differences in total phosphocholine levels were identified in mice infected with CL strain at heart position C when compared to uninfected mice (**, p<0.01 by Wilcoxon-Mann-Whitney test). (C,D) Sylvio X10/4-infected mice showed statistically significant differences (*, p<0.05 by Wilcoxon-Mann-Whitney test) in small phosphocholines (200-400 *m/z*) at heart position C. (E, F) Only CL-infected mice showed statistically significant differences (**, p<0.01 by Wilcoxon-Mann-Whitney test) for mid-sized phosphocholines (401-600 *m/z*) at position C. (G, H) Large phosphocholines (601-800 *m/z*) were not affected by infection for both strains. (I, J) CL-infected mice showed a statistically significant difference (*, p<0.05 by Wilcoxon-Mann-Whitney test) in very long phosphocholines (> 801 *m/z*) at position C.

## Discussion

Currently, there are 7 *T. cruzi* discrete typing units (DTUs TcI – TcVI and Tcbat). TcI to TcVI are infectious to humans [31]. These DTUs, while still currently considered the same species, nevertheless present significant genetic differences [31][32]. However, pathogenic processes are overall similar in cardiac CD across *T. cruzi* strains, with accumulation of fibrosis and inflammation, although timing and magnitude of symptoms may be different depending on parasite and host characteristics [32][33]. These similarities are reflected in the common metabolomic changes observed for strain Sylvio X10/4 (TcI) and strain CL (TcVI)-infected heart tissue in this study, including chronic infection-induced increases in phosphocholines and decreases in acylcarnitines.

Our results also highlight the importance of considering metabolic changes at the level of chemical families, beyond just individual metabolites. While there was little overlap of highly significant metabolite *m/z* at each position between strains, most differential metabolites were from these two chemical families. McCall *et al.* described these two chemical families as discriminatory compounds between fatal and non-fatal acute *T. cruzi* infected heart tissue [8]. Considering acute stage infection progresses into chronic stage infection, it is not surprising that changes in the relative abundance of these molecules are also observed in chronic CD. Phosphocholines have been linked to coronary heart disease due to production of lysophosphatidylcholines and choline. [29,34]. Increased acylcarnitine levels have been linked to cardiovascular disease as well as cardiac symptoms in those already possessing a cardiac disease [35,36]. However, our results show the opposite pattern compared to non-infectious heart disease, highlighting the need to specifically study CD rather than extrapolate from other cardiac conditions (Fig 3). In a study addressing gene expression differences between human CD cardiomyopathy and dilated cardiomyopathy, there was an upregulation of gene expression associated with lipid metabolism from heart samples of human cardiac CD patients, while the opposite was seen in non-infectious dilated cardiomyopathy patient samples [37]. Higher lipid metabolism would increase acylcarnitine catabolism and thus decrease overall acylcarnitine abundance. Decreased carnitine palmitoyltransferase and acetyltransferase levels, as observed by proteomic analysis of infected mouse heart tissue [38], may alternatively also contribute to the decreased acylcarnitine levels we observed.

Interestingly, in a study on the effects of diet on chronic *T. cruzi* mouse infection, a similar pattern was observed as in our study, where serum acylcarnitines were amongst the most differential compounds in infected samples compared to uninfected samples, with most short- and mid-chain acylcarnitines decreased, and select long-chain acylcarnitines increased [17]. In addition, significant acylcarnitine differences were seen in the gastrointestinal tract of acute *T. cruzi-*infected mice [18]. Likewise, both long chain acylcarnitines and phospholipid synthesis were increased in the heart tissue of acutely infected mice in prior studies [14]. Thus, our data agree with and expand upon the existing *T. cruzi* metabolomics literature. Overall, the different patterns in metabolites we observe here in contrast to other cardiac diseases is consistent with differences in gene expression in humans with CD compared to other diseases.

Differences in pathogenesis between strains may be due to differential strain tropism. Indeed, TcI strains tend to produce cardiomyopathy, while TcVI strains commonly produce megacolon and megaesophagus, although cardiomyopathy can still occur [39]. Our results indicate a disconnect between sites of highest parasite burden and sites of metabolic perturbation. Although parasite levels were highest in central heart segments following strain Sylvio X10/4 infection, we observed statistically significant perturbations in metabolism at the apex of the heart (Fig 1). Apical aneurysms are one of the major symptoms in chronic CD patients [40]. In addition, lateral heart wall damage is also common among chronic CD patients, in central regions of the heart [41], and we observed significant perturbations in cardiac metabolism at lower central heart positions in strain CL infection (Fig 1B) [41]. Based on these results, we propose a concept of spatial disease tolerance, whereby some tissue regions are more affected by infection, while others are less functionally affected. This is likely due to a combination of host and pathogen factors, given the differences we observe here between strain CL and strain Sylvio X10/4 infection in the same C3H mouse genetic background. Importantly, the localization of maximal metabolic perturbation in acute strain Sylvio X10/4 infection was also the heart apex, indicating that the spatial course of disease may be set early in CD [8]. Likewise, host factors likely contribute, such as the higher production of antiparasitic but tissue-damaging IFNγ at the heart apex or specific cardiac regions being more prone to microvasculature disruptions [8].

These results set a foundation for biomarker studies and for host-directed therapeutic development. CD may be particularly amenable to such treatment strategies, due to the contribution of host-mediated tissue damage to CD pathogenesis [1,7]. Indeed, we have previously shown that carnitine supplementation can be used to treat acute CD [18]. Our observation of decreases in cardiac acylcarnitines in chronic CD indicate that this approach may also be useful to treat chronic CD. Importantly the fact that acylcarnitines are affected in both chronic CL and Sylvio X10/4 infection suggests broad applicability. Other studies have emphasized the impact of metabolism modulators on CD progression. High fat diet reduces parasite levels and increases survival in acute CD mouse models [42]. Treatment of acutely *T. cruzi* infected mice with metformin (a metabolic modulator used to treat diabetic patients) also led to an increase in overall survival rate and decreased p blood parasitemia [43].

In addition to the need for novel treatments, several studies have highlighted the importance of novel diagnostic methods for CD [44,45]. Current diagnostic methods rely on serological and microscopic exams and polymerase chain reaction (PCR) [46]. In addition, during the chronic stage, parasite levels decrease drastically therefore PCR techniques have to be used instead of microscopy. PCR however only detects the presence of infection but not cardiac damage [46]. Hence, biomarkers in the form of small molecules or chemical families, as identified in this study, can aid in addressing this issue. Future work will investigate whether the infection-induced perturbations observed here in the heart are also detectable in clinically-accessible biofluids.

Due to the low parasite burden in chronic Chagas disease and instrumental limits of detection, we anticipate most if not all detected metabolites to be host-derived, supported by their detection in uninfected tissues. As such, this study is focused on the impact of *T. cruzi* infection on host metabolism. A further limitation is that many of the differential metabolites were not annotatable, as is usual in metabolomic studies [47]. Nevertheless, we were able to annotate metabolites affected by chronic infection that make up important host biochemical pathways.

Overall, our study highlights the importance of not only identifying overall differences but also positional metabolic differences associated with multiple *T. cruzi* strains, and the strength of systematic chemical cartography in understanding disease tropism and how it differs from pathogen tropism. These results will serve as stepping stones for further CD drug development and biomarker discovery, something that is urgently needed.

## Supporting information

**S1 Table. Annotated metabolites of combined extracts perturbed by infection at position A, identified through random forest classifier.**

**S2 Table. Annotated metabolites of combined extracts perturbed by infection at position B, identified through random forest classifier.**

**S3 Table. Annotated metabolites of combined extracts perturbed by infection at position C, identified through random forest classifier.**

**S4 Table. Annotated metabolites of combined extracts perturbed by infection at position D, identified through random forest classifier.**

**S5 Table. Annotated metabolites of combined extracts perturbed by infection at positions A-D, identified through random forest classifier.**

**S6 Table. Annotated metabolites of combined extracts identified as perturbed by infection at all positions (FDR-corrected Mann Whitney p<0.05).**

**S1 Figure. Principal coordinate analysis plot of *T. cruzi* strain CL infected (red) and uninfected (blue) heart tissue samples.** Statistically different clustering found in position C (PERMANOVA p-value<0.05).

**S2 Figure. Principal coordinate analysis plot of *T. cruzi* strain Sylvio X10/4 infected (gold) and uninfected (blue) heart tissue samples.** Statistically different clustering found in position D (PERMANOVA p-value<0.05).

**S3 Figure. Sub-molecular networks and mirror plot of aqueous and organic extract acylcarnitines and phosphocholines.** Each pie chart is one metabolite colored by MS2 spectral count in CL-infected and Sylvio X10/4-infected samples where red is CL and gold is Sylvio X10/4. (A) Subnetwork of aqueous extract acylcarnitines with representative acylcarnitine mirror plot (acetylcarnitine, *m/z* -204.124). (B) Subnetwork of aqueous extract phosphocholines with representative phosphocholine mirror plot (Spectral match to 1-Hexadecanoyl-2-(9Z-octadecenoyl)-sn-glycero-3-phosphocholine, *m/z* 758.65). (C) Subnetwork of organic extract acylcarnitines with representative acylcarnitine mirror plot (acetylcarnitine, *m/z*- 204.126). (D) Subnetwork of organic extract phosphocholines with representative phosphocholine mirror plot (Spectral Match to 1-Oleoyl-2-palmitoyl-sn-glycero-3-phosphocholine, *m/z* -760.601).

**S4 Figure. GNPS mirror plots of annotated metabolites.** (A) mirror plot of *m/z* 703.575, RT 286s (top, black) to reference library spectrum (SM(d18:1/16:0), bottom, green). (B) mirror plot of *m/z* 454.294, RT 206s (top, black) to reference library spectrum (hexadecanoyl-lysophosphatidylethanolamine, bottom, green). (C) mirror plot of *m/z* 377.146, RT 137s (top, black) to reference library spectrum (riboflavin, bottom, green). (D) mirror plot of *m/z* 646.614, RT 417s (top, black) to reference library spectrum (ceramide, bottom, green). (E) mirror plot of *m/z* 716.523, RT 395s (top, black) to reference library spectrum (1-palmitoyl-2-oleoyl-sn-glycero-3-phosphoethanolamine, bottom, green).

## References

1. Rassi A Jr, Rassi A, Marin-Neto JA. Chagas disease. Lancet. 2010;375: 1388–1402.

2. Bern C. Chagas’ Disease. The New England journal of medicine. 2015. p. 1882.

3. Beatty NL, Perez-Velez CM, Yaglom HD, Carson S, Liu E, Khalpey ZI, et al. Evidence of Likely Autochthonous Transmission of Chagas Disease in Arizona. Am J Trop Med Hyg. 2018;99: 1534–1536.

4. Gunter SM, Murray KO, Gorchakov R, Beddard R, Rossmann SN, Montgomery SP, et al. Likely Autochthonous Transmission of Trypanosoma cruzi to Humans, South Central Texas, USA. Emerg Infect Dis. 2017;23: 500–503.

5. Miller DA, Hernandez S, De Armas LR, Eells SJ, Traina MM, Miller LG, et al. Tolerance of Benznidazole in a United States Chagas Disease Clinic. Clinical Infectious Diseases. 2015. pp. 1237–1240. doi:10.1093/cid/civ005

6. Forsyth CJ, Hernandez S, Olmedo W, Abuhamidah A, Traina MI, Sanchez DR, et al. Safety Profile of Nifurtimox for Treatment of Chagas Disease in the United States. Clin Infect Dis. 2016;63: 1056–1062.

7. Tarleton RL. Chagas disease: a role for autoimmunity? Trends in Parasitology. 2003. pp. 447–451. doi:10.1016/j.pt.2003.08.008

8. McCall L-I, Morton JT, Bernatchez JA, de Siqueira-Neto JL, Knight R, Dorrestein PC, et al. Mass Spectrometry-Based Chemical Cartography of a Cardiac Parasitic Infection. Anal Chem. 2017;89: 10414–10421.

9. Lewis MD, Kelly JM. Putting Infection Dynamics at the Heart of Chagas Disease. Trends Parasitol. 2016;32: 899–911.

10. McCall L-I, McKerrow JH. Determinants of disease phenotype in trypanosomatid parasites. Trends Parasitol. 2014;30: 342–349.

11. Breitling R, Bakker BM, Barrett MP, Decuypere S, Dujardin J-C. Metabolomic Systems Biology of Protozoan Parasites. Genetics Meets Metabolomics. 2012. pp. 73–84. doi:10.1007/978-1-4614-1689-0_6

12. Caradonna KL, Engel JC, Jacobi D, Lee C-H, Burleigh BA. Host metabolism regulates intracellular growth of Trypanosoma cruzi. Cell Host Microbe. 2013;13: 108–117.

13. Cestari I, Haas P, Moretti NS, Schenkman S, Stuart K. Chemogenetic Characterization of Inositol Phosphate Metabolic Pathway Reveals Druggable Enzymes for Targeting Kinetoplastid Parasites. Cell Chem Biol. 2016;23: 608–617.

14. Gironès N, Carbajosa S, Guerrero NA, Poveda C, Chillón-Marinas C, Fresno M. Global metabolomic profiling of acute myocarditis caused by Trypanosoma cruzi infection. PLoS Negl Trop Dis. 2014;8: e3337.

15. Knubel CP, Martínez FF, Acosta Rodríguez EV, Altamirano A, Rivarola HW, Diaz Luján C, et al. 3-Hydroxy kynurenine treatment controls T. cruzi replication and the inflammatory pathology preventing the clinical symptoms of chronic Chagas disease. PLoS One. 2011;6: e26550.

16. Garg N, Gerstner A, Bhatia V, DeFord J, Papaconstantinou J. Gene expression analysis in mitochondria from chagasic mice: alterations in specific metabolic pathways. Biochem J. 2004;381: 743–752.

17. Lizardo K, Ayyappan JP, Ganapathi U, Dutra WO, Qiu Y, Weiss LM, et al. Diet Alters Serum Metabolomic Profiling in the Mouse Model of Chronic Chagas Cardiomyopathy. Dis Markers. 2019;2019: 4956016.

18. Hossain E, Khanam S, Wu C, Lostracco-Johnson S, Thomas D, Katemauswa M, et al. 3D mapping of host-parasite-microbiome interactions reveals metabolic determinants of tissue tropism and disease tolerance in Chagas disease. doi:10.1101/727917

19. Jr AR, Rassi A Jr, Rassi A, Little WC. Chagas’ Heart Disease. Clinical Cardiology. 2000. pp. 883–889. doi:10.1002/clc.4960231205

20. Acquatella H. Echocardiography in Chagas heart disease. Circulation. 2007;115: 1124–1131.

21. Want EJ, Masson P, Michopoulos F, Wilson ID, Theodoridis G, Plumb RS, et al. Global metabolic profiling of animal and human tissues via UPLC-MS. Nat Protoc. 2013;8: 17–32.

22. Sturm M, Bertsch A, Gröpl C, Hildebrandt A, Hussong R, Lange E, et al. OpenMS – an open-source software framework for mass spectrometry. BMC Bioinformatics. 2008;9: 163.

23. Caporaso JG, Kuczynski J, Stombaugh J, Bittinger K, Bushman FD, Costello EK, et al. QIIME allows analysis of high-throughput community sequencing data. Nat Methods. 2010;7: 335–336.

24. Vázquez-Baeza Y, Pirrung M, Gonzalez A, Knight R. EMPeror: a tool for visualizing high-throughput microbial community data. Gigascience. 2013;2: 16.

25. Protsyuk I, Melnik AV, Nothias L-F, Rappez L, Phapale P, Aksenov AA, et al. 3D molecular cartography using LC-MS facilitated by Optimus and ’ili software. Nat Protoc. 2018;13: 134–154.

26. Wang M, Carver JJ, Phelan VV, Sanchez LM, Garg N, Peng Y, et al. Sharing and community curation of mass spectrometry data with Global Natural Products Social Molecular Networking. Nat Biotechnol. 2016;34: 828–837.

27. Shannon P, Markiel A, Ozier O, Baliga NS, Wang JT, Ramage D, et al. Cytoscape: a software environment for integrated models of biomolecular interaction networks. Genome Res. 2003;13: 2498–2504.

28. Longo N, Frigeni M, Pasquali M. Carnitine transport and fatty acid oxidation. Biochim Biophys Acta. 2016;1863: 2422–2435.

29. Tang WHW, Wilson Tang WH, Wang Z, Levison BS, Koeth RA, Britt EB, et al. Intestinal Microbial Metabolism of Phosphatidylcholine and Cardiovascular Risk. New England Journal of Medicine. 2013. pp. 1575–1584. doi:10.1056/nejmoa1109400

30. van der Veen JN, Kennelly JP, Wan S, Vance JE, Vance DE, Jacobs RL. The critical role of phosphatidylcholine and phosphatidylethanolamine metabolism in health and disease. Biochim Biophys Acta Biomembr. 2017;1859: 1558–1572.

31. Brenière SF, Waleckx E, Barnabé C. Over Six Thousand Trypanosoma cruzi Strains Classified into Discrete Typing Units (DTUs): Attempt at an Inventory. PLoS Negl Trop Dis. 2016;10: e0004792.

32. Dorn PL, McClure AG, Gallaspy MD, Waleckx E, Woods AS, Monroy MC, et al. The diversity of the Chagas parasite, Trypanosoma cruzi, infecting the main Central American vector, Triatoma dimidiata, from Mexico to Colombia. PLoS Negl Trop Dis. 2017;11: e0005878.

33. Messenger LA, Miles MA, Bern C. Between a bug and a hard place: Trypanosoma cruzi genetic diversity and the clinical outcomes of Chagas disease. Expert Rev Anti Infect Ther. 2015;13: 995–1029.

34. Ganna A, Salihovic S, Sundström J, Broeckling CD, Hedman AK, Magnusson PKE, et al. Large-scale metabolomic profiling identifies novel biomarkers for incident coronary heart disease. PLoS Genet. 2014;10: e1004801.

35. Guasch-Ferré M, Zheng Y, Ruiz-Canela M, Hruby A, Martínez-González MA, Clish CB, et al. Plasma acylcarnitines and risk of cardiovascular disease: effect of Mediterranean diet interventions. Am J Clin Nutr. 2016;103: 1408–1416.

36. Strand E, Pedersen ER, Svingen GFT, Olsen T, Bjørndal B, Karlsson T, et al. Serum Acylcarnitines and Risk of Cardiovascular Death and Acute Myocardial Infarction in Patients With Stable Angina Pectoris. J Am Heart Assoc. 2017;6. doi:10.1161/JAHA.116.003620

37. Cunha-Neto E, Dzau VJ, Allen PD, Stamatiou D, Benvenutti L, Higuchi ML, et al. Cardiac gene expression profiling provides evidence for cytokinopathy as a molecular mechanism in Chagas’ disease cardiomyopathy. Am J Pathol. 2005;167: 305–313.

38. Wozniak JM, Silva TA, Thomas D, Siqueira-Neto JL, McKerrow JH, Gonzalez DJ, et al. Molecular dissection of chagas induced cardiomyopathy reveals central disease associated and druggable signaling pathways. PLoS Negl Trop Dis. 2020;14: e0007980.

39. Ramos-Ligonio A, Torres-Montero J, López-Monteon A, Dumonteil E. Extensive diversity of Trypanosoma cruzi discrete typing units circulating in Triatoma dimidiata from central Veracruz, Mexico. Infection, Genetics and Evolution. 2012. pp. 1341–1343. doi:10.1016/j.meegid.2012.04.024

40. Marin-Neto JA, Cunha-Neto E, Maciel BC, Simões MV. Pathogenesis of chronic Chagas heart disease. Circulation. 2007;115: 1109–1123.

41. Lee-Felker SA, Thomas M, Felker ER, Traina M, Salih M, Hernandez S, et al. Value of cardiac MRI for evaluation of chronic Chagas disease cardiomyopathy. Clin Radiol. 2016;71: 618.e1–7.

42. Harris EV, de Roode JC, Gerardo NM. Diet–microbiome–disease: Investigating diet’s influence on infectious disease resistance through alteration of the gut microbiome. PLOS Pathogens. 2019. p. e1007891. doi:10.1371/journal.ppat.1007891

43. Brima W, Eden DJ, Mehdi SF, Bravo M, Wiese MM, Stein J, et al. The brighter (and evolutionarily older) face of the metabolic syndrome: evidence from Trypanosoma cruzi infection in CD-1 mice. Diabetes Metab Res Rev. 2015;31: 346–359.

44. Alonso-Padilla J, Cortés-Serra N, Pinazo MJ, Bottazzi ME, Abril M, Barreira F, et al. Strategies to enhance access to diagnosis and treatment for Chagas disease patients in Latin America. Expert Rev Anti Infect Ther. 2019;17: 145–157.

45. Picado A, Angheben A, Marchiol A, Alarcón de Noya B, Flevaud L, Pinazo MJ, et al. Development of Diagnostics for Chagas Disease: Where Should We Put Our Limited Resources? PLoS Negl Trop Dis. 2017;11: e0005148.

46. Keating SM, Deng X, Fernandes F, Cunha-Neto E, Ribeiro AL, Adesina B, et al. Inflammatory and cardiac biomarkers are differentially expressed in clinical stages of Chagas disease. Int J Cardiol. 2015;199: 451–459.

47. da Silva RR, Dorrestein PC, Quinn RA. Illuminating the dark matter in metabolomics. Proceedings of the National Academy of Sciences of the United States of America. 2015. pp. 12549–12550.

