## Supplemental Information for "Spatial metabolomics identifies localized chemical changes in heart tissue during chronic cardiac Chagas disease"

### Supporting Information

**S1 Table. Annotated metabolites of combined extracts perturbed by infection at position A, identified through random forest classifier.**

| Top-ranking metabolites for CL *T.cruzi* strain that met the random forest cut-off criteria | | | | | | | | | |
|---|---|---|---|---|---|---|---|---|---|
| ***m/z*** | **RT (sec)** | **Annotation** | **Cosine Score** | **No. of Shared Peaks** | **Mass Diff. to Library Reference** | **ppm error** | **Impact of Infection at position A (Sylvio X10/4 vs uninfected 147 days)** | **Impact of Infection at position A (CL vs uninfected 90 days)** | **Extract** |
| 153.042 | 32 | NA | NA | NA | NA | NA | N/S | N/S | aqueous |
| 180.103 | 143 | NA | NA | NA | NA | NA | N/S | N/S | aqueous |
| 188.129 | 148 | NA | NA | NA | NA | NA | N/S | major decrease | aqueous |
| 189.077 | 142 | NA | NA | NA | NA | NA | N/S | minor increase | aqueous |
| 204.124 | 24 | C2:0 acylcarnitine (acetylcarnitine)[1] | NA | NA | NA | -0.65 | N/S | major decrease | aqueous |
| 211.145 | 151 | NA | NA | NA | NA | NA | N/S | major decrease | aqueous |
| 216.197 | 163 | NA | NA | NA | NA | NA | N/S | N/S | aqueous |
| 225.149 | 173 | NA | NA | NA | NA | NA | N/S | N/S | aqueous |
| 226.181 | 168 | NA | NA | NA | NA | NA | N/S | N/S | aqueous |
| 241.155 | 26 | NA | NA | NA | NA | NA | major increase | minor decrease | aqueous |
| 248.151 | 47 | C4:0-OH acylcarnitine[1] | NA | NA | NA | 2.64 | N/S | major decrease | aqueous |
| 250.095 | 66 | NA | NA | NA | NA | NA | major increase | major decrease | aqueous |
| 251.103 | 147 | NA | NA | NA | NA | NA | N/S | major decrease | aqueous |
| 252.233 | 190 | NA | NA | NA | NA | NA | N/S | major decrease | aqueous |
| 256.229 | 193 | NA | NA | NA | NA | NA | N/S | major decrease | aqueous |
| 270.244 | 200 | NA | NA | NA | NA | NA | N/S | major decrease | aqueous |
| 275.054 | 40 | NA | NA | NA | NA | NA | N/S | major increase | aqueous |
| 278.248 | 179 | NA | NA | NA | NA | NA | N/S | major decrease | aqueous |
| 280.264 | 206 | NA | NA | NA | NA | NA | N/S | N/S | aqueous |
| 286.140 | 137 | NA | NA | NA | NA | NA | N/S | N/S | aqueous |

| | | | | | | | | | |
|---|---|---|---|---|---|---|---|---|---|
| 286.202 | 154 | C8:1 acylcarnitine[1] | NA | NA | NA | -1.33 | N/S | minor decrease | aqueous |
| 288.254 | 187 | NA | NA | NA | NA | NA | N/S | major decrease | aqueous |
| 291.071 | 32 | NA | NA | NA | NA | NA | N/S | minor decrease | aqueous |
| 292.227 | 199 | NA | NA | NA | NA | NA | N/S | major decrease | aqueous |
| 295.263 | 261 | NA | NA | NA | NA | NA | N/S | major increase | aqueous |
| 296.148 | 207 | NA | NA | NA | NA | NA | N/S | N/S | aqueous |
| 296.259 | 187 | NA | NA | NA | NA | NA | N/S | major decrease | aqueous |
| 298.275 | 213 | NA | NA | NA | NA | NA | N/S | major decrease | aqueous |
| 304.652 | 208 | NA | NA | NA | NA | NA | N/S | N/S | aqueous |
| 307.045 | 22 | NA | NA | NA | NA | NA | N/S | N/S | aqueous |
| 312.254 | 194 | NA | NA | NA | NA | NA | N/S | major decrease | aqueous |
| 315.666 | 137 | NA | NA | NA | NA | NA | N/S | N/S | aqueous |
| 316.285 | 191 | NA | NA | NA | NA | NA | N/S | major decrease | aqueous |
| 318.191 | 190 | NA | NA | NA | NA | NA | N/S | N/S | aqueous |
| 320.256 | 206 | NA | NA | NA | NA | NA | N/S | major decrease | aqueous |
| 320.256 | 217 | NA | NA | NA | NA | NA | N/S | major decrease | aqueous |
| 327.234 | 214 | NA | NA | NA | NA | NA | N/S | minor decrease | aqueous |
| 332.280 | 175 | NA | NA | NA | NA | NA | N/S | major decrease | aqueous |
| 343.296 | 177 | NA | NA | NA | NA | NA | N/S | major decrease | aqueous |
| 380.210 | 258 | NA | NA | NA | NA | NA | N/S | minor decrease | aqueous |
| 443.083 | 24 | NA | NA | NA | NA | NA | N/S | N/S | aqueous |
| 503.307 | 145 | NA | NA | NA | NA | NA | N/S | minor decrease | aqueous |
| 605.357 | 169 | NA | NA | NA | NA | NA | N/S | major decrease | aqueous |
| 619.339 | 163 | NA | NA | NA | NA | NA | N/S | major decrease | aqueous |
| 666.139 | 329 | NA | NA | NA | NA | NA | N/S | major decrease | aqueous |
| 931.451 | 189 | NA | NA | NA | NA | NA | N/S | major decrease | aqueous |
| 1231.347 | 181 | NA | NA | NA | NA | NA | N/S | major decrease | aqueous |
| 134.455 | 39 | NA | NA | NA | NA | NA | N/S | minor increase | organic |
| 188.068 | 46 | NA | NA | NA | NA | NA | N/S | N/S | organic |
| 192.159 | 19 | NA | NA | NA | NA | NA | N/S | N/S | organic |
| 195.094 | 139 | NA | NA | NA | NA | NA | N/S | major increase | organic |
| 254.249 | 221 | NA | NA | NA | NA | NA | N/S | N/S | organic |

| | | | | | | | | | |
|---|---|---|---|---|---|---|---|---|---|
| 269.090 | 26 | NA | NA | NA | NA | NA | N/S | major increase | organic |
| 280.265 | 177 | NA | NA | NA | NA | NA | N/S | major decrease | organic |
| 280.265 | 250 | NA | NA | NA | NA | NA | N/S | major decrease | organic |
| 284.239 | 155 | NA | NA | NA | NA | NA | N/S | major decrease | organic |
| 291.148 | 142 | NA | NA | NA | NA | NA | major decrease | N/S | organic |
| 292.177 | 142 | NA | NA | NA | NA | NA | N/S | N/S | organic |
| 296.259 | 162 | NA | NA | NA | NA | NA | N/S | major decrease | organic |
| 296.259 | 181 | NA | NA | NA | NA | NA | N/S | major decrease | organic |
| 298.275 | 190 | NA | NA | NA | NA | NA | N/S | major decrease | organic |
| 308.296 | 238 | NA | NA | NA | NA | NA | N/S | major decrease | organic |
| 320.257 | 190 | NA | NA | NA | NA | NA | N/S | major decrease | organic |
| 326.379 | 206 | NA | NA | NA | NA | NA | N/S | major decrease | organic |
| 326.379 | 231 | NA | NA | NA | NA | NA | N/S | N/S | organic |
| 337.238 | 154 | NA | NA | NA | NA | NA | N/S | minor decrease | organic |
| 371.327 | 167 | NA | NA | NA | NA | NA | N/S | major decrease | organic |
| 377.146 | 137 | Riboflavin | 0.73 | 7 | 0 | 10.34 | N/S | major decrease | organic |
| 400.343 | 209 | Spectral Match to Palmitoylcarnitine | 0.91 | 12 | 0 | 6.74 | N/S | major increase | organic |
| 401.347 | 189 | NA | NA | NA | NA | NA | N/S | major increase | organic |
| 401.355 | 255 | NA | NA | NA | NA | NA | N/S | minor increase | organic |
| 496.341 | 201 | NA | NA | NA | NA | NA | N/S | major increase | organic |
| 511.521 | 218 | NA | NA | NA | NA | NA | N/S | major decrease | organic |
| 538.520 | 362 | NA | NA | NA | NA | NA | N/S | minor increase | organic |
| 566.274 | 152 | NA | NA | NA | NA | NA | N/S | major increase | organic |
| 670.406 | 190 | NA | NA | NA | NA | NA | N/S | minor increase | organic |
| 731.588 | 351 | NA | NA | NA | NA | NA | N/S | N/S | organic |
| 768.554 | 519 | NA | NA | NA | NA | NA | N/S | major decrease | organic |
| 796.612 | 420 | NA | NA | NA | NA | NA | major decrease | N/S | organic |
| 802.574 | 299 | NA | NA | NA | NA | NA | N/S | minor increase | organic |
| 809.563 | 308 | NA | NA | NA | NA | NA | N/S | major increase | organic |
| 850.632 | 417 | NA | NA | NA | NA | NA | N/S | minor decrease | organic |
| 1039.673 | 175 | NA | NA | NA | NA | NA | N/S | N/S | organic |

| | | | | | | | | | |
|---|---|---|---|---|---|---|---|---|---|
| 1313.977 | 336 | NA | NA | NA | NA | NA | N/S | N/S | organic |
| 1408.040 | 303 | NA | NA | NA | NA | NA | N/S | major increase | organic |
| **Top-ranking metabolites for CL and Sylvio X10/4 *T.cruzi* strains that met the random forest cut-off criteria** | | | | | | | | | |
| ***m/z*** | **RT (sec)** | **Annotation** | **Cosine Score** | **No. of Shared Peaks** | **Mass Diff. to Library Reference** | **ppm error** | **Impact of Infection at position A (Sylvio X10/4 vs uninfected 147 days)** | **Impact of Infection at position A (CL vs uninfected 90 days)** | **Extract** |
| 276.181 | 104 | NA | NA | NA | NA | NA | major increase | N/S | aqueous |
| 268.105 | 35 | NA | NA | NA | NA | NA | N/S | major decrease | organic |
| 162.114 | 23 | NA | NA | NA | NA | NA | N/S | N/S | aqueous |
| 210.063 | 32 | NA | NA | NA | NA | NA | major decrease | N/S | aqueous |
| **Top-ranking metabolites for Sylvio X10/4 *T.cruzi* strain that met the random forest cut-off criteria** | | | | | | | | | |
| ***m/z*** | **RT (sec)** | **Annotation** | **Cosine Score** | **No. of Shared Peaks** | **Mass Diff. to Library Reference** | **ppm error** | **Impact of Infection at position A (Sylvio X10/4 vs uninfected 147 days)** | **Impact of Infection at position A (CL vs uninfected 90 days)** | **Extract** |
| 147.077 | 24 | NA | NA | NA | NA | NA | N/S | N/S | aqueous |
| 148.061 | 27 | NA | NA | NA | NA | NA | minor increase | N/S | aqueous |
| 189.124 | 74 | NA | NA | NA | NA | NA | major increase | N/S | aqueous |
| 199.146 | 69 | NA | NA | NA | NA | NA | minor increase | N/S | aqueous |
| 200.069 | 18 | NA | NA | NA | NA | NA | minor increase | N/S | aqueous |
| 214.158 | 186 | NA | NA | NA | NA | NA | N/S | N/S | aqueous |
| 248.150 | 30 | C4:0-OH acylcarnitine[1] | NA | NA | NA | -1.39 | major increase | N/S | aqueous |
| 249.222 | 217 | NA | NA | NA | NA | NA | minor decrease | N/S | aqueous |
| 251.238 | 223 | NA | NA | NA | NA | NA | major decrease | N/S | aqueous |
| 260.187 | 148 | C6:0 acylcarnitine[1] | NA | NA | NA | 1.03 | major increase | N/S | aqueous |
| 274.184 | 166 | NA | NA | NA | NA | NA | major increase | N/S | aqueous |
| 277.216 | 240 | NA | NA | NA | NA | NA | minor decrease | N/S | aqueous |
| 277.217 | 182 | NA | NA | NA | NA | NA | minor decrease | N/S | aqueous |
| 299.201 | 171 | NA | NA | NA | NA | NA | major decrease | N/S | aqueous |

| | | | | | | | | | |
|---|---|---|---|---|---|---|---|---|---|
| 300.085 | 58 | NA | NA | NA | NA | NA | major increase | N/S | aqueous |
| 302.197 | 145 | NA | NA | NA | NA | NA | major increase | N/S | aqueous |
| 304.213 | 149 | CAR (8:1(OH))[1] | NA | NA | NA | 0.17 | major increase | N/S | aqueous |
| 307.213 | 245 | NA | NA | NA | NA | NA | major increase | N/S | aqueous |
| 317.210 | 217 | NA | NA | NA | NA | NA | minor decrease | N/S | aqueous |
| 329.221 | 189 | NA | NA | NA | NA | NA | major increase | N/S | aqueous |
| 332.244 | 158 | C10:0-OH acylcarnitine[1] | NA | NA | NA | -0.74 | major increase | N/S | aqueous |
| 342.264 | 227 | C12:1 acylcarnitine[1] | NA | NA | NA | -2.87 | N/S | N/S | aqueous |
| 384.179 | 139 | NA | NA | NA | NA | NA | minor increase | N/S | aqueous |
| 385.168 | 243 | NA | NA | NA | NA | NA | N/S | N/S | aqueous |
| 394.162 | 139 | NA | NA | NA | NA | NA | major increase | N/S | aqueous |
| 398.195 | 143 | NA | NA | NA | NA | NA | minor increase | N/S | aqueous |
| 400.160 | 29 | NA | NA | NA | NA | NA | N/S | N/S | aqueous |
| 412.210 | 147 | PC family member[1] | NA | NA | NA | NA | minor increase | N/S | aqueous |
| 414.322 | 186 | C16:1-OH acylcarnitine[1] | NA | NA | NA | -1.20 | major increase | N/S | aqueous |
| 416.338 | 192 | C16:0-OH acylcarnitine[1] | NA | NA | NA | -0.35 | major increase | N/S | aqueous |
| 419.316 | 295 | NA | NA | NA | NA | NA | N/S | N/S | aqueous |
| 420.144 | 139 | NA | NA | NA | NA | NA | major increase | N/S | aqueous |
| 422.159 | 119 | NA | NA | NA | NA | NA | major increase | N/S | aqueous |
| 422.195 | 146 | NA | NA | NA | NA | NA | minor increase | minor decrease | aqueous |
| 428.169 | 141 | NA | NA | NA | NA | NA | N/S | N/S | aqueous |
| 433.272 | 141 | NA | NA | NA | NA | NA | N/S | N/S | aqueous |
| 440.338 | 190 | C18:2-OH acylcarnitine[1] | NA | NA | NA | -0.33 | major increase | N/S | aqueous |
| 442.353 | 197 | C18:1-OH acylcarnitine[1] | NA | NA | NA | -1.80 | major increase | N/S | aqueous |
| 444.369 | 204 | C18:0-OH acylcarnitine[1] | NA | NA | NA | -1.01 | major increase | N/S | aqueous |
| 462.191 | 140 | NA | NA | NA | NA | NA | major increase | N/S | aqueous |
| 465.210 | 148 | NA | NA | NA | NA | NA | major increase | N/S | aqueous |
| 478.151 | 139 | NA | NA | NA | NA | NA | major increase | N/S | aqueous |
| 478.168 | 143 | NA | NA | NA | NA | NA | major increase | N/S | aqueous |
| 488.207 | 148 | NA | NA | NA | NA | NA | N/S | N/S | aqueous |
| 492.188 | 136 | NA | NA | NA | NA | NA | minor increase | N/S | aqueous |
| 492.200 | 144 | NA | NA | NA | NA | NA | minor increase | N/S | aqueous |

| | | | | | | | | | |
|---|---|---|---|---|---|---|---|---|---|
| 494.181 | 139 | NA | NA | NA | NA | NA | major increase | N/S | aqueous |
| 494.215 | 147 | NA | NA | NA | NA | NA | minor increase | N/S | aqueous |
| 508.376 | 215 | LPC (O-18:1)[1] | NA | NA | NA | -2.46 | minor decrease | N/S | aqueous |
| 531.409 | 299 | NA | NA | NA | NA | NA | N/S | N/S | aqueous |
| 536.185 | 38 | NA | NA | NA | NA | NA | N/S | major decrease | aqueous |
| 575.414 | 235 | NA | NA | NA | NA | NA | N/S | N/S | aqueous |
| 576.294 | 163 | NA | NA | NA | NA | NA | minor increase | N/S | aqueous |
| 578.343 | 178 | NA | NA | NA | NA | NA | minor increase | N/S | aqueous |
| 600.330 | 174 | NA | NA | NA | NA | NA | N/S | N/S | aqueous |
| 604.329 | 174 | NA | NA | NA | NA | NA | major increase | N/S | aqueous |
| 749.424 | 194 | NA | NA | NA | NA | NA | major increase | N/S | aqueous |
| 816.575 | 273 | NA | NA | NA | NA | NA | minor increase | N/S | aqueous |
| 213.210 | 42 | NA | NA | NA | NA | NA | major increase | N/S | organic |
| 220.099 | 143 | NA | NA | NA | NA | NA | N/S | N/S | organic |
| 269.093 | 43 | NA | NA | NA | NA | NA | major decrease | N/S | organic |
| 291.611 | 172 | NA | NA | NA | NA | NA | N/S | N/S | organic |
| 300.291 | 210 | NA | NA | NA | NA | NA | N/S | N/S | organic |
| 302.207 | 157 | NA | NA | NA | NA | NA | major increase | N/S | organic |
| 302.215 | 161 | NA | NA | NA | NA | NA | major increase | N/S | organic |
| 304.213 | 141 | CAR (8:1(OH))[1] | NA | NA | NA | 0.17 | N/S | N/S | organic |
| 321.278 | 162 | NA | NA | NA | NA | NA | N/S | N/S | organic |
| 345.265 | 195 | NA | NA | NA | NA | NA | minor increase | N/S | organic |
| 358.260 | 150 | NA | NA | NA | NA | NA | major increase | N/S | organic |
| 360.275 | 153 | C12:0-OH acylcarnitine[1] | NA | NA | NA | -1.52 | major increase | N/S | organic |
| 365.342 | 253 | NA | NA | NA | NA | NA | minor decrease | N/S | organic |
| 388.307 | 160 | C14:0-OH acylcarnitine[1] | NA | NA | NA | 0.39 | major increase | N/S | organic |
| 398.328 | 179 | 9-Hexadecenoylcarnitine (Hexadecenoyl-L-carnitine)[1] | NA | NA | NA | 1.05 | major increase | N/S | organic |
| 414.359 | 193 | C17:0 acylcarnitine[1] | NA | NA | NA | 0.28 | N/S | N/S | organic |
| 416.338 | 171 | C16:0-OH acylcarnitine[1] | NA | NA | NA | -0.35 | major increase | N/S | organic |
| 432.239 | 168 | NA | NA | NA | NA | NA | minor decrease | N/S | organic |
| 442.354 | 175 | C18:1-OH acylcarnitine[1] | NA | NA | NA | 0.46 | major increase | N/S | organic |

| | | | | | | | | | |
|---|---|---|---|---|---|---|---|---|---|
| 444.369 | 185 | C18:0-OH acylcarnitine[1] | NA | NA | NA | -1.01 | major increase | N/S | organic |
| 454.389 | 250 | C20:1 acylcarnitine[1] | NA | NA | NA | 2.60 | major increase | N/S | organic |
| 456.406 | 229 | C20:0 acylcarnitine[1] | NA | NA | NA | 0.37 | N/S | N/S | organic |
| 470.385 | 189 | C20:1-OH acylcarnitine[1] | NA | NA | NA | -0.21 | major increase | N/S | organic |
| 482.420 | 231 | C22:1 acylcarnitine[1] | NA | NA | NA | -3.07 | major increase | N/S | organic |
| 484.364 | 175 | NA | NA | NA | NA | NA | major increase | N/S | organic |
| 508.377 | 187 | LPC (O-18:1)[1] | NA | NA | NA | -0.49 | major decrease | N/S | organic |
| 508.583 | 414 | NA | NA | NA | NA | NA | N/S | N/S | organic |
| 526.280 | 164 | NA | NA | NA | NA | NA | major decrease | N/S | organic |
| 530.324 | 184 | NA | NA | NA | NA | NA | N/S | N/S | organic |
| 561.397 | 200 | NA | NA | NA | NA | NA | N/S | N/S | organic |
| 562.389 | 219 | NA | NA | NA | NA | NA | major decrease | N/S | organic |
| 566.305 | 148 | NA | NA | NA | NA | NA | N/S | N/S | organic |
| 568.567 | 394 | NA | NA | NA | NA | NA | N/S | N/S | organic |
| 570.349 | 202 | NA | NA | NA | NA | NA | minor decrease | N/S | organic |
| 592.333 | 143 | NA | NA | NA | NA | NA | N/S | N/S | organic |
| 594.656 | 454 | NA | NA | NA | NA | NA | major increase | N/S | organic |
| 600.468 | 199 | NA | NA | NA | NA | NA | N/S | N/S | organic |
| 606.377 | 191 | NA | NA | NA | NA | NA | N/S | N/S | organic |
| 629.356 | 204 | NA | NA | NA | NA | NA | N/S | N/S | organic |
| 718.573 | 354 | NA | NA | NA | NA | NA | major decrease | N/S | organic |
| 720.590 | 455 | PC family member[1] | NA | NA | NA | NA | minor decrease | N/S | organic |
| 726.542 | 400 | NA | NA | NA | NA | NA | minor decrease | N/S | organic |
| 726.542 | 521 | NA | NA | NA | NA | NA | minor decrease | N/S | organic |
| 746.603 | 523 | Spectral Match to 1-Hexadecyl-2-(9Z-octadecenoyl)-sn-glycero-3-phosphocholine | 0.84 | 17 | 0 | 8.57 | N/S | N/S | organic |
| 746.605 | 457 | Spectral Match to 1-Hexadecyl-2-(9Z-octadecenoyl)-sn-glycero-3-phosphocholine | 0.84 | 17 | 0 | 8.57 | minor decrease | N/S | organic |
| 747.609 | 420 | NA | NA | NA | NA | NA | minor decrease | minor increase | organic |

| | | | | | | | | | |
|---|---|---|---|---|---|---|---|---|---|
| 770.591 | 321 | NA | NA | NA | NA | NA | minor decrease | N/S | organic |
| 789.527 | 334 | NA | NA | NA | NA | NA | N/S | N/S | organic |
| 796.615 | 397 | NA | NA | NA | NA | NA | major decrease | N/S | organic |

[1]Annotated based on molecular networking to an annotated sub-network; NA: not applicable; N/S: not statistically significant

**S2 Table. Annotated metabolites of combined extracts perturbed by infection at position B, identified through random forest classifier.**

| Top-ranking metabolites for CL *T.cruzi* strain that met the random forest cut-off criteria | | | | | | | | | |
|---|---|---|---|---|---|---|---|---|---|
| ***m/z*** | **RT (sec)** | **Annotation** | **Cosine Score** | **No. of Shared Peaks** | **Mass Diff. to Library Reference** | **ppm error** | **Impact of Infection at position B (Sylvio X10/4 vs uninfected 147 days)** | **Impact of Infection at position B (CL vs uninfected 90 days)** | **Extract** |
| 154.059 | 19 | NA | NA | NA | NA | NA | N/S | major increase | aqueous |
| 163.979 | 21 | NA | NA | NA | NA | NA | N/S | minor increase | aqueous |
| 189.124 | 74 | NA | NA | NA | NA | NA | N/S | minor decrease | aqueous |
| 193.610 | 145 | NA | NA | NA | NA | NA | N/S | major decrease | aqueous |
| 241.155 | 26 | NA | NA | NA | NA | NA | N/S | minor decrease | aqueous |
| 252.233 | 190 | NA | NA | NA | NA | NA | N/S | major decrease | aqueous |
| 270.244 | 200 | NA | NA | NA | NA | NA | N/S | major decrease | aqueous |
| 280.264 | 206 | NA | NA | NA | NA | NA | N/S | major decrease | aqueous |
| 296.259 | 187 | NA | NA | NA | NA | NA | N/S | major decrease | aqueous |
| 305.209 | 232 | NA | NA | NA | NA | NA | N/S | N/S | aqueous |
| 316.285 | 191 | NA | NA | NA | NA | NA | N/S | major decrease | aqueous |
| 320.256 | 206 | NA | NA | NA | NA | NA | N/S | major decrease | aqueous |
| 332.280 | 175 | NA | NA | NA | NA | NA | N/S | major decrease | aqueous |
| 336.251 | 190 | NA | NA | NA | NA | NA | N/S | major decrease | aqueous |
| 346.348 | 217 | NA | NA | NA | NA | NA | N/S | N/S | aqueous |
| 368.201 | 147 | NA | NA | NA | NA | NA | N/S | major decrease | aqueous |
| 382.296 | 182 | C15:2 acylcarnitine[1] | NA | NA | NA | -0.74 | N/S | minor increase | aqueous |
| 438.228 | 148 | NA | NA | NA | NA | NA | N/S | minor increase | aqueous |
| 446.196 | 147 | NA | NA | NA | NA | NA | N/S | minor decrease | aqueous |
| 462.191 | 140 | NA | NA | NA | NA | NA | N/S | minor decrease | aqueous |
| 556.443 | 228 | NA | NA | NA | NA | NA | N/S | major decrease | aqueous |
| 560.334 | 180 | NA | NA | NA | NA | NA | N/S | minor increase | aqueous |
| 588.111 | 138 | NA | NA | NA | NA | NA | N/S | major decrease | aqueous |
| 605.357 | 169 | NA | NA | NA | NA | NA | N/S | major decrease | aqueous |
| 1015.673 | 207 | NA | NA | NA | NA | NA | N/S | N/S | aqueous |

| | | | | | | | | | |
|---|---|---|---|---|---|---|---|---|---|
| 1039.673 | 203 | NA | NA | NA | NA | NA | N/S | N/S | aqueous |
| 159.027 | 20 | NA | NA | NA | NA | NA | N/S | minor increase | organic |
| 228.196 | 177 | NA | NA | NA | NA | NA | N/S | minor decrease | organic |
| 230.176 | 150 | NA | NA | NA | NA | NA | N/S | major decrease | organic |
| 241.156 | 20 | NA | NA | NA | NA | NA | N/S | major decrease | organic |
| 285.219 | 135 | NA | NA | NA | NA | NA | N/S | minor increase | organic |
| 296.259 | 181 | NA | NA | NA | NA | NA | N/S | major decrease | organic |
| 339.075 | 146 | NA | NA | NA | NA | NA | N/S | minor increase | organic |
| 381.263 | 212 | NA | NA | NA | NA | NA | N/S | minor increase | organic |
| 415.213 | 168 | NA | NA | NA | NA | NA | N/S | major increase | organic |
| 440.240 | 144 | NA | NA | NA | NA | NA | N/S | major decrease | organic |
| 466.292 | 159 | NA | NA | NA | NA | NA | N/S | minor increase | organic |
| 494.568 | 366 | NA | NA | NA | NA | NA | N/S | major increase | organic |
| 496.422 | 226 | NA | NA | NA | NA | NA | N/S | major increase | organic |
| 534.561 | 450 | NA | NA | NA | NA | NA | N/S | major decrease | organic |
| 608.635 | 474 | NA | NA | NA | NA | NA | N/S | minor increase | organic |
| 692.421 | 143 | PC family member[1] | NA | NA | NA | NA | N/S | N/S | organic |
| **Top-ranking metabolites for CL and Sylvio X10/4 *T.cruzi* strains that met the random forest cut-off criteria** | | | | | | | | | |
| ***m/z*** | **RT (sec)** | **Annotation** | **Cosine Score** | **No. of Shared Peaks** | **Mass Diff. to Library Reference** | **ppm error** | **Impact of Infection at position B (Sylvio X10/4 vs uninfected 147 days)** | **Impact of Infection at position B (CL vs uninfected 90 days)** | **Extract** |
| 398.195 | 143 | NA | NA | NA | NA | NA | minor increase | minor decrease | aqueous |
| **Top-ranking metabolites for Sylvio X10/4 *T.cruzi* strain that met the random forest cut-off criteria** | | | | | | | | | |
| ***m/z*** | **RT (sec)** | **Annotation** | **Cosine Score** | **No. of Shared Peaks** | **Mass Diff. to Library Reference** | **ppm error** | **Impact of Infection at position B (Sylvio X10/4 vs uninfected 147 days)** | **Impact of Infection at position B (CL vs uninfected 90 days)** | **Extract** |
| 147.114 | 14 | NA | NA | NA | NA | NA | N/S | N/S | aqueous |
| 149.024 | 171 | NA | NA | NA | NA | NA | major decrease | N/S | aqueous |
| 152.058 | 41 | NA | NA | NA | NA | NA | N/S | N/S | aqueous |
| 153.042 | 32 | NA | NA | NA | NA | NA | minor decrease | N/S | aqueous |

| | | | | | | | | | |
|---|---|---|---|---|---|---|---|---|---|
| 185.596 | 143 | NA | NA | NA | NA | NA | major increase | major decrease | aqueous |
| 223.133 | 183 | NA | NA | NA | NA | NA | minor increase | N/S | aqueous |
| 244.155 | 140 | NA | NA | NA | NA | NA | major increase | N/S | aqueous |
| 248.150 | 30 | C4:0-OH acylcarnitine[1] | NA | NA | NA | -1.39 | major increase | N/S | aqueous |
| 248.151 | 47 | C4:0-OH acylcarnitine[1] | NA | NA | NA | 2.64 | major increase | N/S | aqueous |
| 256.229 | 193 | NA | NA | NA | NA | NA | N/S | N/S | aqueous |
| 258.106 | 140 | NA | NA | NA | NA | NA | N/S | N/S | aqueous |
| 260.187 | 148 | C6:0 acylcarnitine[1] | NA | NA | NA | 1.03 | major increase | N/S | aqueous |
| 264.170 | 136 | NA | NA | NA | NA | NA | minor decrease | N/S | aqueous |
| 275.202 | 206 | NA | NA | NA | NA | NA | major decrease | N/S | aqueous |
| 353.232 | 179 | NA | NA | NA | NA | NA | minor decrease | N/S | aqueous |
| 370.262 | 186 | NA | NA | NA | NA | NA | minor decrease | N/S | aqueous |
| 384.117 | 130 | NA | NA | NA | NA | NA | N/S | N/S | aqueous |
| 399.252 | 222 | NA | NA | NA | NA | NA | minor decrease | N/S | aqueous |
| 400.269 | 187 | C14:2-DC acylcarnitine[1] | NA | NA | NA | -3.65 | minor decrease | major increase | aqueous |
| 404.150 | 137 | NA | NA | NA | NA | NA | minor increase | N/S | aqueous |
| 407.242 | 190 | NA | NA | NA | NA | NA | minor decrease | minor increase | aqueous |
| 409.258 | 186 | NA | NA | NA | NA | NA | minor decrease | N/S | aqueous |
| 412.200 | 132 | NA | NA | NA | NA | NA | minor increase | minor decrease | aqueous |
| 412.307 | 179 | C16:2-OH acylcarnitine[1] | NA | NA | NA | 0.37 | major increase | N/S | aqueous |
| 414.322 | 186 | C16:1-OH acylcarnitine[1] | NA | NA | NA | -1.20 | major increase | N/S | aqueous |
| 421.221 | 177 | NA | NA | NA | NA | NA | minor decrease | N/S | aqueous |
| 421.234 | 221 | NA | NA | NA | NA | NA | minor decrease | N/S | aqueous |
| 424.269 | 185 | NA | NA | NA | NA | NA | major decrease | major increase | aqueous |
| 438.380 | 237 | NA | NA | NA | NA | NA | major decrease | N/S | aqueous |
| 442.353 | 197 | C18:1-OH acylcarnitine[1] | NA | NA | NA | -1.80 | major increase | N/S | aqueous |
| 462.191 | 148 | NA | NA | NA | NA | NA | N/S | major decrease | aqueous |
| 478.168 | 143 | NA | NA | NA | NA | NA | minor increase | N/S | aqueous |
| 534.319 | 181 | PC family member[1] | NA | NA | NA | NA | minor increase | N/S | aqueous |
| 568.325 | 190 | NA | NA | NA | NA | NA | minor increase | N/S | aqueous |
| 594.339 | 163 | NA | NA | NA | NA | NA | minor increase | N/S | aqueous |
| 645.469 | 224 | NA | NA | NA | NA | NA | minor increase | N/S | aqueous |

| | | | | | | | | | |
|---|---|---|---|---|---|---|---|---|---|
| 742.539 | 303 | PC family member[1] | NA | NA | NA | NA | minor increase | N/S | aqueous |
| 838.267 | 187 | NA | NA | NA | NA | NA | major increase | N/S | aqueous |
| 1019.705 | 225 | NA | NA | NA | NA | NA | N/S | N/S | aqueous |
| 302.215 | 161 | NA | NA | NA | NA | NA | major increase | N/S | organic |
| 368.280 | 158 | C14:2 acylcarnitine[1] | NA | NA | NA | -1.72 | major increase | N/S | organic |
| 370.296 | 166 | C14:1 acylcarnitine[1] | NA | NA | NA | -0.76 | major increase | N/S | organic |
| 388.307 | 160 | C14:0-OH acylcarnitine[1] | NA | NA | NA | 0.39 | major increase | N/S | organic |
| 438.323 | 163 | C18:3-OH acylcarnitine[1] | NA | NA | NA | 1.15 | major increase | N/S | organic |
| 573.401 | 248 | NA | NA | NA | NA | NA | major increase | N/S | organic |
| 636.388 | 185 | NA | NA | NA | NA | NA | major decrease | N/S | organic |
| 636.391 | 205 | NA | NA | NA | NA | NA | major decrease | N/S | organic |
| 734.450 | 189 | NA | NA | NA | NA | NA | minor decrease | N/S | organic |

[1]Annotated based on molecular networking to an annotated sub-network; NA: not applicable; N/S: not statistically significant

**S3 Table. Annotated metabolites of combined extracts perturbed by infection at position C, identified through random forest classifier.**

| Top-ranking metabolites for CL *T.cruzi* strain that met the random forest cut-off criteria | | | | | | | | | |
|---|---|---|---|---|---|---|---|---|---|
| ***m/z*** | **RT (sec)** | **Annotation** | **Cosine Score** | **No. of Shared Peaks** | **Mass Diff. to Library Reference** | **ppm error** | **Impact of Infection at position C (Sylvio X10/4 vs uninfected 147 days)** | **Impact of Infection at position C (CL vs uninfected 90 days)** | **Extract** |
| 150.059 | 27 | NA | NA | NA | NA | NA | minor decrease | minor decrease | aqueous |
| 155.071 | 123 | NA | NA | NA | NA | NA | N/S | minor increase | aqueous |
| 166.087 | 101 | NA | NA | NA | NA | NA | N/S | minor decrease | aqueous |
| 166.088 | 134 | NA | NA | NA | NA | NA | N/S | minor decrease | aqueous |
| 187.097 | 145 | NA | NA | NA | NA | NA | N/S | N/S | aqueous |
| 188.072 | 116 | NA | NA | NA | NA | NA | N/S | major decrease | aqueous |
| 189.124 | 74 | NA | NA | NA | NA | NA | N/S | minor decrease | aqueous |
| 199.146 | 69 | NA | NA | NA | NA | NA | N/S | minor increase | aqueous |
| 220.098 | 151 | NA | NA | NA | NA | NA | N/S | major increase | aqueous |
| 223.133 | 183 | NA | NA | NA | NA | NA | N/S | minor increase | aqueous |
| 223.097 | 184 | NA | NA | NA | NA | NA | N/S | minor increase | aqueous |
| 227.092 | 138 | NA | NA | NA | NA | NA | N/S | minor increase | aqueous |
| 241.155 | 26 | NA | NA | NA | NA | NA | N/S | minor decrease | aqueous |
| 252.233 | 190 | NA | NA | NA | NA | NA | N/S | major decrease | aqueous |
| 263.238 | 234 | NA | NA | NA | NA | NA | N/S | N/S | aqueous |
| 270.244 | 200 | NA | NA | NA | NA | NA | N/S | major decrease | aqueous |
| 278.248 | 179 | NA | NA | NA | NA | NA | N/S | major decrease | aqueous |
| 280.264 | 206 | NA | NA | NA | NA | NA | N/S | major decrease | aqueous |
| 293.212 | 227 | NA | NA | NA | NA | NA | N/S | major decrease | aqueous |
| 296.259 | 187 | NA | NA | NA | NA | NA | N/S | major decrease | aqueous |
| 312.254 | 194 | NA | NA | NA | NA | NA | N/S | major decrease | aqueous |
| 313.239 | 197 | NA | NA | NA | NA | NA | N/S | minor decrease | aqueous |
| 332.280 | 175 | NA | NA | NA | NA | NA | N/S | major decrease | aqueous |
| 336.251 | 190 | NA | NA | NA | NA | NA | N/S | major decrease | aqueous |
| 389.232 | 202 | NA | NA | NA | NA | NA | minor decrease | minor increase | aqueous |

| | | | | | | | | | |
|---|---|---|---|---|---|---|---|---|---|
| 393.263 | 193 | NA | NA | NA | NA | NA | N/S | minor increase | aqueous |
| 407.242 | 190 | NA | NA | NA | NA | NA | N/S | minor increase | aqueous |
| 421.221 | 177 | NA | NA | NA | NA | NA | N/S | minor increase | aqueous |
| 425.253 | 177 | NA | NA | NA | NA | NA | N/S | minor increase | aqueous |
| 454.294 | 206 | NA | NA | NA | NA | NA | N/S | minor increase | aqueous |
| 468.309 | 193 | LPC (14:0)[1] | NA | NA | NA | -1.20 | N/S | minor increase | aqueous |
| 482.325 | 234 | NA | NA | NA | NA | NA | N/S | major increase | aqueous |
| 494.288 | 186 | NA | NA | NA | NA | NA | N/S | minor increase | aqueous |
| 494.567 | 279 | NA | NA | NA | NA | NA | N/S | major increase | aqueous |
| 496.341 | 208 | NA | NA | NA | NA | NA | N/S | minor increase | aqueous |
| 512.334 | 180 | NA | NA | NA | NA | NA | N/S | minor increase | aqueous |
| 518.322 | 206 | LPC (16:0)+Na[1] | NA | NA | NA | -1.56 | N/S | minor increase | aqueous |
| 524.372 | 222 | LPC (18:0)[1] | NA | NA | NA | -0.31 | N/S | major increase | aqueous |
| 528.366 | 197 | NA | NA | NA | NA | NA | N/S | major increase | aqueous |
| 538.350 | 185 | NA | NA | NA | NA | NA | N/S | minor increase | aqueous |
| 572.370 | 214 | NA | NA | NA | NA | NA | N/S | minor decrease | aqueous |
| 675.544 | 270 | NA | NA | NA | NA | NA | N/S | minor increase | aqueous |
| 689.559 | 279 | NA | NA | NA | NA | NA | N/S | major increase | aqueous |
| 200.523 | 171 | NA | NA | NA | NA | NA | N/S | major increase | organic |
| 209.093 | 57 | NA | NA | NA | NA | NA | N/S | major increase | organic |
| 226.181 | 249 | NA | NA | NA | NA | NA | N/S | major decrease | organic |
| 246.171 | 138 | C5:0 acylcarnitine (valerylcarnitine)[1] | NA | NA | NA | -0.33 | N/S | N/S | organic |
| 258.206 | 162 | NA | NA | NA | NA | NA | N/S | major decrease | organic |
| 288.290 | 162 | NA | NA | NA | NA | NA | N/S | N/S | organic |
| 296.259 | 181 | NA | NA | NA | NA | NA | N/S | major decrease | organic |
| 300.280 | 148 | NA | NA | NA | NA | NA | N/S | major increase | organic |
| 304.301 | 177 | NA | NA | NA | NA | NA | N/S | major decrease | organic |
| 319.226 | 151 | NA | NA | NA | NA | NA | N/S | minor increase | organic |
| 326.379 | 206 | NA | NA | NA | NA | NA | major increase | major decrease | organic |
| 326.379 | 231 | NA | NA | NA | NA | NA | major increase | major decrease | organic |
| 332.331 | 219 | NA | NA | NA | NA | NA | N/S | major decrease | organic |

| | | | | | | | | | |
|---|---|---|---|---|---|---|---|---|---|
| 332.331 | 274 | NA | NA | NA | NA | NA | N/S | major decrease | organic |
| 335.221 | 170 | NA | NA | NA | NA | NA | N/S | minor increase | organic |
| 353.118 | 142 | NA | NA | NA | NA | NA | N/S | N/S | organic |
| 353.092 | 19 | NA | NA | NA | NA | NA | N/S | minor increase | organic |
| 354.338 | 235 | NA | NA | NA | NA | NA | N/S | major decrease | organic |
| 359.222 | 179 | NA | NA | NA | NA | NA | N/S | N/S | organic |
| 360.364 | 211 | NA | NA | NA | NA | NA | N/S | minor decrease | organic |
| 364.067 | 26 | NA | NA | NA | NA | NA | N/S | major increase | organic |
| 375.217 | 162 | NA | NA | NA | NA | NA | N/S | minor increase | organic |
| 411.309 | 200 | NA | NA | NA | NA | NA | N/S | N/S | organic |
| 428.037 | 26 | NA | NA | NA | NA | NA | N/S | major increase | organic |
| 436.174 | 140 | NA | NA | NA | NA | NA | N/S | N/S | organic |
| 439.101 | 136 | NA | NA | NA | NA | NA | N/S | major increase | organic |
| 502.294 | 172 | NA | NA | NA | NA | NA | N/S | minor decrease | organic |
| 518.377 | 307 | NA | NA | NA | NA | NA | N/S | major increase | organic |
| 528.307 | 179 | NA | NA | NA | NA | NA | N/S | minor decrease | organic |
| 530.290 | 179 | NA | NA | NA | NA | NA | N/S | minor decrease | organic |
| 530.324 | 184 | NA | NA | NA | NA | NA | N/S | minor decrease | organic |
| 538.387 | 204 | LPC (19:0)[1] | NA | NA | NA | -1.51 | N/S | major decrease | organic |
| 550.351 | 190 | NA | NA | NA | NA | NA | N/S | major decrease | organic |
| 550.387 | 199 | LPC (20:1)[1] | NA | NA | NA | -1.48 | N/S | minor decrease | organic |
| 552.403 | 215 | LPC (20:0)[1] | NA | NA | NA | -0.84 | major increase | major decrease | organic |
| 556.303 | 187 | NA | NA | NA | NA | NA | N/S | major decrease | organic |
| 558.337 | 194 | NA | NA | NA | NA | NA | N/S | N/S | organic |
| 560.332 | 161 | NA | NA | NA | NA | NA | N/S | minor increase | organic |
| 566.552 | 387 | NA | NA | NA | NA | NA | N/S | major increase | organic |
| 573.401 | 248 | NA | NA | NA | NA | NA | N/S | N/S | organic |
| 576.294 | 146 | NA | NA | NA | NA | NA | N/S | minor increase | organic |
| 576.572 | 412 | NA | NA | NA | NA | NA | N/S | major increase | organic |
| 582.355 | 142 | NA | NA | NA | NA | NA | N/S | minor increase | organic |
| 582.414 | 187 | NA | NA | NA | NA | NA | N/S | N/S | organic |
| 594.583 | 411 | NA | NA | NA | NA | NA | N/S | major increase | organic |

| | | | | | | | | | |
|---|---|---|---|---|---|---|---|---|---|
| 606.450 | 245 | NA | NA | NA | NA | NA | N/S | N/S | organic |
| 606.920 | 34 | NA | NA | NA | NA | NA | N/S | major increase | organic |
| 616.325 | 153 | NA | NA | NA | NA | NA | N/S | minor increase | organic |
| 618.340 | 146 | NA | NA | NA | NA | NA | N/S | minor increase | organic |
| 618.377 | 207 | NA | NA | NA | NA | NA | N/S | minor decrease | organic |
| 660.554 | 234 | NA | NA | NA | NA | NA | N/S | minor increase | organic |
| 663.454 | 424 | NA | NA | NA | NA | NA | N/S | major increase | organic |
| 672.423 | 204 | NA | NA | NA | NA | NA | N/S | minor decrease | organic |
| 672.424 | 233 | NA | NA | NA | NA | NA | N/S | minor decrease | organic |
| 719.570 | 296 | NA | NA | NA | NA | NA | N/S | N/S | organic |
| 722.496 | 425 | NA | NA | NA | NA | NA | N/S | major increase | organic |
| 748.618 | 349 | NA | NA | NA | NA | NA | N/S | minor decrease | organic |
| 754.502 | 251 | NA | NA | NA | NA | NA | N/S | minor decrease | organic |
| 756.480 | 229 | NA | NA | NA | NA | NA | N/S | minor decrease | organic |
| 782.542 | 302 | NA | NA | NA | NA | NA | N/S | major decrease | organic |
| 782.570 | 336 | NA | NA | NA | NA | NA | N/S | minor increase | organic |
| 784.583 | 346 | PC (34:0)+Na[1] | NA | NA | NA | -0.98 | N/S | minor increase | organic |
| 788.547 | 391 | NA | NA | NA | NA | NA | N/S | N/S | organic |
| 788.581 | 294 | NA | NA | NA | NA | NA | major increase | N/S | organic |
| 798.531 | 255 | NA | NA | NA | NA | NA | N/S | N/S | organic |
| 810.599 | 274 | PC (36:1)+Na[1] | NA | NA | NA | -0.52 | N/S | minor increase | organic |
| 811.605 | 276 | NA | NA | NA | NA | NA | N/S | minor increase | organic |
| 812.516 | 244 | NA | NA | NA | NA | NA | N/S | minor decrease | organic |
| 820.521 | 257 | NA | NA | NA | NA | NA | N/S | minor decrease | organic |
| 828.525 | 236 | NA | NA | NA | NA | NA | N/S | N/S | organic |
| 834.601 | 390 | NA | NA | NA | NA | NA | N/S | minor increase | organic |
| 842.589 | 305 | NA | NA | NA | NA | NA | N/S | N/S | organic |
| 850.632 | 417 | NA | NA | NA | NA | NA | N/S | minor decrease | organic |
| 858.552 | 248 | NA | NA | NA | NA | NA | N/S | N/S | organic |
| 862.559 | 261 | NA | NA | NA | NA | NA | major increase | minor decrease | organic |
| 864.573 | 282 | NA | NA | NA | NA | NA | N/S | minor decrease | organic |
| 870.696 | 349 | NA | NA | NA | NA | NA | N/S | major increase | organic |

| | | | | | | | | | |
|---|---|---|---|---|---|---|---|---|---|
| 878.545 | 234 | NA | NA | NA | NA | NA | N/S | minor decrease | organic |
| 890.558 | 238 | NA | NA | NA | NA | NA | N/S | minor decrease | organic |
| 894.549 | 234 | NA | NA | NA | NA | NA | N/S | minor decrease | organic |
| 896.564 | 250 | NA | NA | NA | NA | NA | N/S | minor decrease | organic |
| 906.569 | 244 | NA | NA | NA | NA | NA | minor increase | minor decrease | organic |
| 932.583 | 248 | NA | NA | NA | NA | NA | N/S | minor decrease | organic |
| 1027.579 | 172 | NA | NA | NA | NA | NA | N/S | major decrease | organic |
| 1312.972 | 335 | NA | NA | NA | NA | NA | N/S | major increase | organic |
| **Top-ranking metabolites for CL and Sylvio X10/4 *T.cruzi* strains that met the random forest cut-off criteria** | | | | | | | | | |
| ***m/z*** | **RT (sec)** | **Annotation** | **Cosine Score** | **No. of Shared Peaks** | **Mass Diff. to Library Reference** | **ppm error** | **Impact of Infection at position C (Sylvio X10/4 vs uninfected 147 days)** | **Impact of Infection at position C (CL vs uninfected 90 days)** | **Extract** |
| 209.093 | 53 | NA | NA | NA | NA | NA | major decrease | major increase | aqueous |
| 320.256 | 217 | NA | NA | NA | NA | NA | major increase | major decrease | aqueous |
| 375.216 | 186 | NA | NA | NA | NA | NA | minor decrease | minor increase | aqueous |
| 787.535 | 427 | NA | NA | NA | NA | NA | N/S | major increase | organic |
| **Top-ranking metabolites for Sylvio X10/4 *T.cruzi* strain that met the random forest cut-off criteria** | | | | | | | | | |
| ***m/z*** | **RT (sec)** | **Annotation** | **Cosine Score** | **No. of Shared Peaks** | **Mass Diff. to Library Reference** | **ppm error** | **Impact of Infection at position C (Sylvio X10/4 vs uninfected 147 days)** | **Impact of Infection at position C (CL vs uninfected 90 days)** | **Extract** |
| 147.114 | 14 | NA | NA | NA | NA | NA | minor decrease | N/S | aqueous |
| 175.120 | 19 | NA | NA | NA | NA | NA | minor decrease | N/S | aqueous |
| 193.610 | 145 | NA | NA | NA | NA | NA | major increase | N/S | aqueous |
| 220.136 | 160 | NA | NA | NA | NA | NA | major increase | N/S | aqueous |
| 235.119 | 140 | NA | NA | NA | NA | NA | N/S | N/S | aqueous |
| 248.132 | 152 | NA | NA | NA | NA | NA | minor decrease | N/S | aqueous |
| 248.150 | 30 | C4:0-OH acylcarnitine[1] | NA | NA | NA | -1.39 | major increase | N/S | aqueous |
| 253.217 | 203 | NA | NA | NA | NA | NA | N/S | N/S | aqueous |
| 257.101 | 143 | NA | NA | NA | NA | NA | minor decrease | N/S | aqueous |

| | | | | | | | | | |
|---|---|---|---|---|---|---|---|---|---|
| 258.112 | 146 | sn-glycero-3-phosphocholine[1] | NA | NA | NA | 3.11 | major decrease | N/S | aqueous |
| 258.171 | 145 | C6:1 acylcarnitine[1] | NA | NA | NA | -0.32 | major increase | N/S | aqueous |
| 260.187 | 148 | C6:0 acylcarnitine[1] | NA | NA | NA | 1.03 | major increase | N/S | aqueous |
| 262.166 | 43 | CAR(5:0(OH))[1] | NA | NA | NA | 0.02 | minor increase | N/S | aqueous |
| 268.108 | 33 | NA | NA | NA | NA | NA | major increase | N/S | aqueous |
| 274.184 | 166 | NA | NA | NA | NA | NA | major increase | N/S | aqueous |
| 285.290 | 177 | NA | NA | NA | NA | NA | N/S | N/S | aqueous |
| 287.078 | 94 | NA | NA | NA | NA | NA | N/S | N/S | aqueous |
| 291.072 | 17 | NA | NA | NA | NA | NA | N/S | N/S | aqueous |
| 296.067 | 18 | NA | NA | NA | NA | NA | minor decrease | N/S | aqueous |
| 302.216 | 178 | NA | NA | NA | NA | NA | major increase | N/S | aqueous |
| 326.208 | 137 | NA | NA | NA | NA | NA | N/S | N/S | aqueous |
| 329.221 | 189 | NA | NA | NA | NA | NA | N/S | N/S | aqueous |
| 330.265 | 197 | C11:0 acylcarnitine[1] | NA | NA | NA | 0.05 | minor increase | N/S | aqueous |
| 332.244 | 158 | C10:0-OH acylcarnitine[1] | NA | NA | NA | -0.74 | major increase | N/S | aqueous |
| 339.346 | 275 | NA | NA | NA | NA | NA | N/S | major increase | aqueous |
| 346.259 | 182 | NA | NA | NA | NA | NA | major increase | N/S | aqueous |
| 346.259 | 187 | NA | NA | NA | NA | NA | major increase | N/S | aqueous |
| 348.071 | 29 | NA | NA | NA | NA | NA | major increase | N/S | aqueous |
| 358.260 | 164 | NA | NA | NA | NA | NA | major increase | N/S | aqueous |
| 359.235 | 204 | NA | NA | NA | NA | NA | minor decrease | N/S | aqueous |
| 360.275 | 170 | C12:0-OH acylcarnitine[1] | NA | NA | NA | -1.52 | major increase | N/S | aqueous |
| 361.278 | 174 | NA | NA | NA | NA | NA | major increase | N/S | aqueous |
| 368.279 | 215 | NA | NA | NA | NA | NA | major increase | N/S | aqueous |
| 370.053 | 26 | NA | NA | NA | NA | NA | major increase | N/S | aqueous |
| 386.211 | 146 | NA | NA | NA | NA | NA | major increase | N/S | aqueous |
| 398.195 | 143 | NA | NA | NA | NA | NA | minor increase | N/S | aqueous |
| 398.327 | 198 | 9-Hexadecenoylcarnitine (Hexadecenoyl-L-carnitine)[1] | NA | NA | NA | -1.46 | major increase | N/S | aqueous |
| 400.343 | 205 | Spectral Match to Palmitoylcarnitine | 0.91 | 11 | 0 | 2 | major increase | major increase | aqueous |

| | | | | | | | | | |
|---|---|---|---|---|---|---|---|---|---|
| 410.290 | 173 | NA | NA | NA | NA | NA | N/S | N/S | aqueous |
| 412.200 | 132 | NA | NA | NA | NA | NA | minor increase | N/S | aqueous |
| 412.210 | 147 | PC family member[1] | NA | NA | NA | NA | minor increase | N/S | aqueous |
| 414.226 | 148 | NA | NA | NA | NA | NA | N/S | N/S | aqueous |
| 414.322 | 186 | C16:1-OH acylcarnitine[1] | NA | NA | NA | -1.20 | major increase | N/S | aqueous |
| 416.153 | 33 | NA | NA | NA | NA | NA | major decrease | N/S | aqueous |
| 416.338 | 192 | C16:0-OH acylcarnitine[1] | NA | NA | NA | -0.35 | major increase | N/S | aqueous |
| 424.269 | 185 | NA | NA | NA | NA | NA | N/S | N/S | aqueous |
| 424.343 | 203 | C18:2 acylcarnitine[1] | NA | NA | NA | -0.55 | major increase | major increase | aqueous |
| 426.359 | 210 | Spectral Match to Oleoyl-L-carnitine | 0.85 | 10 | 0 | 0 | major increase | major increase | aqueous |
| 428.374 | 217 | C18:0 acylcarnitine[1] | NA | NA | NA | -1.24 | major increase | N/S | aqueous |
| 438.228 | 148 | NA | NA | NA | NA | NA | N/S | N/S | aqueous |
| 438.299 | 210 | NA | NA | NA | NA | NA | minor decrease | N/S | aqueous |
| 440.338 | 190 | C18:2-OH acylcarnitine[1] | NA | NA | NA | -0.33 | major increase | major increase | aqueous |
| 444.369 | 204 | C18:0-OH acylcarnitine[1] | NA | NA | NA | -1.01 | major increase | major increase | aqueous |
| 462.191 | 140 | NA | NA | NA | NA | NA | minor increase | N/S | aqueous |
| 462.191 | 148 | NA | NA | NA | NA | NA | major increase | N/S | aqueous |
| 464.206 | 147 | NA | NA | NA | NA | NA | major increase | major decrease | aqueous |
| 468.368 | 198 | C20:2-OH acylcarnitine[1] | NA | NA | NA | -3.09 | major increase | N/S | aqueous |
| 502.223 | 155 | NA | NA | NA | NA | NA | minor increase | N/S | aqueous |
| 508.303 | 174 | NA | NA | NA | NA | NA | minor decrease | N/S | aqueous |
| 508.340 | 212 | NA | NA | NA | NA | NA | minor decrease | N/S | aqueous |
| 521.344 | 219 | NA | NA | NA | NA | NA | minor increase | N/S | aqueous |
| 534.474 | 217 | NA | NA | NA | NA | NA | N/S | N/S | aqueous |
| 536.185 | 38 | NA | NA | NA | NA | NA | major increase | N/S | aqueous |
| 548.371 | 216 | LPC (20:2)[1] | NA | NA | NA | -2.12 | minor decrease | minor increase | aqueous |
| 570.340 | 155 | Lysophosphatidylcholine (22:4)[1] | NA | NA | NA | -28.95 | minor increase | N/S | aqueous |
| 570.341 | 173 | NA | NA | NA | NA | NA | minor increase | minor increase | aqueous |
| 584.335 | 182 | NA | NA | NA | NA | NA | minor decrease | N/S | aqueous |
| 634.249 | 187 | NA | NA | NA | NA | NA | N/S | N/S | aqueous |
| 646.408 | 222 | NA | NA | NA | NA | NA | N/S | N/S | aqueous |

| | | | | | | | | | |
|---|---|---|---|---|---|---|---|---|---|
| 688.492 | 285 | NA | NA | NA | NA | NA | N/S | N/S | aqueous |
| 716.522 | 299 | NA | NA | NA | NA | NA | minor increase | N/S | aqueous |
| 1135.509 | 200 | NA | NA | NA | NA | NA | N/S | N/S | aqueous |
| 137.526 | 22 | NA | NA | NA | NA | NA | N/S | N/S | organic |
| 184.014 | 20 | NA | NA | NA | NA | NA | N/S | N/S | organic |
| 216.197 | 150 | NA | NA | NA | NA | NA | major decrease | N/S | organic |
| 226.157 | 42 | NA | NA | NA | NA | NA | major decrease | N/S | organic |
| 228.196 | 177 | NA | NA | NA | NA | NA | major decrease | N/S | organic |
| 238.024 | 21 | NA | NA | NA | NA | NA | minor increase | N/S | organic |
| 248.151 | 19 | C4:0-OH acylcarnitine[1] | NA | NA | NA | 2.64 | major increase | N/S | organic |
| 268.264 | 216 | NA | NA | NA | NA | NA | minor decrease | N/S | organic |
| 274.183 | 153 | NA | NA | NA | NA | NA | major increase | N/S | organic |
| 279.095 | 159 | NA | NA | NA | NA | NA | major decrease | N/S | organic |
| 291.611 | 172 | NA | NA | NA | NA | NA | N/S | N/S | organic |
| 302.207 | 157 | NA | NA | NA | NA | NA | major increase | N/S | organic |
| 332.244 | 146 | C10:0-OH acylcarnitine[1] | NA | NA | NA | -0.74 | major increase | N/S | organic |
| 340.323 | 274 | NA | NA | NA | NA | NA | minor increase | N/S | organic |
| 360.275 | 153 | C12:0-OH acylcarnitine[1] | NA | NA | NA | -1.52 | major increase | N/S | organic |
| 370.296 | 166 | C14:1 acylcarnitine[1] | NA | NA | NA | -0.76 | major increase | N/S | organic |
| 371.053 | 40 | NA | NA | NA | NA | NA | N/S | N/S | organic |
| 386.291 | 157 | C14:1 acylcarnitine[1] | NA | NA | NA | -0.76 | major increase | N/S | organic |
| 388.307 | 160 | C14:0-OH acylcarnitine[1] | NA | NA | NA | 0.39 | major increase | N/S | organic |
| 395.560 | 139 | NA | NA | NA | NA | NA | N/S | N/S | organic |
| 398.328 | 179 | 9-Hexadecenoylcarnitine (Hexadecenoyl-L-carnitine)[1] | NA | NA | NA | 1.05 | major increase | N/S | organic |
| 400.343 | 187 | Spectral Match to palmitoylcarnitine | 0.91 | 12 | 0 | 6.74 | major increase | N/S | organic |
| 400.343 | 209 | Spectral Match to palmitoylcarnitine | 0.91 | 12 | 0 | 6.74 | major increase | N/S | organic |
| 414.322 | 164 | C16:1-OH acylcarnitine[1] | NA | NA | NA | -1.20 | major increase | N/S | organic |
| 414.359 | 193 | C17:0 acylcarnitine[1] | NA | NA | NA | 0.28 | major increase | N/S | organic |
| 416.338 | 171 | C16:0-OH acylcarnitine[1] | NA | NA | NA | -0.35 | major increase | N/S | organic |

| | | | | | | | | | |
|---|---|---|---|---|---|---|---|---|---|
| 424.343 | 184 | C18:2 acylcarnitine[1] | NA | NA | NA | -0.55 | major increase | N/S | organic |
| 426.358 | 229 | Spectral Match to Oleoyl-L-carnitine | 0.89 | 12 | 0 | 9 | major increase | N/S | organic |
| 426.359 | 193 | Spectral Match to Oleoyl-L-carnitine | 0.89 | 12 | 0 | 9 | major increase | N/S | organic |
| 428.373 | 240 | C18:0 acylcarnitine[1] | NA | NA | NA | -3.58 | major increase | N/S | organic |
| 442.354 | 175 | C18:1-OH acylcarnitine[1] | NA | NA | NA | 0.46 | major increase | N/S | organic |
| 444.369 | 185 | C18:0-OH acylcarnitine[1] | NA | NA | NA | -1.01 | major increase | N/S | organic |
| 450.359 | 190 | C20:4 acylcarnitine[1] | NA | NA | NA | 0.26 | major increase | N/S | organic |
| 452.374 | 199 | C20:2 acylcarnitine[1] | NA | NA | NA | -1.18 | major increase | N/S | organic |
| 454.390 | 210 | C20:1 acylcarnitine[1] | NA | NA | NA | -0.40 | major increase | N/S | organic |
| 455.394 | 209 | NA | NA | NA | NA | NA | major increase | N/S | organic |
| 456.406 | 229 | C20:0 acylcarnitine[1] | NA | NA | NA | 0.37 | major increase | N/S | organic |
| 468.369 | 179 | C20:2-OH acylcarnitine[1] | NA | NA | NA | -0.95 | major increase | N/S | organic |
| 472.401 | 202 | C20:1-OH acylcarnitine[1] | NA | NA | NA | 0.54 | major increase | N/S | organic |
| 482.420 | 231 | C22:1 acylcarnitine[1] | NA | NA | NA | -3.07 | major increase | N/S | organic |
| 504.515 | 404 | NA | NA | NA | NA | NA | N/S | N/S | organic |
| 510.453 | 249 | C24:1 acylcarnitine[1] | NA | NA | NA | 0.43 | major increase | N/S | organic |
| 512.468 | 276 | C24:0 acylcarnitine[1] | NA | NA | NA | -0.84 | major increase | N/S | organic |
| 531.388 | 213 | NA | NA | NA | NA | NA | N/S | minor increase | organic |
| 541.368 | 228 | NA | NA | NA | NA | NA | major increase | N/S | organic |
| 549.375 | 187 | NA | NA | NA | NA | NA | N/S | N/S | organic |
| 590.323 | 174 | NA | NA | NA | NA | NA | N/S | N/S | organic |
| 605.424 | 198 | NA | NA | NA | NA | NA | major decrease | N/S | organic |
| 630.403 | 203 | NA | NA | NA | NA | NA | minor decrease | N/S | organic |
| 693.480 | 216 | NA | NA | NA | NA | NA | minor increase | N/S | organic |
| 702.543 | 354 | NA | NA | NA | NA | NA | N/S | N/S | organic |
| 706.502 | 248 | NA | NA | NA | NA | NA | major increase | N/S | organic |
| 720.556 | 354 | NA | NA | NA | NA | NA | minor decrease | minor increase | organic |
| 728.559 | 412 | NA | NA | NA | NA | NA | N/S | N/S | organic |
| 730.562 | 323 | NA | NA | NA | NA | NA | N/S | N/S | organic |
| 731.492 | 428 | NA | NA | NA | NA | NA | N/S | N/S | organic |
| 742.540 | 360 | NA | NA | NA | NA | NA | major increase | N/S | organic |

| | | | | | | | | | |
|---|---|---|---|---|---|---|---|---|---|
| 754.572 | 399 | NA | NA | NA | NA | NA | minor decrease | N/S | organic |
| 760.580 | 340 | PC (34:1)[1] | NA | NA | NA | -8.12 | N/S | N/S | organic |
| 788.609 | 398 | NA | NA | NA | NA | NA | minor increase | N/S | organic |
| 800.617 | 452 | NA | NA | NA | NA | NA | major increase | N/S | organic |
| 800.653 | 357 | NA | NA | NA | NA | NA | N/S | N/S | organic |
| 814.559 | 280 | NA | NA | NA | NA | NA | major increase | N/S | organic |
| 817.577 | 275 | NA | NA | NA | NA | NA | minor increase | N/S | organic |
| 819.594 | 297 | PC family member[1] | NA | NA | NA | NA | minor increase | N/S | organic |
| 828.685 | 320 | NA | NA | NA | NA | NA | major decrease | N/S | organic |
| 866.591 | 301 | NA | NA | NA | NA | NA | minor increase | N/S | organic |
| 1043.701 | 181 | NA | NA | NA | NA | NA | major increase | major decrease | organic |
| 1045.626 | 174 | NA | NA | NA | NA | NA | major increase | N/S | organic |

[1]Annotated based on molecular networking to an annotated sub-network; NA: not applicable; N/S: not statistically significant

**S4 Table. Annotated metabolites of combined extracts perturbed by infection at position D, identified through random forest classifier.**

| Top-ranking metabolites for CL *T.cruzi* strain that met the random forest cut-off criteria | | | | | | | | | |
|---|---|---|---|---|---|---|---|---|---|
| ***m/z*** | **RT (sec)** | **Annotation** | **Cosine Score** | **No. of Shared Peaks** | **Mass Diff. to Library Reference** | **ppm error** | **Impact of Infection at position D (Sylvio X10/4 vs uninfected 147 days)** | **Impact of Infection at position D (CL vs uninfected 90 days)** | **Extract** |
| 160.566 | 189 | NA | NA | NA | NA | NA | N/S | N/S | aqueous |
| 182.082 | 32 | NA | NA | NA | NA | NA | N/S | minor decrease | aqueous |
| 185.596 | 143 | NA | NA | NA | NA | NA | N/S | major decrease | aqueous |
| 186.113 | 149 | NA | NA | NA | NA | NA | N/S | minor increase | aqueous |
| 188.072 | 116 | NA | NA | NA | NA | NA | N/S | minor decrease | aqueous |
| 199.146 | 69 | NA | NA | NA | NA | NA | N/S | minor increase | aqueous |
| 241.155 | 26 | NA | NA | NA | NA | NA | N/S | minor decrease | aqueous |
| 279.138 | 142 | NA | NA | NA | NA | NA | N/S | minor increase | aqueous |
| 296.259 | 187 | NA | NA | NA | NA | NA | N/S | major decrease | aqueous |
| 299.084 | 135 | NA | NA | NA | NA | NA | minor increase | N/S | aqueous |
| 302.126 | 32 | NA | NA | NA | NA | NA | N/S | N/S | aqueous |
| 304.652 | 208 | NA | NA | NA | NA | NA | N/S | minor increase | aqueous |
| 312.254 | 194 | NA | NA | NA | NA | NA | N/S | major decrease | aqueous |
| 316.285 | 191 | NA | NA | NA | NA | NA | N/S | major decrease | aqueous |
| 332.280 | 175 | NA | NA | NA | NA | NA | N/S | major decrease | aqueous |
| 347.220 | 227 | NA | NA | NA | NA | NA | N/S | minor increase | aqueous |
| 358.260 | 164 | NA | NA | NA | NA | NA | major increase | N/S | aqueous |
| 370.178 | 142 | NA | NA | NA | NA | NA | N/S | minor decrease | aqueous |
| 459.281 | 147 | NA | NA | NA | NA | NA | N/S | minor decrease | aqueous |
| 524.372 | 222 | LPC(18:0)[1] | NA | NA | NA | -0.31 | N/S | minor increase | aqueous |
| 546.354 | 219 | NA | NA | NA | NA | NA | N/S | minor increase | aqueous |
| 949.626 | 209 | NA | NA | NA | NA | NA | N/S | major increase | aqueous |
| 1005.689 | 222 | NA | NA | NA | NA | NA | N/S | major increase | aqueous |
| 1049.338 | 20 | NA | NA | NA | NA | NA | N/S | major decrease | aqueous |
| 132.078 | 17 | NA | NA | NA | NA | NA | N/S | N/S | organic |

| | | | | | | | | | |
|---|---|---|---|---|---|---|---|---|---|
| 137.047 | 30 | NA | NA | NA | NA | NA | N/S | major increase | organic |
| 195.094 | 139 | NA | NA | NA | NA | NA | N/S | minor increase | organic |
| 316.250 | 151 | NA | NA | NA | NA | NA | N/S | N/S | organic |
| 509.405 | 212 | NA | NA | NA | NA | NA | N/S | minor increase | organic |
| 522.598 | 509 | NA | NA | NA | NA | NA | N/S | major increase | organic |
| 534.302 | 160 | NA | NA | NA | NA | NA | N/S | N/S | organic |
| 584.430 | 200 | NA | NA | NA | NA | NA | N/S | minor increase | organic |
| 646.614 | 417 | NA | NA | NA | NA | NA | N/S | minor increase | organic |
| 678.430 | 202 | NA | NA | NA | NA | NA | N/S | minor decrease | organic |
| 689.560 | 277 | NA | NA | NA | NA | NA | N/S | major decrease | organic |
| 704.577 | 313 | NA | NA | NA | NA | NA | N/S | minor decrease | organic |
| 774.895 | 136 | NA | NA | NA | NA | NA | N/S | major increase | organic |
| 802.557 | 291 | NA | NA | NA | NA | NA | N/S | minor decrease | organic |
| 904.549 | 227 | NA | NA | NA | NA | NA | N/S | minor decrease | organic |
| **Top-ranking metabolites for CL and Sylvio X10/4 *T.cruzi* strains that met the random forest cut-off criteria** | | | | | | | | | |
| ***m/z*** | **RT (sec)** | **Annotation** | **Cosine Score** | **No. of Shared Peaks** | **Mass Diff. to Library Reference** | **ppm error** | **Impact of Infection at position D (Sylvio X10/4 vs uninfected 147 days)** | **Impact of Infection at position D (CL vs uninfected 90 days)** | **Extract** |
| 193.610 | 145 | NA | NA | NA | NA | NA | major increase | major decrease | aqueous |
| 209.093 | 53 | NA | NA | NA | NA | NA | major decrease | major increase | aqueous |
| 368.201 | 147 | NA | NA | NA | NA | NA | major increase | major decrease | aqueous |
| **Top-ranking metabolites for Sylvio X10/4 *T.cruzi* strain that met the random forest cut-off criteria** | | | | | | | | | |
| ***m/z*** | **RT (sec)** | **Annotation** | **Cosine Score** | **No. of Shared Peaks** | **Mass Diff. to Library Reference** | **ppm error** | **Impact of Infection at position D (Sylvio X10/4 vs uninfected 147 days)** | **Impact of Infection at position D (CL vs uninfected 90 days)** | **Extract** |
| 147.077 | 24 | NA | NA | NA | NA | NA | minor decrease | N/S | aqueous |
| 220.098 | 151 | NA | NA | NA | NA | NA | minor decrease | N/S | aqueous |
| 232.155 | 115 | C4:0 acylcarnitine (butyrylcarnitine)[1] | NA | NA | NA | -1.86 | major increase | N/S | aqueous |

| | | | | | | | | | |
|---|---|---|---|---|---|---|---|---|---|
| 246.171 | 142 | C5:0 acylcarnitine (valerylcarnitine)[1] | NA | NA | NA | -0.33 | major increase | N/S | aqueous |
| 248.150 | 30 | C4:0-OH acylcarnitine[1] | NA | NA | NA | -1.39 | major increase | N/S | aqueous |
| 260.187 | 148 | C6:0 acylcarnitine[1] | NA | NA | NA | 1.03 | major increase | N/S | aqueous |
| 262.166 | 43 | CAR (5:0(OH))[1] | NA | NA | NA | 0.02 | minor increase | N/S | aqueous |
| 276.181 | 104 | NA | NA | NA | NA | NA | major increase | N/S | aqueous |
| 279.093 | 171 | NA | NA | NA | NA | NA | N/S | N/S | aqueous |
| 284.296 | 277 | NA | NA | NA | NA | NA | N/S | N/S | aqueous |
| 286.202 | 154 | C8:1 acylcarnitine[1] | NA | NA | NA | -1.33 | major increase | N/S | aqueous |
| 288.218 | 159 | C8:0 acylcarnitine[1] | NA | NA | NA | -0.11 | major increase | N/S | aqueous |
| 293.212 | 205 | NA | NA | NA | NA | NA | N/S | N/S | aqueous |
| 295.058 | 20 | NA | NA | NA | NA | NA | minor increase | N/S | aqueous |
| 298.274 | 192 | NA | NA | NA | NA | NA | N/S | N/S | aqueous |
| 302.216 | 178 | NA | NA | NA | NA | NA | major increase | N/S | aqueous |
| 304.213 | 149 | CAR (8:1(OH))[1] | NA | NA | NA | 0.17 | major increase | N/S | aqueous |
| 309.207 | 176 | NA | NA | NA | NA | NA | N/S | N/S | aqueous |
| 316.249 | 170 | C10:0 acylcarnitine[1] | NA | NA | NA | -1.05 | major increase | N/S | aqueous |
| 320.256 | 217 | NA | NA | NA | NA | NA | N/S | N/S | aqueous |
| 332.244 | 158 | C10:0-OH acylcarnitine[1] | NA | NA | NA | -0.74 | major increase | N/S | aqueous |
| 353.232 | 179 | NA | NA | NA | NA | NA | N/S | N/S | aqueous |
| 361.278 | 174 | NA | NA | NA | NA | NA | minor increase | N/S | aqueous |
| 368.390 | 294 | NA | NA | NA | NA | NA | minor increase | N/S | aqueous |
| 381.227 | 178 | NA | NA | NA | NA | NA | minor decrease | N/S | aqueous |
| 386.291 | 175 | NA | NA | NA | NA | NA | major increase | N/S | aqueous |
| 388.307 | 181 | C14:0-OH acylcarnitine[1] | NA | NA | NA | 0.39 | major increase | N/S | aqueous |
| 390.206 | 149 | NA | NA | NA | NA | NA | major increase | N/S | aqueous |
| 398.327 | 198 | 9-Hexadecenoylcarnitine (Hexadecenoyl-L-carnitine)[1] | NA | NA | NA | -1.46 | major increase | N/S | aqueous |
| 401.342 | 263 | NA | NA | NA | NA | NA | major decrease | N/S | aqueous |
| 405.226 | 186 | NA | NA | NA | NA | NA | N/S | N/S | aqueous |
| 414.322 | 186 | C16:1-OH acylcarnitine[1] | NA | NA | NA | -1.20 | major increase | N/S | aqueous |
| 416.153 | 33 | NA | NA | NA | NA | NA | major decrease | N/S | aqueous |
| 416.338 | 192 | C16:0-OH acylcarnitine[1] | NA | NA | NA | -0.35 | major increase | major increase | aqueous |

| | | | | | | | | | |
|---|---|---|---|---|---|---|---|---|---|
| 419.373 | 255 | NA | NA | NA | NA | NA | major increase | N/S | aqueous |
| 424.343 | 203 | C18:2 acylcarnitine[1] | NA | NA | NA | -0.55 | major increase | N/S | aqueous |
| 440.338 | 190 | C18:2-OH acylcarnitine[1] | NA | NA | NA | -0.33 | major increase | N/S | aqueous |
| 442.353 | 197 | C18:1-OH acylcarnitine[1] | NA | NA | NA | -1.80 | major increase | N/S | aqueous |
| 444.369 | 204 | C18:0-OH acylcarnitine[1] | NA | NA | NA | -1.01 | major increase | N/S | aqueous |
| 478.190 | 152 | NA | NA | NA | NA | NA | N/S | N/S | aqueous |
| 482.361 | 210 | PC family member[1] | NA | NA | NA | NA | minor decrease | N/S | aqueous |
| 496.340 | 214 | NA | NA | NA | NA | NA | minor decrease | minor increase | aqueous |
| 496.340 | 222 | NA | NA | NA | NA | NA | minor decrease | minor increase | aqueous |
| 508.340 | 212 | NA | NA | NA | NA | NA | minor decrease | N/S | aqueous |
| 508.376 | 215 | LPC(O-18:1)[1] | NA | NA | NA | -2.46 | major decrease | N/S | aqueous |
| 510.283 | 168 | NA | NA | NA | NA | NA | minor increase | N/S | aqueous |
| 512.334 | 180 | NA | NA | NA | NA | NA | minor decrease | minor increase | aqueous |
| 518.216 | 150 | NA | NA | NA | NA | NA | N/S | N/S | aqueous |
| 522.598 | 289 | NA | NA | NA | NA | NA | major decrease | N/S | aqueous |
| 546.352 | 205 | NA | NA | NA | NA | NA | N/S | N/S | aqueous |
| 568.325 | 190 | NA | NA | NA | NA | NA | minor increase | N/S | aqueous |
| 570.340 | 155 | Lysophosphatidylcholine(22:4)[1] | NA | NA | NA | -28.95 | minor increase | N/S | aqueous |
| 600.330 | 174 | NA | NA | NA | NA | NA | minor decrease | N/S | aqueous |
| 704.524 | 287 | NA | NA | NA | NA | NA | minor increase | N/S | aqueous |
| 754.540 | 285 | NA | NA | NA | NA | NA | major decrease | N/S | aqueous |
| 137.526 | 22 | NA | NA | NA | NA | NA | N/S | N/S | organic |
| 184.014 | 20 | NA | NA | NA | NA | NA | N/S | N/S | organic |
| 216.197 | 150 | NA | NA | NA | NA | NA | major decrease | N/S | organic |
| 226.157 | 42 | NA | NA | NA | NA | NA | major decrease | N/S | organic |
| 228.196 | 177 | NA | NA | NA | NA | NA | major decrease | N/S | organic |
| 238.024 | 21 | NA | NA | NA | NA | NA | minor increase | N/S | organic |
| 248.151 | 19 | C4:0-OH acylcarnitine[1] | NA | NA | NA | 2.64 | major increase | N/S | organic |
| 268.264 | 216 | NA | NA | NA | NA | NA | minor decrease | N/S | organic |
| 274.183 | 153 | NA | NA | NA | NA | NA | major increase | N/S | organic |
| 279.095 | 159 | NA | NA | NA | NA | NA | major decrease | N/S | organic |

| | | | | | | | | | |
|---|---|---|---|---|---|---|---|---|---|
| 291.611 | 172 | NA | NA | NA | NA | NA | N/S | N/S | organic |
| 302.207 | 157 | NA | NA | NA | NA | NA | major increase | N/S | organic |
| 332.244 | 146 | C10:0-OH acylcarnitine[1] | NA | NA | NA | -0.74 | major increase | N/S | organic |
| 340.323 | 274 | NA | NA | NA | NA | NA | major increase | N/S | organic |
| 360.275 | 153 | C12:0-OH acylcarnitine[1] | NA | NA | NA | -1.52 | major increase | N/S | organic |
| 370.296 | 166 | C14:1 acylcarnitine[1] | NA | NA | NA | -0.76 | major increase | N/S | organic |
| 371.053 | 40 | NA | NA | NA | NA | NA | N/S | N/S | organic |
| 386.291 | 157 | C14:1 acylcarnitine[1] | NA | NA | NA | -0.76 | major increase | N/S | organic |
| 388.307 | 160 | C14:0-OH acylcarnitine[1] | NA | NA | NA | 0.39 | major increase | N/S | organic |
| 395.560 | 139 | NA | NA | NA | NA | NA | major decrease | N/S | organic |
| 398.328 | 179 | 9-Hexadecenoylcarnitine (Hexadecenoyl-L-carnitine)[1] | NA | NA | NA | 1.05 | major increase | N/S | organic |
| 400.343 | 187 | Spectral Match to palmitoylcarnitine | 0.91 | 12 | 0 | 6.74 | major increase | N/S | organic |
| 400.343 | 209 | Spectral Match to palmitoylcarnitine | 0.91 | 12 | 0 | 6.74 | major increase | N/S | organic |
| 414.322 | 164 | C16:1-OH acylcarnitine[1] | NA | NA | NA | -1.20 | major increase | N/S | organic |
| 414.359 | 193 | C17:0 acylcarnitine[1] | NA | NA | NA | 0.28 | major increase | N/S | organic |
| 416.338 | 171 | C16:0-OH acylcarnitine[1] | NA | NA | NA | -0.35 | major increase | N/S | organic |
| 424.343 | 184 | C18:2 acylcarnitine1 | NA | NA | NA | -0.55 | major increase | N/S | organic |
| 426.358 | 229 | Spectral Match to Oleoyl-L-carnitine | 0.89 | 12 | 0 | 9 | major increase | N/S | organic |
| 426.359 | 193 | Spectral Match to Oleoyl-L-carnitine | 0.89 | 12 | 0 | 9 | major increase | N/S | organic |
| 428.373 | 240 | C18:0 acylcarnitine[1] | NA | NA | NA | -3.58 | major increase | N/S | organic |
| 442.354 | 175 | C18:1-OH acylcarnitine[1] | NA | NA | NA | 0.46 | major increase | N/S | organic |
| 444.369 | 185 | C18:0-OH acylcarnitine[1] | NA | NA | NA | -1.01 | major increase | N/S | organic |
| 450.359 | 190 | C20:4 acylcarnitine[1] | NA | NA | NA | 0.26 | major increase | N/S | organic |
| 452.374 | 199 | C20:2 acylcarnitine[1] | NA | NA | NA | -1.18 | major increase | N/S | organic |
| 454.390 | 210 | C20:1 acylcarnitine[1] | NA | NA | NA | -0.40 | major increase | N/S | organic |
| 455.394 | 209 | NA | NA | NA | NA | NA | major increase | N/S | organic |
| 456.406 | 229 | C20:0 acylcarnitine[1] | NA | NA | NA | 0.37 | major increase | N/S | organic |
| 468.369 | 179 | C20:2-OH acylcarnitine[1] | NA | NA | NA | -0.95 | major increase | N/S | organic |
| 472.401 | 202 | C20:1-OH acylcarnitine[1] | NA | NA | NA | 0.54 | major increase | N/S | organic |

| | | | | | | | | | |
|---|---|---|---|---|---|---|---|---|---|
| 482.420 | 231 | C22:1 acylcarnitine[1] | NA | NA | NA | -3.07 | major increase | N/S | organic |
| 504.515 | 404 | NA | NA | NA | NA | NA | N/S | major decrease | organic |
| 510.453 | 249 | C24:1 acylcarnitine[1] | NA | NA | NA | 0.43 | major increase | N/S | organic |
| 512.468 | 276 | C24:0 acylcarnitine[1] | NA | NA | NA | -0.84 | major increase | N/S | organic |
| 531.388 | 213 | NA | NA | NA | NA | NA | N/S | N/S | organic |
| 541.368 | 228 | NA | NA | NA | NA | NA | major increase | N/S | organic |
| 549.375 | 187 | NA | NA | NA | NA | NA | N/S | N/S | organic |
| 590.323 | 174 | NA | NA | NA | NA | NA | N/S | N/S | organic |
| 605.424 | 198 | NA | NA | NA | NA | NA | major decrease | N/S | organic |
| 630.403 | 203 | NA | NA | NA | NA | NA | minor decrease | N/S | organic |
| 693.480 | 216 | NA | NA | NA | NA | NA | minor decrease | N/S | organic |
| 702.543 | 354 | NA | NA | NA | NA | NA | N/S | N/S | organic |
| 706.502 | 248 | NA | NA | NA | NA | NA | major increase | N/S | organic |
| 720.556 | 354 | NA | NA | NA | NA | NA | minor decrease | N/S | organic |
| 728.559 | 412 | NA | NA | NA | NA | NA | N/S | N/S | organic |
| 730.562 | 323 | NA | NA | NA | NA | NA | N/S | N/S | organic |
| 731.492 | 428 | NA | NA | NA | NA | NA | N/S | N/S | organic |
| 742.540 | 360 | NA | NA | NA | NA | NA | major increase | N/S | organic |
| 754.572 | 399 | NA | NA | NA | NA | NA | minor decrease | N/S | organic |
| 760.580 | 340 | PC (34:1)[1] | NA | NA | NA | -8.12 | N/S | N/S | organic |
| 787.535 | 427 | NA | NA | NA | NA | NA | N/S | N/S | organic |
| 788.609 | 398 | NA | NA | NA | NA | NA | minor increase | N/S | organic |
| 800.617 | 452 | NA | NA | NA | NA | NA | major increase | N/S | organic |
| 800.653 | 357 | NA | NA | NA | NA | NA | N/S | N/S | organic |
| 814.559 | 280 | NA | NA | NA | NA | NA | major increase | N/S | organic |
| 817.577 | 275 | NA | NA | NA | NA | NA | minor increase | N/S | organic |
| 819.594 | 297 | PC family member[1] | NA | NA | NA | NA | minor increase | N/S | organic |
| 828.685 | 320 | NA | NA | NA | NA | NA | major decrease | N/S | organic |
| 866.591 | 301 | NA | NA | NA | NA | NA | minor increase | N/S | organic |
| 1043.701 | 181 | NA | NA | NA | NA | NA | major increase | N/S | organic |
| 1045.626 | 174 | NA | NA | NA | NA | NA | major increase | N/S | organic |

[1]Annotated based on molecular networking to an annotated sub-network; NA: not applicable; N/S: not statistically significant

**S5 Table. Annotated metabolites of combined extracts perturbed by infection at positions A-D, identified through random forest classifier.**

| Top-ranking metabolites for CL *T.cruzi* strain that met the random forest cut-off criteria | | | | | | | | | |
|---|---|---|---|---|---|---|---|---|---|
| ***m/z*** | **RT (sec)** | **Annotation** | **Cosine Score** | **No. of Shared Peaks** | **Mass Diff. to Library Reference** | **ppm error** | **Impact of Infection (Sylvio X10/4 vs uninfected 147 days)** | **Impact of Infection (CL vs uninfected 90 days)** | **Extract** |
| 149.024 | 184 | NA | NA | NA | NA | NA | N/S | N/S | aqueous |
| 153.042 | 32 | NA | NA | NA | NA | NA | minor decrease | N/S | aqueous |
| 154.059 | 19 | NA | NA | NA | NA | NA | minor increase | N/S | aqueous |
| 155.071 | 123 | NA | NA | NA | NA | NA | minor decrease | minor increase | aqueous |
| 160.566 | 189 | NA | NA | NA | NA | NA | N/S | N/S | aqueous |
| 166.087 | 101 | NA | NA | NA | NA | NA | N/S | minor decrease | aqueous |
| 172.098 | 143 | NA | NA | NA | NA | NA | N/S | N/S | aqueous |
| 178.133 | 135 | NA | NA | NA | NA | NA | N/S | major decrease | aqueous |
| 182.082 | 32 | NA | NA | NA | NA | NA | minor decrease | minor decrease | aqueous |
| 184.074 | 25 | phosphocholine[1] | NA | NA | NA | -2.27 | minor increase | minor increase | aqueous |
| 185.596 | 143 | NA | NA | NA | NA | NA | major increase | major decrease | aqueous |
| 198.092 | 25 | NA | NA | NA | NA | NA | N/S | N/S | aqueous |
| 200.632 | 80 | NA | NA | NA | NA | NA | N/S | minor increase | aqueous |
| 211.145 | 151 | NA | NA | NA | NA | NA | N/S | major decrease | aqueous |
| 216.197 | 163 | NA | NA | NA | NA | NA | N/S | N/S | aqueous |
| 218.140 | 39 | NA | NA | NA | NA | NA | N/S | minor decrease | aqueous |
| 223.097 | 184 | NA | NA | NA | NA | NA | N/S | minor increase | aqueous |
| 227.092 | 138 | NA | NA | NA | NA | NA | N/S | minor increase | aqueous |
| 229.144 | 170 | NA | NA | NA | NA | NA | N/S | N/S | aqueous |
| 235.673 | 145 | NA | NA | NA | NA | NA | N/S | N/S | aqueous |
| 241.155 | 26 | NA | NA | NA | NA | NA | N/S | minor decrease | aqueous |
| 251.103 | 147 | NA | NA | NA | NA | NA | N/S | minor decrease | aqueous |
| 252.233 | 190 | NA | NA | NA | NA | NA | N/S | major decrease | aqueous |
| 268.108 | 33 | NA | NA | NA | NA | NA | major increase | minor decrease | aqueous |
| 270.244 | 200 | NA | NA | NA | NA | NA | N/S | major decrease | aqueous |

| | | | | | | | | | |
|---|---|---|---|---|---|---|---|---|---|
| 278.248 | 179 | NA | NA | NA | NA | NA | N/S | major decrease | aqueous |
| 285.134 | 146 | NA | NA | NA | NA | NA | N/S | minor increase | aqueous |
| 285.290 | 177 | NA | NA | NA | NA | NA | N/S | N/S | aqueous |
| 292.227 | 199 | NA | NA | NA | NA | NA | N/S | major decrease | aqueous |
| 296.259 | 187 | NA | NA | NA | NA | NA | N/S | major decrease | aqueous |
| 309.207 | 176 | NA | NA | NA | NA | NA | N/S | minor increase | aqueous |
| 312.254 | 194 | NA | NA | NA | NA | NA | N/S | major decrease | aqueous |
| 316.285 | 191 | NA | NA | NA | NA | NA | major increase | major decrease | aqueous |
| 318.241 | 212 | NA | NA | NA | NA | NA | N/S | major decrease | aqueous |
| 320.256 | 217 | NA | NA | NA | NA | NA | major increase | major decrease | aqueous |
| 325.238 | 227 | NA | NA | NA | NA | NA | N/S | major increase | aqueous |
| 332.180 | 144 | NA | NA | NA | NA | NA | N/S | N/S | aqueous |
| 332.280 | 175 | NA | NA | NA | NA | NA | N/S | major decrease | aqueous |
| 336.251 | 190 | NA | NA | NA | NA | NA | N/S | major decrease | aqueous |
| 343.230 | 204 | NA | NA | NA | NA | NA | N/S | minor increase | aqueous |
| 344.228 | 143 | NA | NA | NA | NA | NA | N/S | minor decrease | aqueous |
| 353.268 | 202 | NA | NA | NA | NA | NA | N/S | minor increase | aqueous |
| 354.301 | 239 | NA | NA | NA | NA | NA | N/S | N/S | aqueous |
| 367.247 | 186 | NA | NA | NA | NA | NA | N/S | minor increase | aqueous |
| 368.280 | 177 | NA | NA | NA | NA | NA | N/S | major increase | aqueous |
| 370.178 | 142 | NA | NA | NA | NA | NA | N/S | minor decrease | aqueous |
| 370.296 | 188 | C14:1 acylcarnitine[1] | NA | NA | NA | -0.76 | N/S | major increase | aqueous |
| 371.039 | 31 | NA | NA | NA | NA | NA | N/S | N/S | aqueous |
| 375.216 | 186 | NA | NA | NA | NA | NA | minor decrease | minor increase | aqueous |
| 379.210 | 214 | NA | NA | NA | NA | NA | N/S | minor increase | aqueous |
| 380.113 | 41 | NA | NA | NA | NA | NA | N/S | major decrease | aqueous |
| 382.296 | 182 | C15:2 acylcarnitine[1] | NA | NA | NA | -0.74 | major increase | minor increase | aqueous |
| 383.163 | 74 | NA | NA | NA | NA | NA | N/S | minor decrease | aqueous |
| 384.179 | 139 | NA | NA | NA | NA | NA | minor increase | minor decrease | aqueous |
| 388.255 | 144 | NA | NA | NA | NA | NA | N/S | minor decrease | aqueous |
| 400.269 | 187 | C14:2-DC acylcarnitine[1] | NA | NA | NA | -3.65 | minor decrease | minor increase | aqueous |
| 407.242 | 190 | NA | NA | NA | NA | NA | minor decrease | minor increase | aqueous |

| | | | | | | | | | |
|---|---|---|---|---|---|---|---|---|---|
| 412.190 | 137 | NA | NA | NA | NA | NA | minor increase | minor decrease | aqueous |
| 428.374 | 217 | C18:0 acylcarnitine[1] | NA | NA | NA | -1.24 | major increase | major increase | aqueous |
| 432.281 | 145 | NA | NA | NA | NA | NA | N/S | minor decrease | aqueous |
| 438.299 | 210 | NA | NA | NA | NA | NA | N/S | minor increase | aqueous |
| 446.196 | 147 | NA | NA | NA | NA | NA | minor increase | minor decrease | aqueous |
| 462.191 | 140 | NA | NA | NA | NA | NA | minor increase | minor decrease | aqueous |
| 462.191 | 148 | NA | NA | NA | NA | NA | minor increase | minor decrease | aqueous |
| 465.210 | 148 | NA | NA | NA | NA | NA | minor increase | minor decrease | aqueous |
| 466.330 | 224 | NA | NA | NA | NA | NA | N/S | minor increase | aqueous |
| 468.309 | 193 | LPC (14:0)[1] | NA | NA | NA | -1.20 | minor decrease | minor increase | aqueous |
| 470.215 | 141 | NA | NA | NA | NA | NA | minor increase | minor decrease | aqueous |
| 476.307 | 146 | NA | NA | NA | NA | NA | N/S | minor decrease | aqueous |
| 480.165 | 138 | NA | NA | NA | NA | NA | N/S | minor decrease | aqueous |
| 482.325 | 215 | NA | NA | NA | NA | NA | N/S | N/S | aqueous |
| 482.325 | 234 | NA | NA | NA | NA | NA | N/S | minor increase | aqueous |
| 492.239 | 166 | NA | NA | NA | NA | NA | N/S | N/S | aqueous |
| 494.288 | 186 | NA | NA | NA | NA | NA | N/S | minor increase | aqueous |
| 496.340 | 214 | NA | NA | NA | NA | NA | N/S | minor increase | aqueous |
| 520.333 | 147 | NA | NA | NA | NA | NA | N/S | minor decrease | aqueous |
| 520.340 | 216 | NA | NA | NA | NA | NA | minor increase | minor increase | aqueous |
| 520.510 | 313 | NA | NA | NA | NA | NA | N/S | minor increase | aqueous |
| 524.372 | 222 | LPC (18:0)[1] | NA | NA | NA | -0.31 | N/S | minor increase | aqueous |
| 547.335 | 147 | NA | NA | NA | NA | NA | N/S | N/S | aqueous |
| 560.334 | 180 | NA | NA | NA | NA | NA | N/S | minor increase | aqueous |
| 605.357 | 169 | NA | NA | NA | NA | NA | N/S | major decrease | aqueous |
| 616.177 | 186 | NA | NA | NA | NA | NA | N/S | minor decrease | aqueous |
| 634.187 | 180 | NA | NA | NA | NA | NA | N/S | minor decrease | aqueous |
| 657.204 | 180 | NA | NA | NA | NA | NA | N/S | minor decrease | aqueous |
| 700.528 | 309 | NA | NA | NA | NA | NA | N/S | N/S | aqueous |
| 703.575 | 286 | NA | NA | NA | NA | NA | N/S | major increase | aqueous |
| 706.538 | 312 | PC (30:0)[1] | NA | NA | NA | -1.74 | N/S | major increase | aqueous |
| 724.526 | 307 | NA | NA | NA | NA | NA | N/S | N/S | aqueous |

| | | | | | | | | | |
|---|---|---|---|---|---|---|---|---|---|
| 758.503 | 160 | NA | NA | NA | NA | NA | N/S | minor increase | aqueous |
| 758.570 | 309 | NA | NA | NA | NA | NA | N/S | minor increase | aqueous |
| 790.559 | 255 | NA | NA | NA | NA | NA | N/S | N/S | aqueous |
| 808.584 | 309 | NA | NA | NA | NA | NA | N/S | N/S | aqueous |
| 931.451 | 189 | NA | NA | NA | NA | NA | major increase | major decrease | aqueous |
| 949.626 | 209 | NA | NA | NA | NA | NA | N/S | major increase | aqueous |
| 1047.736 | 222 | NA | NA | NA | NA | NA | N/S | major increase | aqueous |
| 1249.357 | 179 | NA | NA | NA | NA | NA | N/S | minor decrease | aqueous |
| 209.093 | 57 | NA | NA | NA | NA | NA | major decrease | major increase | organic |
| 228.196 | 177 | NA | NA | NA | NA | NA | N/S | minor decrease | organic |
| 229.101 | 116 | NA | NA | NA | NA | NA | N/S | minor increase | organic |
| 269.090 | 26 | NA | NA | NA | NA | NA | major decrease | major increase | organic |
| 296.259 | 162 | NA | NA | NA | NA | NA | N/S | major decrease | organic |
| 296.259 | 181 | NA | NA | NA | NA | NA | N/S | major decrease | organic |
| 310.312 | 256 | NA | NA | NA | NA | NA | N/S | N/S | organic |
| 339.290 | 178 | NA | NA | NA | NA | NA | N/S | major decrease | organic |
| 399.310 | 216 | NA | NA | NA | NA | NA | N/S | N/S | organic |
| 401.347 | 189 | NA | NA | NA | NA | NA | minor increase | major increase | organic |
| 480.515 | 416 | NA | NA | NA | NA | NA | N/S | major decrease | organic |
| 504.515 | 404 | NA | NA | NA | NA | NA | major increase | major decrease | organic |
| 506.531 | 422 | NA | NA | NA | NA | NA | major increase | major decrease | organic |
| 508.340 | 214 | NA | NA | NA | NA | NA | N/S | minor decrease | organic |
| 508.546 | 444 | NA | NA | NA | NA | NA | N/S | major decrease | organic |
| 517.369 | 222 | NA | NA | NA | NA | NA | minor increase | major increase | organic |
| 556.303 | 187 | NA | NA | NA | NA | NA | N/S | N/S | organic |
| 574.278 | 166 | NA | NA | NA | NA | NA | N/S | N/S | organic |
| 604.362 | 208 | NA | NA | NA | NA | NA | N/S | N/S | organic |
| 620.436 | 184 | NA | NA | NA | NA | NA | N/S | N/S | organic |
| 726.542 | 521 | NA | NA | NA | NA | NA | N/S | N/S | organic |
| 772.618 | 417 | NA | NA | NA | NA | NA | N/S | minor decrease | organic |
| 782.570 | 418 | NA | NA | NA | NA | NA | N/S | minor increase | organic |
| 804.522 | 284 | NA | NA | NA | NA | NA | N/S | N/S | organic |

| | | | | | | | | | |
|---|---|---|---|---|---|---|---|---|---|
| 834.594 | 329 | NA | NA | NA | NA | NA | N/S | minor increase | organic |
| 850.632 | 417 | NA | NA | NA | NA | NA | N/S | minor increase | organic |
| 932.583 | 248 | NA | NA | NA | NA | NA | N/S | minor increase | organic |
| 1408.040 | 303 | NA | NA | NA | NA | NA | N/S | N/S | organic |
| **Top-ranking metabolites for CL and Sylvio X10/4 *T.cruzi* strains that met the random forest cut-off criteria** | | | | | | | | | |
| ***m/z*** | **RT (sec)** | **Annotation** | **Cosine Score** | **No. of Shared Peaks** | **Mass Diff. to Library Reference** | **ppm error** | **Impact of Infection (Sylvio X10/4 vs uninfected 147 days)** | **Impact of Infection (CL vs uninfected 90 days)** | **Extract** |
| 147.077 | 24 | NA | NA | NA | NA | NA | minor decrease | N/S | aqueous |
| 150.059 | 27 | NA | NA | NA | NA | NA | minor decrease | minor decrease | aqueous |
| 193.610 | 145 | NA | NA | NA | NA | NA | major increase | major decrease | aqueous |
| 209.093 | 53 | NA | NA | NA | NA | NA | major decrease | major increase | aqueous |
| 280.264 | 206 | NA | NA | NA | NA | NA | major increase | major decrease | aqueous |
| 320.256 | 206 | NA | NA | NA | NA | NA | major increase | major decrease | aqueous |
| 368.201 | 147 | NA | NA | NA | NA | NA | major increase | minor decrease | aqueous |
| 381.227 | 178 | NA | NA | NA | NA | NA | minor decrease | minor increase | aqueous |
| 412.200 | 132 | NA | NA | NA | NA | NA | minor increase | minor decrease | aqueous |
| 426.359 | 210 | Spectral Match to Oleoyl-L-carnitine | 0.85 | 10 | 0 | 0 | major increase | major increase | aqueous |
| 440.338 | 190 | C18:2-OH acylcarnitine[1] | NA | NA | NA | -0.33 | major increase | minor increase | aqueous |
| 464.206 | 147 | NA | NA | NA | NA | NA | minor increase | minor decrease | aqueous |
| 153.041 | 28 | NA | NA | NA | NA | NA | minor decrease | minor increase | organic |
| 269.093 | 43 | NA | NA | NA | NA | NA | major decrease | major increase | organic |
| 326.379 | 231 | NA | NA | NA | NA | NA | major increase | major decrease | organic |
| 414.359 | 193 | C17:0 acylcarnitine[1] | NA | NA | NA | 0.28 | major increase | minor increase | organic |
| 456.406 | 229 | C20:0 acylcarnitine[1] | NA | NA | NA | 0.37 | major increase | major increase | organic |
| 646.614 | 417 | NA | NA | NA | NA | NA | minor decrease | minor increase | organic |
| 801.578 | 282 | NA | NA | NA | NA | NA | minor increase | minor decrease | organic |
| **Top-ranking metabolites for Sylvio X10/4 *T.cruzi* strain that met the random forest cut-off criteria** | | | | | | | | | |
| ***m/z*** | **RT (sec)** | **Annotation** | **Cosine Score** | **No. of Shared Peaks** | **Mass Diff. to** | **ppm error** | **Impact of Infection (Sylvio X10/4 vs** | **Impact of Infection (CL** | **Extract** |

| | | | | | Library Reference | | uninfected 147 days) | vs uninfected 90 days) | |
|---|---|---|---|---|---|---|---|---|---|
| 147.114 | 14 | NA | NA | NA | NA | NA | minor decrease | N/S | aqueous |
| 148.061 | 27 | NA | NA | NA | NA | NA | minor increase | N/S | aqueous |
| 149.06 | 55 | NA | NA | NA | NA | NA | N/S | N/S | aqueous |
| 150.058 | 45 | NA | NA | NA | NA | NA | N/S | N/S | aqueous |
| 150.058 | 47 | NA | NA | NA | NA | NA | minor decrease | N/S | aqueous |
| 172.985 | 26 | NA | NA | NA | NA | NA | minor increase | N/S | aqueous |
| 175.120 | 19 | NA | NA | NA | NA | NA | minor decrease | N/S | aqueous |
| 176.072 | 143 | NA | NA | NA | NA | NA | N/S | N/S | aqueous |
| 188.069 | 56 | NA | NA | NA | NA | NA | N/S | N/S | aqueous |
| 195.138 | 202 | NA | NA | NA | NA | NA | N/S | N/S | aqueous |
| 201.088 | 33 | NA | NA | NA | NA | NA | minor decrease | N/S | aqueous |
| 202.072 | 43 | NA | NA | NA | NA | NA | minor increase | N/S | aqueous |
| 220.136 | 160 | NA | NA | NA | NA | NA | major increase | N/S | aqueous |
| 227.177 | 137 | NA | NA | NA | NA | NA | N/S | N/S | aqueous |
| 228.135 | 138 | NA | NA | NA | NA | NA | N/S | N/S | aqueous |
| 228.233 | 244 | NA | NA | NA | NA | NA | N/S | N/S | aqueous |
| 228.234 | 225 | NA | NA | NA | NA | NA | major decrease | N/S | aqueous |
| 232.155 | 115 | C4:0 acylcarnitine (butyrylcarnitine)[1] | NA | NA | NA | -1.86 | minor increase | N/S | aqueous |
| 235.199 | 140 | NA | NA | NA | NA | NA | N/S | minor increase | aqueous |
| 238.155 | 148 | NA | NA | NA | NA | NA | N/S | N/S | aqueous |
| 248.150 | 30 | C4:0-OH acylcarnitine[1] | NA | NA | NA | -1.39 | major increase | N/S | aqueous |
| 251.249 | 189 | NA | NA | NA | NA | NA | N/S | N/S | aqueous |
| 254.149 | 148 | NA | NA | NA | NA | NA | N/S | N/S | aqueous |
| 255.233 | 210 | NA | NA | NA | NA | NA | major increase | N/S | aqueous |
| 257.117 | 31 | NA | NA | NA | NA | NA | minor increase | N/S | aqueous |
| 258.106 | 140 | NA | NA | NA | NA | NA | N/S | N/S | aqueous |
| 258.171 | 145 | C6:1 acylcarnitine[1] | NA | NA | NA | -0.32 | major increase | N/S | aqueous |
| 260.187 | 148 | C6:0 acylcarnitine[1] | NA | NA | NA | 1.03 | major increase | N/S | aqueous |
| 262.166 | 43 | CAR (5:0(OH))[1] | NA | NA | NA | 0.02 | minor increase | minor decrease | aqueous |
| 267.098 | 75 | NA | NA | NA | NA | NA | N/S | N/S | aqueous |

| | | | | | | | | | |
|---|---|---|---|---|---|---|---|---|---|
| 269.175 | 165 | NA | NA | NA | NA | NA | N/S | N/S | aqueous |
| 274.166 | 141 | NA | NA | NA | NA | NA | minor increase | N/S | aqueous |
| 274.184 | 166 | NA | NA | NA | NA | NA | major increase | N/S | aqueous |
| 275.202 | 206 | NA | NA | NA | NA | NA | minor decrease | N/S | aqueous |
| 276.181 | 104 | NA | NA | NA | NA | NA | major increase | N/S | aqueous |
| 276.181 | 139 | NA | NA | NA | NA | NA | major increase | N/S | aqueous |
| 280.093 | 16 | NA | NA | NA | NA | NA | N/S | N/S | aqueous |
| 286.202 | 154 | C8:1 acylcarnitine[1] | NA | NA | NA | -1.33 | major increase | N/S | aqueous |
| 288.218 | 159 | C8:0 acylcarnitine[1] | NA | NA | NA | -0.11 | major increase | N/S | aqueous |
| 289.142 | 218 | NA | NA | NA | NA | NA | minor decrease | minor increase | aqueous |
| 302.197 | 145 | NA | NA | NA | NA | NA | major increase | N/S | aqueous |
| 302.216 | 178 | NA | NA | NA | NA | NA | major increase | N/S | aqueous |
| 304.213 | 149 | CAR (8:1(OH))[1] | NA | NA | NA | 0.17 | major increase | N/S | aqueous |
| 316.249 | 170 | C10:0 acylcarnitine[1] | NA | NA | NA | -1.05 | major increase | N/S | aqueous |
| 332.244 | 158 | C10:0-OH acylcarnitine[1] | NA | NA | NA | -0.74 | major increase | N/S | aqueous |
| 360.275 | 170 | C12:0-OH acylcarnitine[1] | NA | NA | NA | -1.52 | major increase | N/S | aqueous |
| 369.264 | 185 | NA | NA | NA | NA | NA | N/S | N/S | aqueous |
| 384.116 | 136 | NA | NA | NA | NA | NA | major increase | N/S | aqueous |
| 386.211 | 146 | NA | NA | NA | NA | NA | major increase | minor decrease | aqueous |
| 386.291 | 175 | NA | NA | NA | NA | NA | major increase | N/S | aqueous |
| 388.307 | 181 | C14:0-OH acylcarnitine[1] | NA | NA | NA | 0.39 | major increase | minor increase | aqueous |
| 389.232 | 202 | NA | NA | NA | NA | NA | minor decrease | minor increase | aqueous |
| 390.206 | 149 | NA | NA | NA | NA | NA | minor increase | N/S | aqueous |
| 398.195 | 143 | NA | NA | NA | NA | NA | minor increase | N/S | aqueous |
| 398.327 | 198 | 9-Hexadecenoylcarnitine (Hexadecenoyl-L-carnitine)[1] | NA | NA | NA | -1.46 | major increase | major increase | aqueous |
| 400.343 | 205 | Spectral Match to Palmitoylcarnitine | 0.91 | 11 | 0 | 2 | major increase | major increase | aqueous |
| 412.210 | 147 | PC family member[1] | NA | NA | NA | NA | minor increase | N/S | aqueous |
| 412.307 | 179 | C16:2-OH acylcarnitine[1] | NA | NA | NA | 0.37 | major increase | N/S | aqueous |
| 414.322 | 186 | C16:1-OH acylcarnitine[1] | NA | NA | NA | -1.20 | major increase | minor increase | aqueous |
| 415.158 | 29 | NA | NA | NA | NA | NA | minor decrease | N/S | aqueous |
| 416.153 | 33 | NA | NA | NA | NA | NA | major decrease | N/S | aqueous |

| | | | | | | | | | |
|---|---|---|---|---|---|---|---|---|---|
| 416.338 | 192 | C16:0-OH acylcarnitine[1] | NA | NA | NA | -0.35 | major increase | minor increase | aqueous |
| 421.221 | 177 | NA | NA | NA | NA | NA | minor decrease | minor increase | aqueous |
| 422.195 | 146 | NA | NA | NA | NA | NA | minor increase | minor decrease | aqueous |
| 428.205 | 146 | NA | NA | NA | NA | NA | minor increase | N/S | aqueous |
| 436.211 | 150 | NA | NA | NA | NA | NA | minor increase | N/S | aqueous |
| 442.353 | 197 | C18:1-OH acylcarnitine[1] | NA | NA | NA | -1.80 | major increase | minor increase | aqueous |
| 444.369 | 204 | C18:0-OH acylcarnitine[1] | NA | NA | NA | -1.01 | major increase | major increase | aqueous |
| 453.370 | 191 | NA | NA | NA | NA | NA | minor increase | N/S | aqueous |
| 468.368 | 198 | C20:2-OH acylcarnitine[1] | NA | NA | NA | -3.09 | major increase | N/S | aqueous |
| 480.201 | 147 | NA | NA | NA | NA | NA | minor increase | N/S | aqueous |
| 480.343 | 212 | LPC (O-16:1)[1] | NA | NA | NA | -6.14 | major decrease | N/S | aqueous |
| 492.272 | 183 | NA | NA | NA | NA | NA | minor increase | N/S | aqueous |
| 508.303 | 174 | NA | NA | NA | NA | NA | minor decrease | minor increase | aqueous |
| 510.283 | 168 | NA | NA | NA | NA | NA | minor increase | N/S | aqueous |
| 528.293 | 157 | NA | NA | NA | NA | NA | major increase | N/S | aqueous |
| 552.330 | 169 | PC family member[1] | NA | NA | NA | NA | minor increase | N/S | aqueous |
| 568.324 | 162 | NA | NA | NA | NA | NA | minor increase | N/S | aqueous |
| 568.325 | 190 | NA | NA | NA | NA | NA | minor increase | N/S | aqueous |
| 570.340 | 155 | Lysophosphatidylcholine(22:4)[1] | NA | NA | NA | -28.95 | minor increase | N/S | aqueous |
| 570.341 | 173 | NA | NA | NA | NA | NA | minor increase | N/S | aqueous |
| 574.279 | 166 | NA | NA | NA | NA | NA | N/S | N/S | aqueous |
| 584.335 | 182 | NA | NA | NA | NA | NA | minor decrease | minor increase | aqueous |
| 594.339 | 163 | NA | NA | NA | NA | NA | minor increase | N/S | aqueous |
| 614.310 | 174 | NA | NA | NA | NA | NA | minor decrease | N/S | aqueous |
| 632.319 | 153 | NA | NA | NA | NA | NA | minor decrease | N/S | aqueous |
| 1011.639 | 197 | NA | NA | NA | NA | NA | N/S | N/S | aqueous |
| 1135.509 | 200 | NA | NA | NA | NA | NA | major increase | N/S | aqueous |
| 137.047 | 30 | NA | NA | NA | NA | NA | minor decrease | minor increase | organic |
| 138.050 | 29 | NA | NA | NA | NA | NA | minor decrease | minor increase | organic |
| 147.078 | 19 | NA | NA | NA | NA | NA | major decrease | N/S | organic |
| 153.043 | 34 | NA | NA | NA | NA | NA | major decrease | N/S | organic |
| 159.027 | 20 | NA | NA | NA | NA | NA | N/S | minor decrease | organic |

| | | | | | | | | | |
|---|---|---|---|---|---|---|---|---|---|
| 200.041 | 22 | NA | NA | NA | NA | NA | N/S | N/S | organic |
| 242.250 | 213 | NA | NA | NA | NA | NA | N/S | N/S | organic |
| 248.150 | 36 | C4:0-OH acylcarnitine[1] | NA | NA | NA | -1.39 | major increase | N/S | organic |
| 248.151 | 19 | C4:0-OH acylcarnitine[1] | NA | NA | NA | 2.64 | major increase | N/S | organic |
| 250.179 | 157 | NA | NA | NA | NA | NA | major decrease | N/S | organic |
| 260.187 | 141 | C6:0 acylcarnitine[1] | NA | NA | NA | 1.03 | minor increase | N/S | organic |
| 274.183 | 153 | NA | NA | NA | NA | NA | major increase | N/S | organic |
| 302.207 | 157 | NA | NA | NA | NA | NA | major increase | N/S | organic |
| 302.215 | 161 | NA | NA | NA | NA | NA | major increase | N/S | organic |
| 304.213 | 141 | CAR(8:1(OH))[1] | NA | NA | NA | 0.17 | major increase | N/S | organic |
| 326.379 | 206 | NA | NA | NA | NA | NA | major increase | major decrease | organic |
| 332.244 | 146 | C10:0-OH acylcarnitine[1] | NA | NA | NA | -0.74 | major increase | N/S | organic |
| 343.333 | 189 | NA | NA | NA | NA | NA | major increase | N/S | organic |
| 360.275 | 153 | C12:0-OH acylcarnitine[1] | NA | NA | NA | -1.52 | major increase | N/S | organic |
| 368.280 | 158 | C14:2 acylcarnitine[1] | NA | NA | NA | -1.72 | major increase | N/S | organic |
| 370.296 | 166 | C14:1 acylcarnitine[1] | NA | NA | NA | -0.76 | major increase | N/S | organic |
| 370.369 | 194 | NA | NA | NA | NA | NA | N/S | N/S | organic |
| 372.312 | 172 | C14:0 acylcarnitine[1] | NA | NA | NA | 0.18 | major increase | major increase | organic |
| 386.291 | 157 | C14:1 acylcarnitine[1] | NA | NA | NA | -0.76 | major increase | N/S | organic |
| 388.307 | 160 | C14:0-OH acylcarnitine[1] | NA | NA | NA | 0.39 | major increase | N/S | organic |
| 396.312 | 169 | C16:2 acylcarnitine[1] | NA | NA | NA | 0.17 | major increase | N/S | organic |
| 398.328 | 179 | 9-Hexadecenoylcarnitine (Hexadecenoyl-L-carnitine)[1] | NA | NA | NA | 1.05 | major increase | N/S | organic |
| 412.307 | 160 | C16:2-OH acylcarnitine[1] | NA | NA | NA | 0.37 | major increase | N/S | organic |
| 414.322 | 164 | C16:1-OH acylcarnitine[1] | NA | NA | NA | -1.20 | major increase | N/S | organic |
| 416.338 | 171 | C16:0-OH acylcarnitine[1] | NA | NA | NA | -0.35 | major increase | N/S | organic |
| 424.343 | 184 | C18:2 acylcarnitine[1] | NA | NA | NA | -0.55 | major increase | N/S | organic |
| 426.358 | 229 | Spectral Match to Oleoyl-L-carnitine | 0.89 | 12 | 0 | 9 | major increase | N/S | organic |
| 426.359 | 193 | Spectral Match to Oleoyl-L-carnitine | 0.89 | 12 | 0 | 9 | major increase | N/S | organic |
| 428.373 | 240 | C18:0 acylcarnitine[1] | NA | NA | NA | -3.58 | major increase | major increase | organic |
| 428.374 | 205 | C18:0 acylcarnitine[1] | NA | NA | NA | -1.24 | major increase | N/S | organic |

| | | | | | | | | | |
|---|---|---|---|---|---|---|---|---|---|
| 429.378 | 206 | NA | NA | NA | NA | NA | minor increase | minor increase | organic |
| 430.387 | 202 | NA | NA | NA | NA | NA | N/S | N/S | organic |
| 438.323 | 163 | C18:3-OH acylcarnitine[1] | NA | NA | NA | 1.15 | major increase | N/S | organic |
| 440.338 | 168 | C18:2-OH acylcarnitine[1] | NA | NA | NA | -0.33 | major increase | N/S | organic |
| 442.354 | 175 | C18:1-OH acylcarnitine[1] | NA | NA | NA | 0.46 | major increase | N/S | organic |
| 444.369 | 185 | C18:0-OH acylcarnitine[1] | NA | NA | NA | -1.01 | major increase | N/S | organic |
| 450.359 | 190 | C20:4 acylcarnitine[1] | NA | NA | NA | 0.26 | major increase | N/S | organic |
| 452.373 | 244 | C20:2 acylcarnitine[1] | NA | NA | NA | -3.39 | major increase | N/S | organic |
| 452.374 | 199 | C20:2 acylcarnitine[1] | NA | NA | NA | -1.18 | major increase | N/S | organic |
| 454.389 | 250 | C20:1 acylcarnitine[1] | NA | NA | NA | 2.60 | major increase | N/S | organic |
| 454.390 | 210 | C20:1 acylcarnitine[1] | NA | NA | NA | -0.40 | major increase | N/S | organic |
| 455.394 | 209 | NA | NA | NA | NA | NA | major increase | N/S | organic |
| 466.294 | 171 | NA | NA | NA | NA | NA | N/S | N/S | organic |
| 468.369 | 179 | C20:2-OH acylcarnitine[1] | NA | NA | NA | -0.95 | major increase | N/S | organic |
| 470.385 | 189 | C20:1-OH acylcarnitine[1] | NA | NA | NA | -0.21 | major increase | N/S | organic |
| 472.401 | 202 | C20:1-OH acylcarnitine[1] | NA | NA | NA | 0.54 | major increase | N/S | organic |
| 480.406 | 214 | C22:2 acylcarnitine[1] | NA | NA | NA | 0.35 | major increase | N/S | organic |
| 482.420 | 231 | C22:1 acylcarnitine[1] | NA | NA | NA | -3.07 | major increase | N/S | organic |
| 484.437 | 254 | PC family member[1] | NA | NA | NA | NA | major increase | N/S | organic |
| 494.325 | 170 | LPC (16:1)1 | NA | NA | NA | -0.43 | N/S | N/S | organic |
| 510.453 | 249 | C24:1 acylcarnitine[1] | NA | NA | NA | 0.43 | major increase | N/S | organic |
| 512.468 | 276 | C24:0 acylcarnitine[1] | NA | NA | NA | -0.84 | major increase | N/S | organic |
| 513.336 | 207 | NA | NA | NA | NA | NA | major increase | N/S | organic |
| 515.352 | 221 | NA | NA | NA | NA | NA | minor increase | minor increase | organic |
| 541.368 | 228 | NA | NA | NA | NA | NA | major increase | N/S | organic |
| 550.030 | 144 | NA | NA | NA | NA | NA | N/S | N/S | organic |
| 569.398 | 249 | NA | NA | NA | NA | NA | major increase | N/S | organic |
| 570.458 | 211 | NA | NA | NA | NA | NA | N/S | N/S | organic |
| 580.362 | 187 | PC family member[1] | NA | NA | NA | NA | minor decrease | N/S | organic |
| 588.258 | 145 | NA | NA | NA | NA | NA | minor decrease | N/S | organic |
| 596.342 | 177 | NA | NA | NA | NA | NA | minor decrease | N/S | organic |
| 632.392 | 195 | NA | NA | NA | NA | NA | minor increase | N/S | organic |

| | | | | | | | | | |
|---|---|---|---|---|---|---|---|---|---|
| 664.450 | 223 | NA | NA | NA | NA | NA | N/S | N/S | organic |
| 664.528 | 309 | NA | NA | NA | NA | NA | N/S | N/S | organic |
| 677.458 | 214 | NA | NA | NA | NA | NA | N/S | N/S | organic |
| 703.576 | 304 | NA | NA | NA | NA | NA | N/S | N/S | organic |
| 716.449 | 209 | NA | NA | NA | NA | NA | minor decrease | N/S | organic |
| 716.523 | 395 | 1-Palmitoyl-2-oleoyl-sn-glycero-3-phosphoethanolamine | 0.79 | 9 | 1.01 | -0.81 | minor increase | N/S | organic |
| 744.587 | 431 | NA | NA | NA | NA | NA | minor decrease | N/S | organic |
| 800.617 | 452 | NA | NA | NA | NA | NA | minor increase | minor decrease | organic |
| 806.570 | 435 | NA | NA | NA | NA | NA | minor decrease | N/S | organic |
| 807.574 | 354 | NA | NA | NA | NA | NA | minor decrease | N/S | organic |
| 816.590 | 344 | NA | NA | NA | NA | NA | major decrease | N/S | organic |
| 817.577 | 275 | NA | NA | NA | NA | NA | minor increase | N/S | organic |
| 819.594 | 297 | PC family member[1] | NA | NA | NA | NA | minor increase | N/S | organic |
| 868.533 | 230 | NA | NA | NA | NA | NA | minor decrease | N/S | organic |
| 886.540 | 215 | NA | NA | NA | NA | NA | minor decrease | N/S | organic |

[1]Annotated based on molecular networking to an annotated sub-network; NA: not applicable; N/S: not statistically significant

**S6 Table. Annotated metabolites of combined extracts identified as perturbed by infection at all positions (FDR-corrected Mann Whitney p<0.05).**

| **Top-ranking metabolites for CL *T.cruzi* strain** | | | | | | | | | |
|---|---|---|---|---|---|---|---|---|---|
| ***m/z*** | **RT (sec)** | **Annotation** | **Cosine Score** | **No. of Shared Peaks** | **Mass Diff. to Library Reference** | **ppm error** | **Impact of Infection (Sylvio X10/4 vs uninfected 147 days)** | **Impact of Infection (CL vs uninfected 90 days)** | **Extract** |
| 155.071 | 123 | NA | NA | NA | NA | NA | N/S | minor increase | aqueous |
| 165.091 | 170 | NA | NA | NA | NA | NA | N/S | minor increase | aqueous |
| 184.074 | 25 | phosphocholine[1] | NA | NA | NA | -2.27 | N/S | minor increase | aqueous |
| 185.596 | 143 | NA | NA | NA | NA | NA | N/S | major decrease | aqueous |
| 189.124 | 74 | NA | NA | NA | NA | NA | N/S | minor decrease | aqueous |
| 211.145 | 151 | NA | NA | NA | NA | NA | N/S | major decrease | aqueous |
| 218.140 | 39 | NA | NA | NA | NA | NA | N/S | minor decrease | aqueous |
| 251.103 | 147 | NA | NA | NA | NA | NA | N/S | minor decrease | aqueous |
| 252.233 | 190 | NA | NA | NA | NA | NA | N/S | major decrease | aqueous |
| 270.244 | 200 | NA | NA | NA | NA | NA | N/S | major decrease | aqueous |
| 278.248 | 179 | NA | NA | NA | NA | NA | N/S | major decrease | aqueous |
| 280.264 | 206 | NA | NA | NA | NA | NA | N/S | major decrease | aqueous |
| 292.227 | 199 | NA | NA | NA | NA | NA | N/S | major decrease | aqueous |
| 296.259 | 187 | NA | NA | NA | NA | NA | N/S | major decrease | aqueous |
| 312.254 | 194 | NA | NA | NA | NA | NA | N/S | major decrease | aqueous |
| 316.285 | 191 | NA | NA | NA | NA | NA | N/S | major decrease | aqueous |
| 332.280 | 175 | NA | NA | NA | NA | NA | N/S | major decrease | aqueous |
| 336.251 | 190 | NA | NA | NA | NA | NA | N/S | major decrease | aqueous |
| 344.228 | 143 | NA | NA | NA | NA | NA | N/S | minor decrease | aqueous |
| 365.232 | 194 | NA | NA | NA | NA | NA | N/S | major increase | aqueous |
| 368.280 | 177 | NA | NA | NA | NA | NA | N/S | major increase | aqueous |
| 370.178 | 142 | NA | NA | NA | NA | NA | N/S | minor decrease | aqueous |
| 372.312 | 197 | C14:0 acylcarnitine[1] | NA | NA | NA | 0.18 | N/S | major increase | aqueous |
| 379.210 | 214 | NA | NA | NA | NA | NA | N/S | minor increase | aqueous |

| | | | | | | | | | |
|---|---|---|---|---|---|---|---|---|---|
| 380.113 | 41 | NA | NA | NA | NA | NA | N/S | major decrease | aqueous |
| 382.296 | 182 | C15:2 acylcarnitine[1] | NA | NA | NA | -0.74 | N/S | minor increase | aqueous |
| 384.275 | 171 | C14:2-OH acylcarnitine[1] | NA | NA | NA | -1.42 | N/S | minor increase | aqueous |
| 388.255 | 144 | NA | NA | NA | NA | NA | N/S | minor decrease | aqueous |
| 396.312 | 192 | C16:2 acylcarnitine[1] | NA | NA | NA | 0.17 | N/S | major increase | aqueous |
| 400.269 | 187 | C14:2-DC acylcarnitine[1] | NA | NA | NA | -3.65 | N/S | minor increase | aqueous |
| 415.254 | 146 | NA | NA | NA | NA | NA | N/S | minor decrease | aqueous |
| 422.159 | 119 | NA | NA | NA | NA | NA | N/S | major decrease | aqueous |
| 423.237 | 174 | NA | NA | NA | NA | NA | N/S | minor increase | aqueous |
| 432.281 | 145 | NA | NA | NA | NA | NA | N/S | minor decrease | aqueous |
| 446.196 | 147 | NA | NA | NA | NA | NA | N/S | minor decrease | aqueous |
| 465.210 | 148 | NA | NA | NA | NA | NA | N/S | minor decrease | aqueous |
| 476.307 | 146 | NA | NA | NA | NA | NA | N/S | minor decrease | aqueous |
| 478.185 | 141 | NA | NA | NA | NA | NA | N/S | minor decrease | aqueous |
| 482.325 | 234 | NA | NA | NA | NA | NA | N/S | minor increase | aqueous |
| 520.231 | 148 | NA | NA | NA | NA | NA | N/S | minor decrease | aqueous |
| 524.372 | 222 | LPC (18:0)[1] | NA | NA | NA | -0.31 | N/S | minor increase | aqueous |
| 537.340 | 219 | NA | NA | NA | NA | NA | N/S | minor increase | aqueous |
| 605.357 | 169 | NA | NA | NA | NA | NA | N/S | major decrease | aqueous |
| 616.177 | 186 | NA | NA | NA | NA | NA | N/S | minor decrease | aqueous |
| 634.187 | 180 | NA | NA | NA | NA | NA | N/S | minor decrease | aqueous |
| 657.204 | 180 | NA | NA | NA | NA | NA | N/S | minor decrease | aqueous |
| 701.561 | 274 | NA | NA | NA | NA | NA | N/S | minor increase | aqueous |
| 703.575 | 286 | NA | NA | NA | NA | NA | N/S | major increase | aqueous |
| 850.632 | 417 | NA | NA | NA | NA | NA | N/S | minor decrease | organic |
| 931.451 | 189 | NA | NA | NA | NA | NA | N/S | major decrease | aqueous |
| **Top-ranking metabolites for CL and Sylvio X10/4 *T.cruzi* strains** | | | | | | | | | |
| ***m/z*** | **RT (sec)** | **Annotation** | **Cosine Score** | **No. of Shared Peaks** | **Mass Diff. to Library Reference** | **ppm error** | **Impact of Infection (Sylvio X10/4 vs uninfected 147 days)** | **Impact of Infection (CL vs uninfected 90 days)** | **Extract** |
| 193.610 | 145 | NA | NA | NA | NA | NA | major increase | major decrease | aqueous |

| | | | | | | | | | |
|---|---|---|---|---|---|---|---|---|---|
| 209.093 | 57 | NA | NA | NA | NA | NA | major decrease | major increase | organic |
| 209.093 | 53 | NA | NA | NA | NA | NA | major decrease | major increase | aqueous |
| 250.095 | 66 | NA | NA | NA | NA | NA | major increase | minor decrease | aqueous |
| 320.256 | 206 | NA | NA | NA | NA | NA | major increase | major decrease | aqueous |
| 320.256 | 217 | NA | NA | NA | NA | NA | major increase | major decrease | aqueous |
| 368.201 | 147 | NA | NA | NA | NA | NA | major increase | minor decrease | aqueous |
| 375.216 | 186 | NA | NA | NA | NA | NA | minor decrease | minor increase | aqueous |
| 381.227 | 178 | NA | NA | NA | NA | NA | minor decrease | minor increase | aqueous |
| 383.242 | 187 | NA | NA | NA | NA | NA | minor decrease | minor increase | aqueous |
| 389.232 | 202 | NA | NA | NA | NA | NA | minor decrease | minor increase | aqueous |
| 400.343 | 205 | Spectral Match to Palmitoylcarnitine | 0.91 | 11 | 0 | 2 | major increase | major increase | aqueous |
| 405.226 | 186 | NA | NA | NA | NA | NA | minor decrease | minor increase | aqueous |
| 407.242 | 190 | NA | NA | NA | NA | NA | minor decrease | minor increase | aqueous |
| 421.221 | 177 | NA | NA | NA | NA | NA | minor decrease | minor increase | aqueous |
| 426.359 | 210 | Spectral Match to Oleoyl-L-carnitine | 0.85 | 10 | 0 | 0 | major increase | major increase | aqueous |
| 462.191 | 140 | NA | NA | NA | NA | NA | minor increase | minor decrease | aqueous |
| 464.206 | 147 | NA | NA | NA | NA | NA | minor increase | minor decrease | aqueous |
| **Top-ranking metabolites for Sylvio X10/4 *T.cruzi* strain** | | | | | | | | | |
| ***m/z*** | **RT (sec)** | **Annotation** | **Cosine Score** | **No. of Shared Peaks** | **Mass Diff. to Library Reference** | **ppm error** | **Impact of Infection (Sylvio X10/4 vs uninfected 147 days)** | **Impact of Infection (CL vs uninfected 90 days)** | **Extract** |
| 137.047 | 30 | NA | NA | NA | NA | NA | minor decrease | N/S | organic |
| 138.050 | 29 | NA | NA | NA | NA | NA | minor decrease | N/S | organic |
| 147.114 | 14 | NA | NA | NA | NA | NA | minor decrease | N/S | aqueous |
| 148.061 | 27 | NA | NA | NA | NA | NA | minor increase | N/S | aqueous |
| 150.059 | 27 | NA | NA | NA | NA | NA | minor decrease | N/S | aqueous |
| 172.985 | 26 | NA | NA | NA | NA | NA | minor increase | N/S | aqueous |
| 216.197 | 150 | NA | NA | NA | NA | NA | minor decrease | N/S | organic |
| 220.136 | 160 | NA | NA | NA | NA | NA | major increase | N/S | aqueous |
| 223.133 | 183 | NA | NA | NA | NA | NA | minor increase | N/S | aqueous |

| | | | | | | | | | |
|---|---|---|---|---|---|---|---|---|---|
| 227.091 | 144 | NA | NA | NA | NA | NA | minor decrease | N/S | aqueous |
| 232.155 | 115 | C4:0 acylcarnitine (butyrylcarnitine)[1] | NA | NA | NA | -1.86 | minor increase | N/S | aqueous |
| 240.102 | 69 | NA | NA | NA | NA | NA | minor increase | N/S | aqueous |
| 244.155 | 140 | NA | NA | NA | NA | NA | minor increase | N/S | aqueous |
| 246.171 | 142 | C5:0 acylcarnitine (valerylcarnitine)[1] | NA | NA | NA | -0.33 | major increase | N/S | aqueous |
| 248.150 | 36 | C4:0-OH acylcarnitine[1] | NA | NA | NA | -1.39 | major increase | N/S | organic |
| 248.150 | 30 | C4:0-OH acylcarnitine[1] | NA | NA | NA | -1.39 | major increase | N/S | aqueous |
| 248.151 | 19 | C4:0-OH acylcarnitine[1] | NA | NA | NA | 2.64 | major increase | N/S | organic |
| 248.151 | 47 | NA | NA | NA | NA | NA | major increase | N/S | aqueous |
| 257.117 | 31 | NA | NA | NA | NA | NA | minor increase | N/S | aqueous |
| 258.171 | 145 | C6:1 acylcarnitine[1] | NA | NA | NA | -0.32 | major increase | N/S | aqueous |
| 260.187 | 141 | C6:0 acylcarnitine[1] | NA | NA | NA | 1.03 | minor increase | N/S | organic |
| 260.187 | 148 | C6:0 acylcarnitine[1] | NA | NA | NA | 1.03 | major increase | N/S | aqueous |
| 262.166 | 43 | CAR (5:0(OH))[1] | NA | NA | NA | 0.02 | minor increase | N/S | aqueous |
| 269.093 | 43 | NA | NA | NA | NA | NA | major decrease | N/S | organic |
| 274.183 | 153 | NA | NA | NA | NA | NA | major increase | N/S | organic |
| 274.184 | 166 | NA | NA | NA | NA | NA | major increase | N/S | aqueous |
| 276.181 | 104 | NA | NA | NA | NA | NA | major increase | N/S | aqueous |
| 276.181 | 139 | NA | NA | NA | NA | NA | major increase | N/S | aqueous |
| 286.202 | 154 | C8:1 acylcarnitine[1] | NA | NA | NA | -1.33 | major increase | N/S | aqueous |
| 288.218 | 159 | C8:0 acylcarnitine[1] | NA | NA | NA | -0.11 | major increase | N/S | aqueous |
| 299.201 | 171 | NA | NA | NA | NA | NA | minor decrease | N/S | aqueous |
| 302.197 | 145 | NA | NA | NA | NA | NA | major increase | N/S | aqueous |
| 302.207 | 157 | NA | NA | NA | NA | NA | major increase | N/S | organic |
| 302.215 | 161 | NA | NA | NA | NA | NA | major increase | N/S | organic |
| 302.216 | 178 | NA | NA | NA | NA | NA | major increase | N/S | aqueous |
| 304.213 | 141 | CAR (8:1(OH))[1] | NA | NA | NA | 0.17 | major increase | N/S | organic |
| 304.213 | 149 | CAR (8:1(OH))[1] | NA | NA | NA | 0.17 | major increase | N/S | aqueous |
| 316.249 | 170 | C10:0 acylcarnitine[1] | NA | NA | NA | -1.05 | major increase | N/S | aqueous |
| 326.379 | 206 | NA | NA | NA | NA | NA | major increase | N/S | organic |
| 326.379 | 231 | NA | NA | NA | NA | NA | major increase | N/S | organic |

| | | | | | | | | | |
|---|---|---|---|---|---|---|---|---|---|
| 332.244 | 146 | C10:0-OH acylcarnitine[1] | NA | NA | NA | -0.74 | major increase | N/S | organic |
| 332.244 | 158 | C10:0-OH acylcarnitine[1] | NA | NA | NA | -0.74 | major increase | N/S | aqueous |
| 358.260 | 150 | NA | NA | NA | NA | NA | major increase | N/S | organic |
| 360.275 | 153 | C12:0-OH acylcarnitine[1] | NA | NA | NA | -1.52 | major increase | N/S | organic |
| 360.275 | 170 | C12:0-OH acylcarnitine[1] | NA | NA | NA | -1.52 | major increase | N/S | aqueous |
| 368.280 | 158 | C14:2 acylcarnitine[1] | NA | NA | NA | -1.72 | major increase | N/S | organic |
| 370.296 | 166 | C14:1 acylcarnitine[1] | NA | NA | NA | -0.76 | major increase | N/S | organic |
| 372.312 | 172 | C14:0 acylcarnitine[1] | NA | NA | NA | 0.18 | major increase | N/S | organic |
| 384.116 | 136 | NA | NA | NA | NA | NA | major increase | N/S | aqueous |
| 384.179 | 139 | NA | NA | NA | NA | NA | minor increase | N/S | aqueous |
| 386.211 | 146 | NA | NA | NA | NA | NA | major increase | N/S | aqueous |
| 386.291 | 157 | C14:1 acylcarnitine[1] | NA | NA | NA | -0.76 | major increase | N/S | organic |
| 386.291 | 175 | NA | NA | NA | NA | NA | major increase | N/S | aqueous |
| 388.307 | 160 | C14:0-OH acylcarnitine[1] | NA | NA | NA | 0.39 | major increase | N/S | organic |
| 388.307 | 181 | C14:0-OH acylcarnitine[1] | NA | NA | NA | 0.39 | major increase | N/S | aqueous |
| 390.206 | 149 | NA | NA | NA | NA | NA | minor increase | N/S | aqueous |
| 396.312 | 169 | C16:2 acylcarnitine[1] | NA | NA | NA | 0.17 | major increase | N/S | organic |
| 398.195 | 143 | NA | NA | NA | NA | NA | minor increase | N/S | aqueous |
| 398.327 | 198 | 9-Hexadecenoylcarnitine (Hexadecenoyl-L-carnitine)[1] | NA | NA | NA | -1.46 | major increase | N/S | aqueous |
| 398.328 | 179 | 9-Hexadecenoylcarnitine (Hexadecenoyl-L-carnitine)[1] | NA | NA | NA | 1.05 | major increase | N/S | organic |
| 400.343 | 187 | Spectral Match to palmitoylcarnitine | 0.91 | 12 | 0 | 6.74 | major increase | N/S | organic |
| 400.343 | 209 | Spectral Match to palmitoylcarnitine | 0.91 | 12 | 0 | 6.74 | major increase | N/S | organic |
| 412.190 | 137 | NA | NA | NA | NA | NA | minor increase | N/S | aqueous |
| 412.200 | 132 | NA | NA | NA | NA | NA | minor increase | N/S | aqueous |
| 412.210 | 147 | NA | NA | NA | NA | NA | minor increase | N/S | aqueous |
| 412.307 | 160 | C16:2-OH acylcarnitine[1] | NA | NA | NA | 0.37 | major increase | N/S | organic |
| 412.307 | 179 | C16:2-OH acylcarnitine[1] | NA | NA | NA | 0.37 | major increase | N/S | aqueous |
| 414.322 | 164 | C16:1-OH acylcarnitine[1] | NA | NA | NA | -1.20 | major increase | N/S | organic |
| 414.322 | 186 | C16:1-OH acylcarnitine[1] | NA | NA | NA | -1.20 | major increase | N/S | aqueous |

| | | | | | | | | | |
|---|---|---|---|---|---|---|---|---|---|
| 414.359 | 193 | C17:0 acylcarnitine[1] | NA | NA | NA | 0.28 | major increase | N/S | organic |
| 416.144 | 34 | NA | NA | NA | NA | NA | minor increase | N/S | aqueous |
| 416.338 | 171 | C16:0-OH acylcarnitine[1] | NA | NA | NA | -0.35 | major increase | N/S | organic |
| 416.338 | 192 | C16:0-OH acylcarnitine[1] | NA | NA | NA | -0.35 | major increase | N/S | aqueous |
| 420.144 | 139 | NA | NA | NA | NA | NA | minor increase | N/S | aqueous |
| 424.269 | 185 | NA | NA | NA | NA | NA | minor decrease | N/S | aqueous |
| 424.343 | 184 | C18:2 acylcarnitine[1] | NA | NA | NA | -0.55 | major increase | N/S | organic |
| 424.343 | 203 | C18:2 acylcarnitine[1] | NA | NA | NA | -0.55 | major increase | N/S | aqueous |
| 426.358 | 229 | Spectral Match to Oleoyl-L-carnitine | 0.89 | 12 | 0 | 9 | major increase | N/S | organic |
| 426.359 | 193 | Spectral Match to Oleoyl-L-carnitine | 0.89 | 12 | 0 | 9 | major increase | N/S | organic |
| 428.373 | 240 | C18:0 acylcarnitine[1] | NA | NA | NA | -3.58 | major increase | N/S | organic |
| 428.374 | 205 | C18:0 acylcarnitine[1] | NA | NA | NA | -1.24 | major increase | N/S | organic |
| 428.374 | 217 | C18:0 acylcarnitine[1] | NA | NA | NA | -1.24 | major increase | N/S | aqueous |
| 436.211 | 150 | NA | NA | NA | NA | NA | minor increase | N/S | aqueous |
| 438.323 | 163 | C18:3-OH acylcarnitine[1] | NA | NA | NA | 1.15 | major increase | N/S | organic |
| 440.214 | 141 | NA | NA | NA | NA | NA | minor increase | N/S | aqueous |
| 440.338 | 168 | C18:2-OH acylcarnitine[1] | NA | NA | NA | -0.33 | major increase | N/S | organic |
| 440.338 | 190 | C18:2-OH acylcarnitine[1] | NA | NA | NA | -0.33 | major increase | N/S | aqueous |
| 442.353 | 197 | C18:1-OH acylcarnitine[1] | NA | NA | NA | -1.80 | major increase | N/S | aqueous |
| 442.354 | 175 | C18:1-OH acylcarnitine[1] | NA | NA | NA | 0.46 | major increase | N/S | organic |
| 442.391 | 218 | C19:0 acylcarnitine[1] | NA | NA | NA | 1.85 | major increase | N/S | organic |
| 444.369 | 185 | C18:0-OH acylcarnitine[1] | NA | NA | NA | -1.01 | major increase | N/S | organic |
| 444.369 | 204 | C18:0-OH acylcarnitine[1] | NA | NA | NA | -1.01 | major increase | N/S | aqueous |
| 450.359 | 190 | C20:4 acylcarnitine[1] | NA | NA | NA | 0.26 | major increase | N/S | organic |
| 452.373 | 244 | C20:2 acylcarnitine[1] | NA | NA | NA | -3.39 | major increase | N/S | organic |
| 452.374 | 199 | C20:2 acylcarnitine[1] | NA | NA | NA | -1.18 | major increase | N/S | organic |
| 454.389 | 250 | C20:1 acylcarnitine[1] | NA | NA | NA | -2.60 | major increase | N/S | organic |
| 454.390 | 210 | C20:1 acylcarnitine[1] | NA | NA | NA | -0.40 | major increase | N/S | organic |
| 455.394 | 209 | NA | NA | NA | NA | NA | major increase | N/S | organic |
| 456.406 | 229 | C20:0 acylcarnitine[1] | NA | NA | NA | 0.37 | major increase | N/S | organic |
| 462.191 | 148 | NA | NA | NA | NA | NA | minor increase | N/S | aqueous |

| | | | | | | | | | |
|---|---|---|---|---|---|---|---|---|---|
| 468.368 | 198 | C20:2-OH acylcarnitine[1] | NA | NA | NA | -3.09 | major increase | N/S | aqueous |
| 468.369 | 179 | C20:2-OH acylcarnitine[1] | NA | NA | NA | -0.95 | major increase | N/S | organic |
| 470.385 | 189 | C20:2-OH acylcarnitine[1] | NA | NA | NA | -0.21 | major increase | N/S | organic |
| 472.401 | 202 | C20:1-OH acylcarnitine[1] | NA | NA | NA | 0.54 | major increase | N/S | organic |
| 476.170 | 140 | NA | NA | NA | NA | NA | minor increase | N/S | aqueous |
| 478.151 | 139 | NA | NA | NA | NA | NA | minor increase | N/S | aqueous |
| 478.168 | 143 | NA | NA | NA | NA | NA | minor increase | N/S | aqueous |
| 480.201 | 147 | NA | NA | NA | NA | NA | minor increase | N/S | aqueous |
| 480.343 | 212 | LPC (O-16:1)[1] | NA | NA | NA | -6.14 | major decrease | N/S | aqueous |
| 480.406 | 214 | C22:2 acylcarnitine[1] | NA | NA | NA | 0.35 | major increase | N/S | organic |
| 482.420 | 231 | C22:1 acylcarnitine[1] | NA | NA | NA | -3.07 | major increase | N/S | organic |
| 484.364 | 175 | NA | NA | NA | NA | NA | major increase | N/S | organic |
| 484.437 | 254 | NA | NA | NA | NA | NA | major increase | N/S | organic |
| 492.272 | 183 | NA | NA | NA | NA | NA | minor increase | N/S | aqueous |
| 508.376 | 215 | LPC (O-18:1)[1] | NA | NA | NA | -2.46 | major decrease | N/S | aqueous |
| 510.283 | 168 | NA | NA | NA | NA | NA | minor increase | N/S | aqueous |
| 510.453 | 249 | C24:1 acylcarnitine[1] | NA | NA | NA | 0.43 | major increase | N/S | organic |
| 512.468 | 276 | C24:0 acylcarnitine[1] | NA | NA | NA | -0.84 | major increase | N/S | organic |
| 513.336 | 207 | NA | NA | NA | NA | NA | major increase | N/S | organic |
| 528.293 | 157 | NA | NA | NA | NA | NA | major increase | N/S | aqueous |
| 534.319 | 181 | PC family member[1] | NA | NA | NA | NA | minor increase | N/S | aqueous |
| 541.368 | 228 | NA | NA | NA | NA | NA | major increase | N/S | organic |
| 552.330 | 169 | PC family member[1] | NA | NA | NA | NA | major decrease | N/S | aqueous |
| 568.324 | 162 | NA | NA | NA | NA | NA | minor increase | N/S | aqueous |
| 568.325 | 190 | NA | NA | NA | NA | NA | minor increase | N/S | aqueous |
| 569.398 | 249 | NA | NA | NA | NA | NA | major increase | N/S | organic |
| 570.340 | 155 | Lysophosphatidylcholine(22:4)[1] | NA | NA | NA | -28.95 | minor increase | N/S | aqueous |
| 570.341 | 173 | NA | NA | NA | NA | NA | minor increase | N/S | aqueous |
| 580.362 | 187 | PC family member[1] | NA | NA | NA | NA | minor decrease | N/S | organic |
| 584.335 | 182 | NA | NA | NA | NA | NA | minor decrease | N/S | aqueous |
| 594.339 | 163 | NA | NA | NA | NA | NA | minor increase | N/S | aqueous |
| 596.342 | 177 | NA | NA | NA | NA | NA | minor decrease | N/S | organic |

| | | | | | | | | | |
|---|---|---|---|---|---|---|---|---|---|
| 600.330 | 159 | NA | NA | NA | NA | NA | minor decrease | N/S | organic |
| 600.330 | 174 | NA | NA | NA | NA | NA | minor decrease | N/S | aqueous |
| 614.310 | 174 | NA | NA | NA | NA | NA | minor decrease | N/S | aqueous |
| 632.320 | 150 | NA | NA | NA | NA | NA | minor decrease | N/S | organic |
| 716.523 | 395 | 1-Palmitoyl-2-oleoyl-sn-glycero-3-phosphoethanolamine | 0.79 | 9 | 1.01 | -0.81 | minor increase | N/S | organic |
| 744.560 | 406 | NA | NA | NA | NA | NA | minor increase | N/S | organic |
| 800.617 | 416 | NA | NA | NA | NA | NA | minor increase | N/S | organic |
| 800.617 | 452 | NA | NA | NA | NA | NA | minor increase | N/S | organic |
| 816.590 | 344 | NA | NA | NA | NA | NA | major decrease | N/S | organic |
| 817.577 | 275 | NA | NA | NA | NA | NA | minor increase | N/S | organic |
| 819.594 | 297 | PC family member[1] | NA | NA | NA | NA | minor increase | N/S | organic |
| 868.533 | 230 | NA | NA | NA | NA | NA | minor decrease | N/S | organic |

[1]Annotated based on molecular networking to an annotated sub-network; NA: not applicable; N/S: not statistically significant

**S1 Figure. Principal coordinate analysis plot of *T. cruzi* strain CL infected (red) and uninfected (blue) heart tissue samples.** Statistically different clustering found in position C (PERMANOVA p-value<0.05).

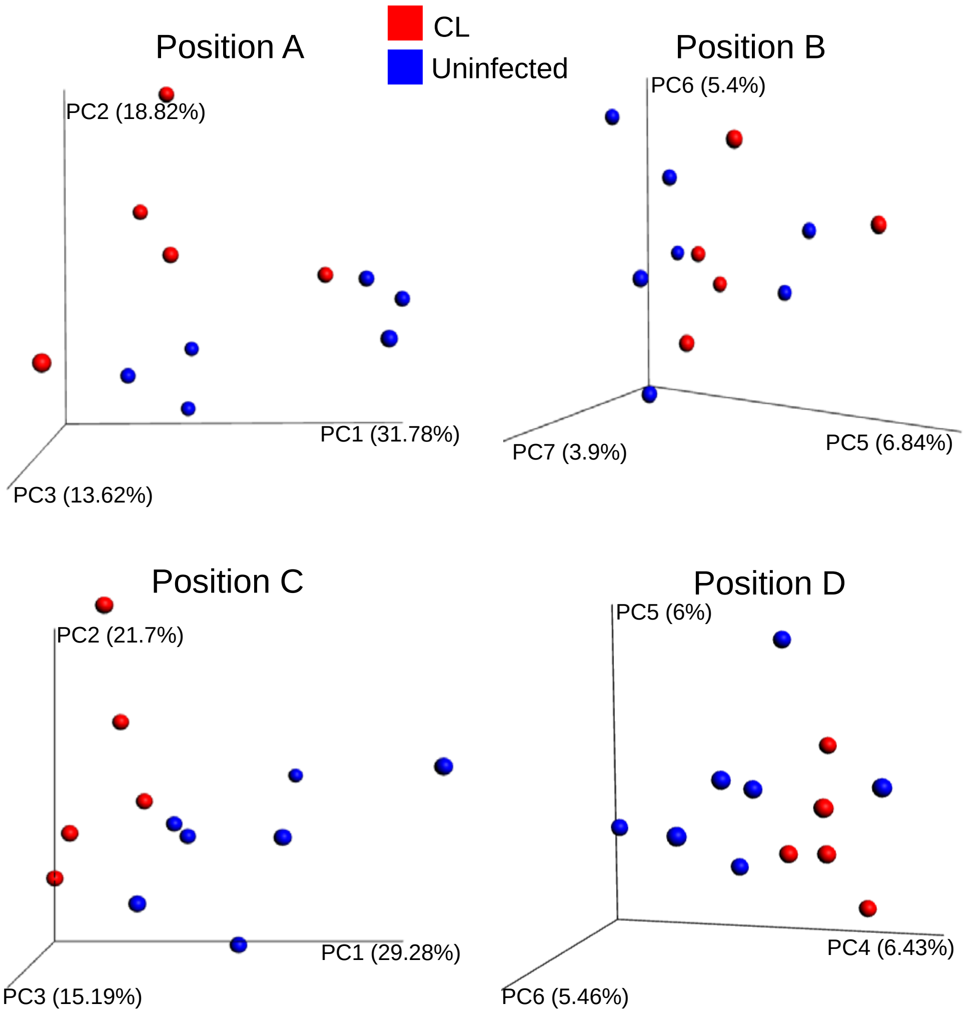

**S2 Figure. Principal coordinate analysis plot of *T. cruzi* strain Sylvio X10/4 infected (gold) and uninfected (blue) heart tissue samples.** Statistically different clustering found in position D (PERMANOVA p-value<0.05).

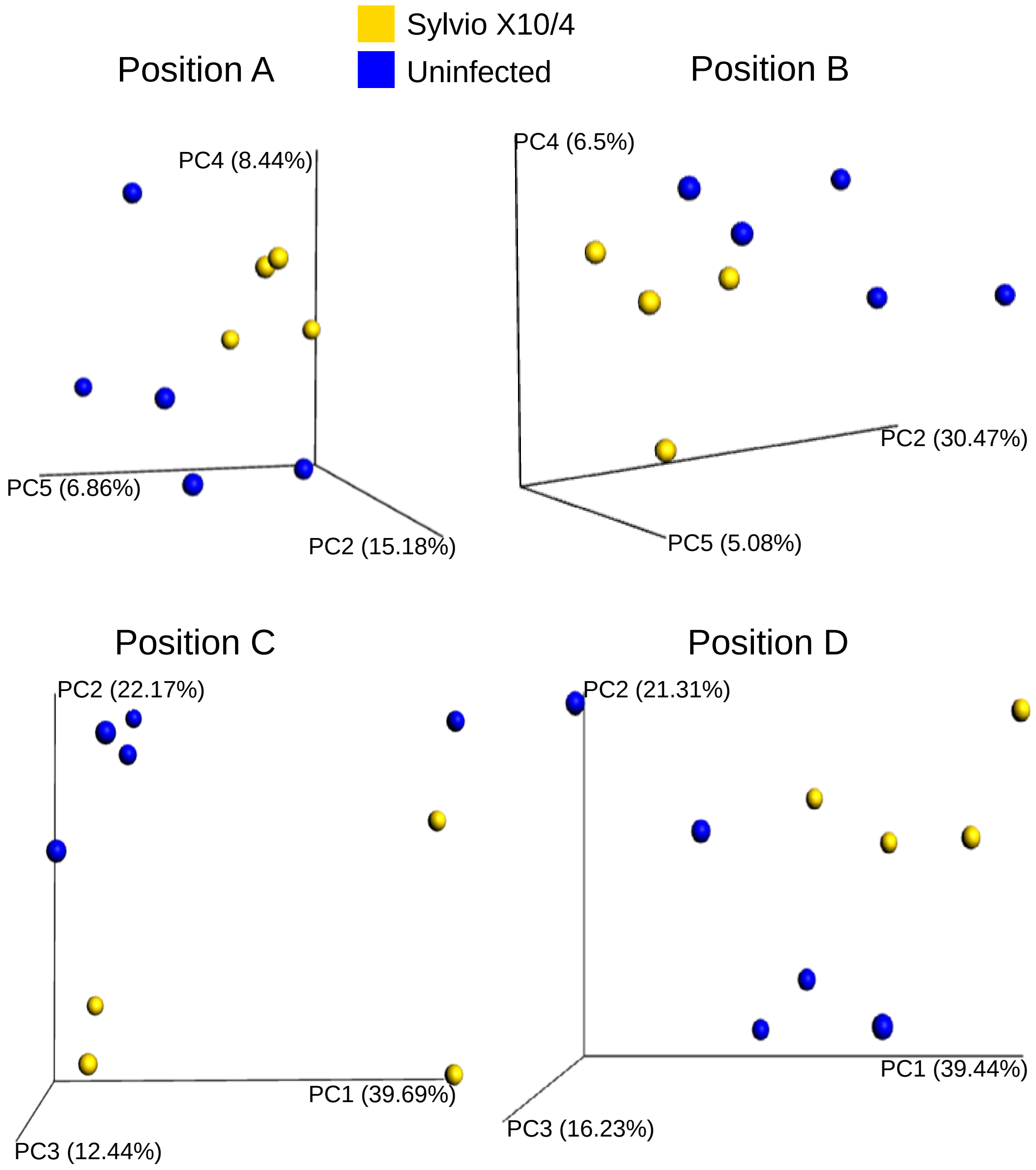

**S3 Figure. Sub-molecular networks and mirror plot of aqueous and organic extract acylcarnitines and phosphocholines.** Each pie chart is one metabolite colored by MS2 spectral count in CL-infected and Sylvio X10/4-infected samples where red is CL and gold is Sylvio X10/4**. (**A) Subnetwork of aqueous extract acylcarnitines with representative acylcarnitine mirror plot (acetylcarnitine, *m/z* -204.124). (B) Subnetwork of aqueous extract phosphocholines with representative phosphocholine mirror plot (Spectral match to 1-Hexadecanoyl-2-(9Z-octadecenoyl)-sn-glycero-3-phosphocholine, *m/z* 758.65). (C) Subnetwork of organic extract acylcarnitines with representative acylcarnitine mirror plot (acetylcarnitine, *m/z*- 204.126 ). (D) Subnetwork of organic extract phosphocholines with representative phosphocholine mirror plot (Spectral Match to 1-Oleoyl-2-palmitoyl-sn-glycero-3-phosphocholine, *m/z* -760.601).

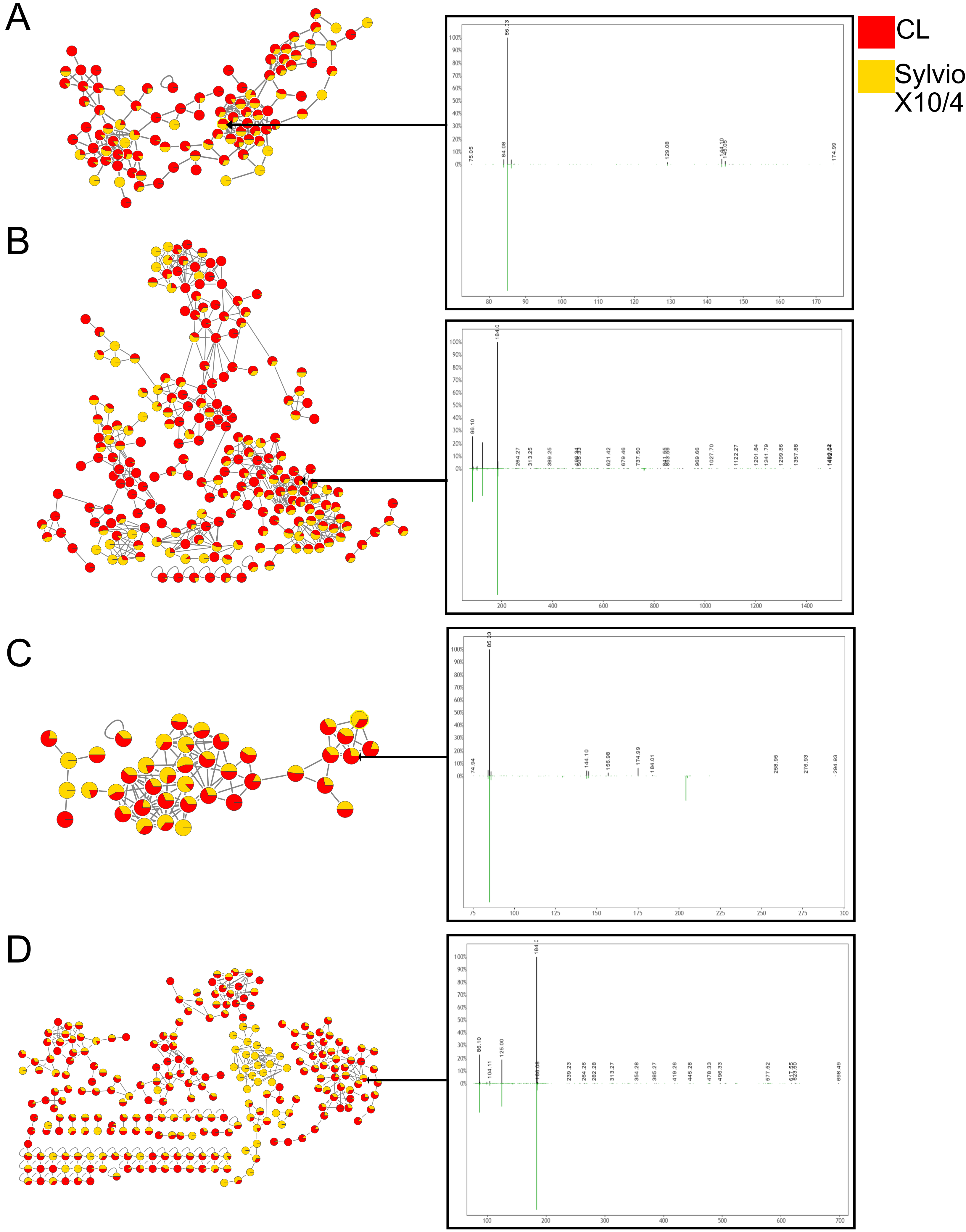
A
B
C
D
CL
Sylvio X10/4
85.03
84.08
75.05
129.08
144.10
145.05
174.99
184.0
86.10
264.27
313.25
389.25
621.42
679.46
737.50
969.66
1027.70
1122.27
1201.84
1241.79
1299.86
1357.88
74.94
144.10
156.98
174.99
184.01
258.95
276.93
294.93
86.10
104.11
125.00
239.23
264.26
282.28
313.27
354.28
385.27
419.26
445.28
478.33
496.33
577.52
698.49

**S4 Figure. GNPS mirror plots of annotated metabolites. (**A) mirror plot of *m/z* 703.575, RT 286s (top, black) to reference library spectrum (SM(d18:1/16:0), bottom, green). (B) mirror plot of *m/z* 454.294, RT 206s (top, black) to reference library spectrum (hexadecanoyl-lysophosphatidylethanolamine, bottom, green). (C) mirror plot of *m/z* 377.146, RT 137s (top, black) to reference library spectrum (riboflavin, bottom, green). (D) mirror plot of *m/z* 646.614, RT 417s (top, black) to reference library spectrum (ceramide, bottom, green). (E) mirror plot of *m/z* 716.523, RT 395s (top, black) to reference library spectrum (1-palmitoyl-2-oleoyl-sn-glycero-3-phosphoethanolamine, bottom, green).

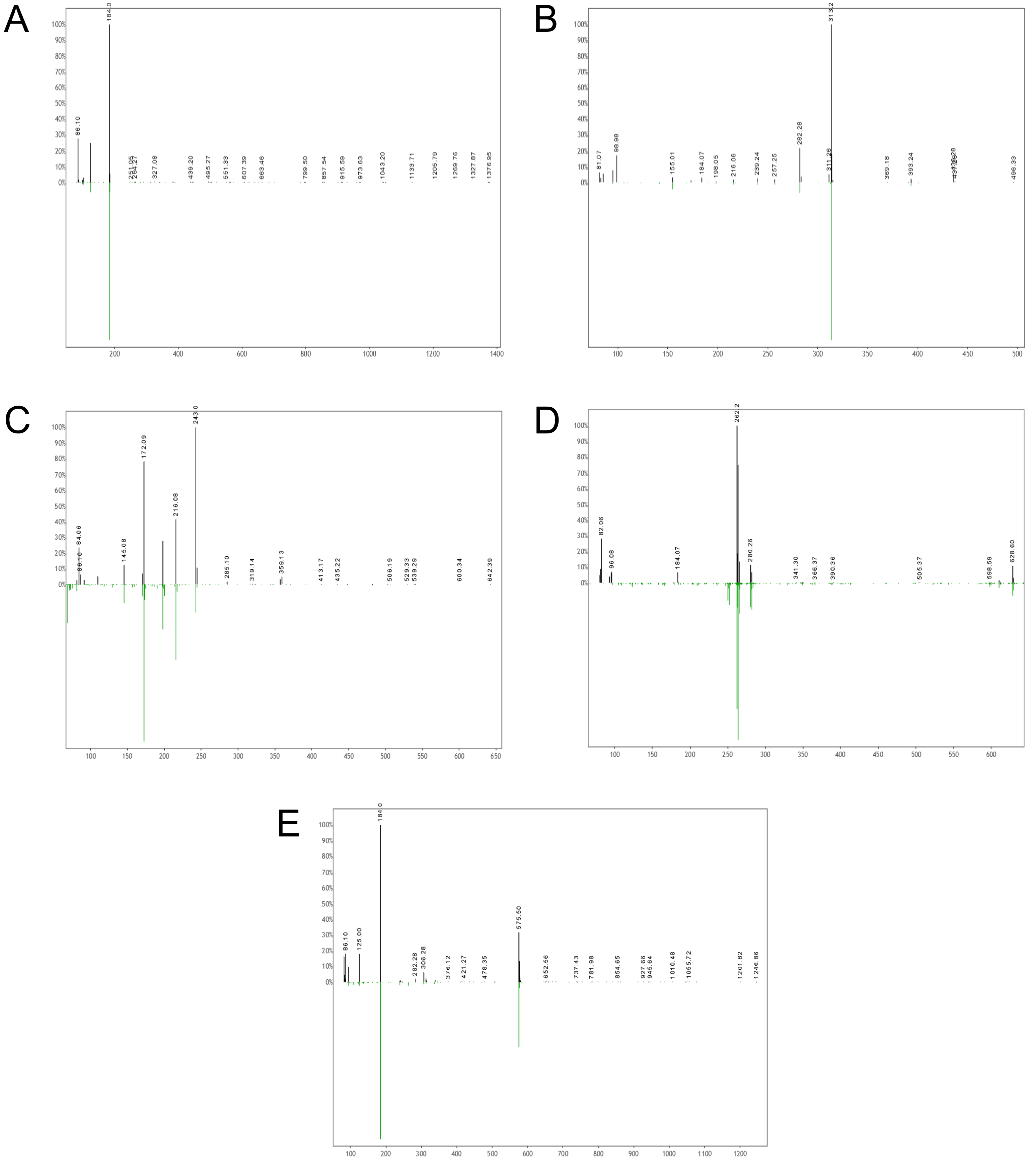
A
184.0
86.10
251.05
264.27
327.08
439.20
495.27
551.33
607.39
663.46
799.50
857.54
915.59
973.63
1043.20
1133.71
1205.79
1269.76
1327.87
1376.95
B
313.2
81.07
98.98
155.01
184.07
198.05
216.06
239.24
257.25
282.28
311.26
369.18
393.24
496.33
C
243.0
172.09
216.08
84.06
145.08
285.10
319.14
359.13
413.17
435.22
506.19
529.33
539.29
600.34
642.39
D
262.2
82.06
96.08
184.07
280.26
341.30
366.37
390.36
505.37
598.59
628.60
E
184.0
575.50
86.10
125.00
282.28
306.28
376.12
421.27
478.35
652.56
737.43
781.98
854.65
927.66
945.64
1010.48
1055.72
1201.82
1246.86
